# Mapping of the hSOX10 protein interactome in human melanoma

**DOI:** 10.1101/2025.09.03.672723

**Authors:** Catherine M. Newsom-Stewart, Dhaval P. Bhatt, Michael B. Major, Charles K. Kaufman

## Abstract

The transcription factor SOX10 is a central regulator of melanoma biology, influencing tumor initiation, progression, phenotypic plasticity, and therapeutic resistance. However, the molecular mechanisms underlying these diverse functions remain incompletely understood. To elucidate the protein-protein interactions (PPIs) that mediate SOX10 activity in melanoma, we mapped the human SOX10 (hSOX10) interactome for the first time in human melanoma cells (A375 line). Given the challenges of capturing transient and weak transcription factor interactions, we employed miniTurbo (mT) proximity-dependent biotinylation coupled with mass spectrometry (MS). Stable A375 cell lines expressing N- and C-terminal mT-tagged hSOX10 fusion proteins at physiologically relevant levels were generated, enabling unbiased biotinylation and MS-based identification of candidate interacting proteins. This approach revealed 847 melanoma-enriched candidate interactors, including 213 high-confidence hits. These included both known hSOX10 partners and previously unidentified putative interactors. Functional annotation of our hSOX10 interactome highlighted associations with chromatin remodeling, transcriptional regulation, SUMOylation, and DNA damage response pathways. Western blot validation of select interactors supported the robustness of our dataset. This first comprehensive map of the hSOX10 interactome in melanoma provides a critical foundation for future investigations into SOX10-driven transcriptional networks and their potential as therapeutic targets and biomarkers in melanoma.

## INTRODUCTION

Melanoma is an aggressive skin cancer originating from melanocytes, the pigment-producing cells of the skin. It poses a major public health challenge due to its high metastatic potential, increasing global incidence, and frequent resistance to even advanced therapies.^1,2^ It accounts for the majority of skin cancer-related deaths and is notorious for its phenotypic heterogeneity, cellular plasticity, and ability to adapt to therapeutic pressures.^1–3^ Despite breakthroughs with targeted therapies, such as BRAF and MEK inhibitors, and immune checkpoint inhibitors, their effectiveness is often curtailed by the emergence of resistance and toxicities, underscoring the need for deeper biological insights and novel therapeutic strategies.^3–5^

Melanoma cells and their melanocyte precursors are derived from the embryonic neural crest (NC), a population of multipotent cells that give rise to diverse lineages during development.^6,7^ Melanoma development and progression are closely linked to NC-derived transcription factors, with SRY-Box Transcription Factor 10 (SOX10) being a key player. SOX10 is essential for melanocyte development and maintenance, and in melanoma, it serves as a major dependency factor.^1,81,4,9^ It governs tumor progression, cellular proliferation, and phenotype plasticity, including transitions between proliferative and invasive states.^10^ These features contribute significantly to melanoma’s aggressiveness and therapeutic resistance.^3,11^ In patients, SOX10 modulates resistance pathways, in part by modifying TGFβ-EGFR signaling and immune checkpoint regulation, highlighting its pivotal role in melanoma pathogenesis.^4,11,12^

Despite its established role as a lineage-defining transcription factor, the precise mechanisms through which SOX10 exerts these diverse effects remain incompletely understood.^13^ SOX10 functions as part of the complex NC gene regulatory network (GRN), interacting with transcription factors, cofactors, and signaling pathways critical to NC development, lineage specification, and melanoma formation and maintenance.^6,8,11,13^ These protein-protein interactions (PPIs) are essential for cellular functions such as transcriptional regulation, chromatin remodeling, and cellular adaptation.^13–16^ Thus, uncovering the factors with which SOX10 interacts, its so-called interactome, is a promising avenue for more fully elucidating its role in these processes.^8,17,18^

Traditional methods for mapping PPIs, such as co-immunoprecipitation, often fail to capture the transient, low-affinity, and context-dependent interactions characteristic of transcription factors like SOX10.^15,16,19–21^ Proximity-dependent labeling approaches, such as miniTurbo (mT) biotinylation, overcome these limitations by enabling the capture of even weak, transient, and dynamic PPIs within native environments.^20–24^

In this study, we leverage mT-based proximity labeling coupled with mass spectrometry (MS) to identify putative members of the human SOX10 (hSOX10) protein interactome in A375 human melanoma cells, and generate a high-resolution map of this physical interaction network.^15,20,22,25^ We employ both N- and C-terminal mT-tagged hSOX10 fusion proteins expressed at physiologic levels in A375 melanoma cells, conduct streptavidin pull-down of biotinylated proteins, and utilize MS to identify candidate hSOX10 interactors. This workflow revealed 847 human melanoma-enriched candidate interactors, with 213 proteins meeting the most stringent analysis criteria. These high-confidence candidate interactors are involved in biological processes such as transcriptional regulation, chromatin remodeling, SUMOylation, and DNA repair. We validated a subset of these interactors using Western blot analysis which supported the quality and utility of this study. Notably, our SOX10 interactome includes multiple proteins from the same complexes, such as the SWI/SNF and MLL3/4 complexes, comprising both established and novel hSOX10 interactors. Many of these proteins are frequently mutated in public human melanoma genome databases (e.g., KMT2C, KMT2D, ARID2, and SMARCA4), further linking hSOX10 to biological processes critical in melanoma, such as DNA damage response pathways, which are frequently dysregulated due to UV-induced mutagenesis, and developmental plasticity, which underlies melanoma’s cellular heterogeneity and adaptive resistance to therapy.

This first comprehensive mapping of the hSOX10 protein interactome in melanoma provides a critical resource to support the identification of novel therapeutic targets and biomarkers, ultimately facilitating more personalized approaches to melanoma management.^3^

## METHODS

### Cell lines

The human melanoma cell line A375 (ATCC, CRL-1619) was used to generate the following three stable lines expressing mT-tagged fusion proteins: N-terminal mT-tagged hSOX10 (mT-hSOX10), C-terminal mT-tagged hSOX10 (hSOX10-mT), and mT with a nuclear localization signal (mT-NLS). A375 cells were cultured in Dulbecco’s Modified Eagle Medium (DMEM, Corning, MT10013CV) supplemented with 10% fetal bovine serum (FBS, Sigma-Aldrich, F2442) and 1% penicillin-streptomycin (Gibco, 15140122). HEK293T/17 cells (ATCC, CRL-11268) were used for lentivirus production and maintained in DMEM with 10% FBS and 1% penicillin-streptomycin, except during transfections, where DMEM with 10% FBS (without penicillin-streptomycin) was used. For stable cell selection, puromycin-containing media consisted of DMEM with 10% FBS, 1% penicillin-streptomycin, and 1 µg/mL puromycin (Fisher Scientific, NC0721599). For biotinylation assays, A375 cells expressing mT-tagged constructs were treated in DMEM with 5% FBS, supplemented with 50µM biotin. All cultures were maintained at 37°C in a humidified chamber containing 5% CO_2_.

### Cloning of hSOX10 constructs and NLS control

Both mT-hSOX10 and hSOX10-mT fusion constructs were generated using the Gateway recombinational cloning system, employing single-site and multisite cloning strategies depending on the construct. The hSOX10 coding sequence was obtained from the pDONR221- hSOX10 plasmid (Addgene, Plasmid #24749). Two versions of this plasmid were utilized: the original plasmid, containing the stop codon, and a modified version with the stop codon removed, which was generated using Q5 site-directed mutagenesis (New England Biolabs). Both versions were recombined into Destination vectors using LR Clonase II Plus enzyme mix (Invitrogen, 12538120).

Gateway cloning was conducted following previously described protocols.^26^ Briefly, the first construct, pLenti663-UBC-mT-hSOX10 (mT-hSOX10), was generated using multisite Gateway cloning to fuse mT to the N-terminus of hSOX10 (with a stop codon). The reaction included 25 ng of mT Entry clone, 25 ng of UBC Entry clone, 40 ng of hSOX10 plasmid DNA with a stop codon, 100 ng of pDest663 Destination vector, and LR Clonase II Plus enzyme mix. The second construct, pLenti667-UBC-hSOX10-mT (hSOX10-mT), was generated using single-site Gateway cloning to fuse hSOX10 (without a stop codon) to the C-terminal mT tag. The reaction included 50 ng of hSOX10 plasmid DNA without a stop codon, 100 ng of pLenti667- UBC-GW-mT Destination vector, and LR Clonase II Plus enzyme mix. For both constructs, the reactions were incubated and transformed into *E. coli* Stbl3™ chemically competent cells (Invitrogen, C737303) for amplification.

All resulting plasmids were validated by restriction digestion, Sanger sequencing, and whole-plasmid sequencing by Plasmidsaurus. Sanger sequencing primers included M13R, in-house primer (TGAAGCTCCGGTTTTGAACT) for the UBC promoter region in both pLenti663 and pLenti667 constructs, in-house primer (CATTTCCTCCCTCTGCTTCC) to confirm the N-terminal mT fusion in pLenti667, and in-house primer (AATAGACCCGTGAAGCTGATC) to confirm the C-terminal mT fusion in pLenti663. An NLS control construct (mT-NLS) was previously generated by and obtained from the Major lab at Washington University School of Medicine in St. Louis, MO.

### Lentiviral production

For lentiviral production, HEK293T/17 cells were used. Lentivirus was produced using a calcium phosphate transfection method. Specifically, 4.4 x 10^6^ HEK293T/17 cells were plated in 10-cm dishes in DMEM with 10% FBS and cultured for 24 hours prior to transfection. On the day of transfection, 8 µg of transfer plasmid—mT-hSOX10, hSOX10-mT, or mT-NLS—along with 6 µg of psPAX2 (packaging plasmid; Addgene, Plasmid #12260) and 2 µg of pMD2.G (envelope plasmid; Addgene, Plasmid #12259), were combined with 2.5 M CaCl2 (Sigma Aldrich, C4901) and mixed with 2X HEPES-buffered saline (HBS; 280 mM NaCl, 1.5 mM Na_2_HPO_4_•H_2_O, 50 mM HEPES) (pH 7.10) to form DNA-calcium phosphate precipitates. These precipitates were added dropwise to the plated cells, which were incubated at 37°C for 24 hours. The medium was then replaced with fresh DMEM/10% FBS, and viral particles were collected another 48 hours later, filtered through a 0.45 µm PVDF filter, aliquoted, and stored at -80 °C.

### Stable cell line generation

For stable cell line generation, 300,000–500,000 A375 cells were plated per well in 10- cm dishes and allowed to adhere for 24-48 hours. Polybrene (10.66 µg/mL final concentration; Sigma-Aldrich, TR-10030G) was then added to the media one hour before viral infection. For viral infection, varying amounts of unconcentrated lentiviral particles (0.5 mL, 1 mL, 2 mL, 3 mL, 4 mL, or 5 mL) were added to each plate to determine optimal lentiviral dosage (**Figure 1D**). Ultimately, consistent hSOX10 expression between lines was observed with 2 mL of mT-NLS lentivirus, and 3 mL of virus for both hSOX10 constructs. Only these three lines were used for further studies. After 24 hours of transduction, media was replaced with fresh media without polybrene. At 48 hours post-infection, media was replaced with fresh media containing puromycin (1 µg/mL). Cells were expanded in selection medium, monitored for confluence, and cryopreserved in 90% FBS and 10% DMSO once stable populations were established. Parallel lines were maintained for biotinylation validation and other assays as needed.

**Figure 1.**
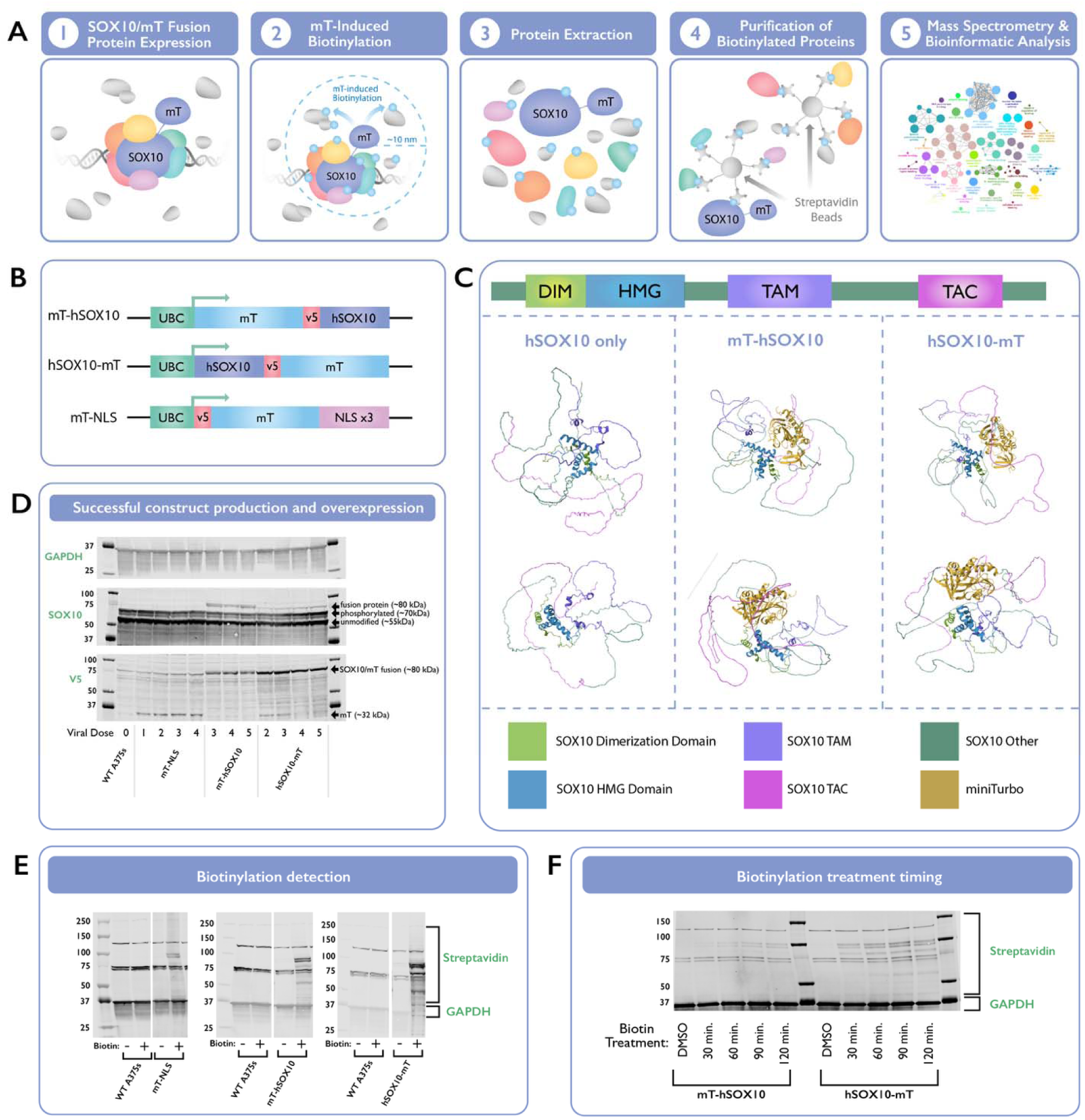
Design and validation of hSOX10-mT fusion protein constructs for proximity biotinylation and mass spectrometry. (A) Overview of the proximity biotinylation strategy utilizing mT-hSOX10 and hSOX10-mT fusion constructs to capture protein interactors in A375 melanoma cells. (B) Schematic representation of fusion protein constructs, including N-terminal (mT-hSOX10), C-terminal (hSOX10-mT) tags, and a nuclear localization signal control (mT-NLS). (C) Predicted protein structure models of hSOX10 with N-terminal and C-terminal mT fusions, generated using AlphaFold2, showing preserved structural integrity of key functional domains. (D) Western blot validation of fusion protein construct expression in A375 cells, with GAPDH as a loading control. (E) Biotinylation efficiency analysis demonstrating specific and robust labeling of hSOX10-associated proteins. (F) Optimization of biotin treatment duration, indicating peak biotinylation at 90 minutes.

### Protein biotinylation and extraction

A375 cells stably expressing hSOX10-mT, mT-hSOX10, or NLS-mT control constructs were plated in 15-cm dishes and grown to 85–90% confluence. For each construct, three biological replicates (seven plates per replicate) were used. To induce biotinylation, cells were treated with 50 μM exogenous biotin in DMEM supplemented with 10% FBS and 1 μg/mL puromycin for 90 minutes at 37°C. Following treatment, the media was discarded, and cells were washed twice with 10 mL of ice-cold PBS per plate. After rinses, cells were scraped on ice into 5 mL of PBS per plate and pooled into pre-chilled 50-mL conical tubes. Residual cells were collected by washing all plates with an additional 10 mL of PBS. For each biological replicate, the pooled cells from seven plates were combined into a single tube, resulting in nine tubes total (three replicates for each construct). The pooled cells were centrifuged at 500xg for 10 minutes at 4°C, and the supernatant was discarded. Cell pellets were resuspended in RIPA lysis buffer (50 mM HEPES pH 8, 150 mM NaCl, 2 mM EDTA, 0.1% SDS, 1% Triton X-100, 0.2% sodium deoxycholate, and 10% glycerol) supplemented with 1X Halt protease inhibitor cocktail (100X; Thermo Scientific, 1861279), 1X NEM (freshly prepared from a 1M ethanol stock; 100X; Thermo Scientific, 23030), and benzonase (1:5000 dilution; Sigma, E1014-25KU). Each tube of cell pellets (representing one biological replicate) resuspended in RIPA lysis buffer was passed through a 21-gauge needle five times to shear membranes and ensure complete lysis. Lysates were then rocked at 4°C for 15 minutes and centrifuged at 20,000xg for 5 minutes at 4°C to pellet debris. Clarified supernatants were transferred to pre-chilled tubes and stored at −80°C.

### Streptavidin pull-down

Protein concentrations were determined using the Pierce BCA Protein Assay Kit according to the manufacturer’s instructions. For each biological replicate, 20 mg of total protein was prepared in a final volume of 8 mL RIPA lysis buffer. Streptavidin-conjugated Sepharose beads (Cytiva, GE17-5113-01) were pre-washed in RIPA lysis buffer three times and centrifuged at 400xg for 2 minutes at 4°C after each wash. After washing, 30 μL of pre-washed beads was added to the protein lysates for each sample. The lysates were incubated with beads overnight at 4°C with gentle rotation to bind biotinylated proteins. Following incubation, the beads were pelleted by centrifugation at 400xg for 2 minutes at 4°C, and the unbound supernatant was discarded. The beads were washed sequentially with 1 mL wash buffers with increasingly stringency to remove non-specific binders. The washes included two rounds with 2% SDS in water (WB1), one round with a high-salt buffer (500 mM NaCl, 0.1% deoxycholate, 1% Triton X-100, 1 mM EDTA, 50 mM HEPES pH 7.5; WB2), one round with a lithium chloride buffer (250 mM LiCl, 0.5% Triton X-100, 0.5% deoxycholate, 1 mM EDTA, 50 mM HEPES pH 8.1; WB3), one round with a low-salt buffer (150 mM NaCl, 50 mM HEPES pH 7.4; WB4), and three rounds with 50 mM ammonium bicarbonate (ABC buffer; WB5). After each wash, the beads were resuspended, inverted gently 10 times, and centrifuged at 400xg for 2 minutes at room temperature. Beads were stored at 4°C or immediately processed for downstream analyses.

### Sample preparation for mass spectrometry analyses

After washing, bound proteins on streptavidin beads were digested by resuspending in 100 μL of 50 mM ammonium bicarbonate (ABC) and overnight incubation at 37°C with 1 μg of a trypsin (Promega V5113) mixed with 0.5 mAU of Lys-C (Wako Chemicals, 129-02541). An additional 0.5 μg of trypsin/Lys-C was added the next day, and digestion continued for 2 hours. Peptides were collected by centrifugation, pooled with washes from the beads, and passed through BioPureSPN columns (Nest Group, 10 µm frit, C100500) pre-washed with 50 mM ABC. Filtered peptides were acidified to 2% formic acid (FA) and dried in a SpeedVac. Dried peptides were resuspended in 25 μL of 0.1% FA and 2% acetonitrile in water. Peptide concentrations were measured with a fluorometric BCA assay, and samples were normalized to 1 μg of peptide per 10 μL injection volume. Final samples were transferred to autosampler vials for mass spectrometry.

### Western blotting

Protein lysates prepared from A375 cells stably expressing hSOX10-mT, mT-hSOX10, or NLS-mT control constructs were analyzed by western blotting to validate bait protein expression, biotinylation, and selected interactors. Proteins were separated on 4–12% Criterion™ XT Bis-Tris Protein Gels (Bio-Rad, 3450124) with an 18-well format and a load volume of 30 µL per well. Protein concentration in lysates was determined using the Pierce BCA Protein Assay Kit, and all samples were normalized based on the lowest concentration to ensure consistent loading. Protein samples were mixed with 4X LDS loading buffer (Invitrogen, NP0007) and lysis buffer to a final volume of 19 µL, and 1 µL of 20X DTT (1M, Cell Signaling, 7016) was added immediately before loading. Samples were heated at 70°C for 5–10 minutes, cooled, centrifuged briefly, vortexed, and loaded onto the gel alongside the Precision Plus Protein Kaleidoscope Prestained Protein Standards ladder (Bio-Rad, 1610395) Electrophoresis was performed using 1X MOPS Running Buffer (Invitrogen, NP0001) prepared by diluting 20X stock in deionized water. The gel was run at 80 V for 20–30 minutes, followed by 120 V for the remaining 1–1.5 hours to achieve optimal protein separation. Proteins were transferred onto a nitrocellulose membrane (Bio-Rad, 1704271) using the Trans-Blot Turbo Transfer System (Bio-Rad) with the standard protocol for midi gels. After transfer, membranes were washed with 1X TBS and blocked in 5% BSA solution for 1 hour at room temperature. Membranes were then incubated overnight at 4°C while rocking in one of the following primary antibodies diluted in blocking buffer: hSOX10 (1:1000, Abcam, ab264405), hSOX10 (1:400, Abcam, ab227680), GAPDH (1:1000, Cell Signaling, 14C10), Anti-V5 antibody (Invitrogen, R960-25), YAP1 (1:1000, Cell Signaling, 4912S), ZNF609 (1:1000, SigmaMillipore, HPA040742), TRIM24 (1:2000, Bethyl, A300-815A-T), MAML1 (1:1000, Cell Signaling, 4608S), STAT3 (1:1000, Cell Signaling, 79D7 #4904), and KMT2a (1:1000, Cell Signaling, 14197). Following primary antibody incubation, membranes were washed four times for 5-10 minutes each with 1X TBS-T. Secondary antibodies were prepared in blocking buffer containing 0.1% Tween-20 and included IRDye 680RD Streptavidin (1:5000, LI-COR), IRDye 800CW Donkey anti-Mouse (1:10,000, LI-COR, 926-32212), and IRDye 680RD Goat anti-Rabbit IgG (1:10,000, LI-COR, 926-68071). Membranes were incubated with secondary antibodies for 1 hour at room temperature and washed four times for 10-20 minutes per wash with 1X TBS-T. Dual-channel fluorescence imaging (700 and 800 nm) was performed using a LI-COR Odyssey Imaging System.

### Mass spectrometry data acquisition

Trypsinized peptides were loaded onto a µPAC Trapping column (Thermo Scientific, COL-TRPNANO16G1B2) and separated on 50 cm µPAC Neo HPLC column (Thermo Scientific, COL-NANO050NEOB). The chromatographic parameters were as follows: initial 2.8 min gradient from 2% to 10% buffer B flowing at 0.750 µL/min, then at 4.5 minutes increased to 12% buffer B and the flow rate was dropped to 300 nL/min, followed by a 57.7 min gradient from 12% to 40% buffer B flowing at 300 nL/min. Buffer B was then ramped to 90% over 0.3 minutes and the column was washed and re-equilibrated. Mass spectrometry analysis was performed on an Orbitrap Eclipse (Thermo Fisher Scientific) operated in data-dependent acquisition mode. The MS1 scans were acquired in the Orbitrap at 240k resolution, with a 250% normalized automated gain control (AGC) target, auto max injection time, and a 375-2000 m/z scan range. MS2 targets were filtered for charges states 2-7, with a dynamic exclusion of 60 seconds, and were accumulated using a 0.7 m/z quadrupole isolation window. MS2 scans were performed in the ion trap at a turbo scan rate following higher energy collisional dissociation at 35% normalized collision energy. MS2 scans used a 100% normalized AGC target and 35 ms max injection time.

### MaxQuant data search

All raw mass spectrometry data files were analyzed for protein identification and label-free quantification (LFQ) using MaxQuant (version 2.5.1.0) (Tyanova S et al., 2016a) using the default settings for “Orbitrap” instruments with minor modifications. The datasets for the N- and C-terminal mT-tagged hSOX10 experiments and the nuclear localized mT biotin ligase control were searched against the human UniProt reference proteome database (UP000005640_9606, updated 2024-02-08, https://www.uniprot.org/proteomes/UP000005640) plus a list of likely contaminants containing streptavidin, Trypsin, Albumin, mT (with different subcellular localizations), and against the contaminants list of MaxQuant. The search parameters were as follows: Methionine oxidation (+15.994914 Da) and protein N-terminal acetylation (+42.010564 Da) were set as variable modifications with a maximum of five modifications per peptide. Digestion was set to Trypsin/P with a maximum of four missed cleavages, and a minimum allowed peptide length of seven. Both unique and razor peptides were used for protein quantification, and the minimum LFQ ratio count was set to 2. Fast LFQ with classic normalization was active with a minimum and an average number of neighbors of 3 and 6, respectively. Match between runs for samples within a group (with a match time window of 0.4 min and a 20 min alignment time window) and second peptides were enabled. The raw mass spectrometry proteomics data has been deposited to the ProteomeXchange Consortium (http://proteomecentral.proteomexchange.org) with the dataset identifier PXD067868.

### Perseus data analysis

Data filtering and statistical analysis for the proximity dataset were done using Perseus (version 2.0.11.0).^27^ The ‘proteinGroups.txt’ output file from MaxQuant was loaded into Perseus using raw intensities for all samples as the Main columns. The data matrix was filtered to remove all proteins labeled as “Only identified by site”, “Reverse”, and “Potential contaminant”. The intensity values were transformed to log_2_(x), and the individual columns were grouped by Bait. Proteins without quantified/valid values in all three replicates for at least one the groups were removed. And the remaining missing values were imputed using the “Replace missing values from normal distribution” function with a width of 0.3 and a downshift of 1.8 from the standard deviation for the total matrix. Principal component analysis was performed to study the distribution of mT-hSOX10, hSOX10-mT and NLS-mT bait replicates. To identify proteins enriched in mT-hSOX10 and hSOX10-mT groups against the NLS-mT control group, two-sided unpaired Student’s t-tests were performed with a permutation-based FDR calculation (250 permutations; FDR = 0.05). Volcano plots were generated using -Log_10_ p-value (y-axis) vs. Log_2_ fold change (x-axis) values. Protein changes were considered significant when the Log_2_ fold change (hSOX10 vs. NLS) >1.0 or <-1.0 and the adjusted p-value (q-values) <0.05.

### Protein structure prediction and visualization

To predict the structural consequences of mT tagging on hSOX10, we used the COSMIC² platform at the San Diego Supercomputer Center, which utilized AlphaFold2 (v2.3.2) for highly accurate protein structure modeling. Structural predictions of N-terminal and C-terminal mT-tagged hSOX10 were generated and analyzed to assess potential conformational changes. To visualize and annotate the predicted protein structures, Protein Data Bank (PDB) files were uploaded into the *Mol 3D Viewer*\* on the RCSB PDB. Key functional domains—including the dimerization, high-mobility group, and transactivation domains—were examined for structural integrity in both fusion constructs.

### Network analysis and functional annotation with Cytoscape

To visualize and analyze the hSOX10 interactome, we generated PPI networks using Cytoscape (v3.10.3). Only the top 100 interactor candidates (ranked by highest FC) were mapped for figure clarity. Initial network layout was generated using the yFiles Organic Layout, after which protein complexes and functional groups identified by the STRING database were manually selected and rearranged with attribute circular layout to sort by FC. Mutation frequencies in melanoma (cBioPortal) were collected and incorporated into the network, with node size reflecting mutation percentages. The BioGRID database was used to differentiate known versus novel hSOX10 interactors, with node shape indicating this classification. Additionally, color shading represented the average FC of each interactor based on our dataset.

For functional enrichment analysis, all 213 high-confidence candidate interactors were input into ClueGO in Cytoscape, which performed Reactome pathway enrichment with Medium- Global Specificity. Of these, 144 proteins had functional annotations within Reactome pathways. The final 21 functionally enriched groups were defined based on KappaScore clustering, providing a structured network of enriched biological pathways.

## RESULTS

To identify novel hSOX10 interactors, we employed a proximity-dependent biotinylation strategy using mT fused to hSOX10 and a nuclear localization signal (NLS) control (**Figure 1A).** This approach enables biotinylation of proteins within a 10-nanometer radius of the fusion protein, allowing efficient labeling, purification, and identification of putative hSOX10 interactors through streptavidin-based purification and MS.

### Generation of mT fusion constructs and protein structure predictions

Three lentiviral constructs were designed to probe the hSOX10 interactome in human A375 melanoma cells: mT-hSOX10, hSOX10-mT, and mT-NLS (negative control) (**Figure 1B**). Both N- and C-terminal mT-tagged constructs were generated to identify robust and specific proximal proteins. The mT-NLS control accounts for non-specific interactions and background biotinylation unrelated to hSOX10. All constructs included a V5 tag for detection and a constitutive UBC promoter for close to endogenous hSOX10 expression.

To evaluate the structural implications of mT fusion to hSOX10, we used the COSMIC² platform at the San Diego Supercomputer Center, leveraging AlphaFold2 for protein structure prediction. **Figure 1C** visualizes the structural domains of hSOX10, including the dimerization domain (DIM), high-mobility group (HMG) domain, and transactivation domains (TAM and TAC). Predicted models indicated that the overall hSOX10 structure remains largely intact in both tagging configurations, with minor positional changes primarily observed in disordered regions, which are predicted with low or very low confidence scores (Low: 70 > pLDDT > 50; Very low: pLDDT < 50). Based on measurements obtained from these predicted 3D models, the estimated fusion protein diameter ranged from ∼5-10 nm. Given that mT-mediated proximity labeling occurs within an estimated 10-40 nm radius, these fusion protein dimensions fall well within the labeling range and suggest that N- and C-terminal tags may access partially distinct but overlapping protein microenvironments. This supports the complementary nature of the two tagging orientations in capturing the full hSOX10 interactome.

Together with the fact that the mT proximity labeling system has been widely applied across diverse biological contexts (e.g., mammalian cells, plants, and microorganisms) yielding significant insights into PPI networks,^28^ these findings suggest that the functional core of hSOX10 is preserved in both N- and C-terminal fusions, supporting their utility for interactome mapping.

### Successful generation and overexpression of mT fusion constructs in A375 melanoma cells

Western blot analysis confirmed stable overexpression of all constructs in A375 human melanoma cells. Multiple lentiviral doses were initially tested to determine the minimum viral load required for sufficient fusion protein expression. Band intensities appeared comparable across the doses used, suggesting that higher viral volumes did not markedly enhance expression (**Figure 1D**). To minimize viral usage while ensuring consistent overexpression, 3 mL of unconcentrated lentivirus was selected to generate each stable line. In **Figure 1D**, the top blot for GAPDH (loading control) shows similar band sizes and intensities across all lanes, indicating consistent protein loading across WT A375, mT-NLS, mT-hSOX10, and hSOX10-mT samples.

The middle blot for hSOX10 shows that endogenous hSOX10 bands (both unmodified and phosphorylated) remain consistent in size and intensity across all lanes, suggesting that lentiviral expression did not significantly alter endogenous hSOX10 levels. Additionally, fusion protein bands are observed exclusively in the mT-hSOX10 (lanes 7-9) and hSOX10-mT (lanes 10-13) samples, confirming successful expression of the tagged constructs.

The bottom blot for V5 (fusion protein detection) further validates construct expression. In mT-NLS control lanes (lanes 3-6), a band is detected around 32 kDa, corresponding to the expected size of the mT-NLS protein. In contrast, both mT-hSOX10 and hSOX10-mT samples (lanes 7-13) exhibit bands around 80 kDa, consistent with the predicted molecular weight of the hSOX10 fusion proteins. Within the mT-hSOX10 group (lanes 7-9), bands appear uniform in size and intensity to one another, and a similar pattern is seen in the hSOX10-mT samples (lanes 10- 13). However, the hSOX10-mT bands appear thicker and more intense, suggesting slightly higher levels of fusion protein overexpression in these samples. Despite these differences, lentiviral doses did not appear to significantly impact overall protein levels.

Notably, the fusion protein bands in the hSOX10 blot do not fully align with those detected in the V5 blot, which may reflect differences in epitope accessibility between the two antibodies. To ensure reliable detection of hSOX10, multiple commercial antibodies were tested, and the polyclonal hSOX10 antibody (ab264405) provided the most reproducible and robust results. This antibody targets the last 51 amino acids (416-466) of the hSOX10 C-terminus, immediately adjacent to the mT/V5 tag in the hSOX10-mT fusion construct. This configuration may lead to steric hindrance or epitope masking, reducing antibody accessibility. Consistent with this, the antibody showed stronger detection in mT-hSOX10 samples (where the C-terminal epitope should be unaltered) compared to hSOX10-mT, despite the opposite pattern observed in the V5 blot. This discrepancy underscores the importance of tag positioning when selecting antibodies for fusion protein validation.

### Biotinylation efficiency and optimization of biotin treatment timing

To confirm functionality of the mT tag fused to hSOX10, biotinylation efficiency was evaluated using western blot analysis of proteins extracted from biotin-treated A375 melanoma cells expressing mT-hSOX10, hSOX10-mT, or mT-NLS (**Figure 1E**). Streptavidin-based detection revealed strong and specific biotinylation patterns for both hSOX10 fusion constructs, with more extensive banding again observed in hSOX10-mT compared to mT-hSOX10. This difference may reflect higher expression levels of the hSOX10-mT construct, slight alterations in hSOX10 activity, or variations in biotinylation efficiency due to the positioning of the mT placement. The mT-NLS control samples exhibited a distinct biotinylation pattern compared to the hSOX10 fusion constructs, apart from the bands corresponding to endogenous biotinylated proteins, which are expected in eukaryotic cells.^28–31^ Specifically, these bands observed at ∼75 kDa and ∼125 kDa align with mitochondrial carboxylases, which naturally incorporate biotin as a cofactor and lead to background signals in streptavidin-based detection assays. These data confirm the utility of mT-NLS as a reliable negative control for specificity.

To optimize biotin treatment conditions, biotinylation treatments were tested at 30, 60, 90, and 120 minutes, alongside DMSO/no-biotin controls (**Figure 1F**). Streptavidin-IRDye analysis revealed a progressive increase in biotinylated protein levels with increased treatment times, peaking at 90 minutes. Based on these results, a 90-minute biotin treatment was chosen for subsequent experiments to maximize biotinylation and minimize non-specific background signal.

### Identification and characterization of the hSOX10 protein interactome using MS

To identify candidate interactors of hSOX10, MS analysis was employed following proximity biotinylation using the mT-hSOX10, hSOX10-mT, and mT-NLS control cell lines. Initial analyses applied relatively permissive thresholds to broadly assess proteins enriched in hSOX10-biotinylated interactomes compared to the NLS control. Specifically, a two-sample t-test was conducted between each tagged hSOX10 construct and the control, with proteins considered enriched if they met a log fold change ≥ 1 or ≤ –1 and an adjusted p-value (q-value) ≤ 0.05 (**Figure 2**, **A-C**). Principal component analysis (PCA) revealed tight clustering within replicates of each construct, demonstrating high reproducibility, while distinct separation between the mT-hSOX10, hSOX10-mT, and NLS-mT groups confirmed construct-specific interactome profiles (**Figure 2A**). Peptide quantification revealed an average of 1,212±187.45 peptides per replicate in NLS-mT controls, 2,465±226.69 peptides per replicate in mT-hSOX10 samples, and 5,066±430.50 peptides per replicate in hSOX10-mT samples (**Figure 2B**). The majority of proteins enriched in the mT-hSOX10 interactome were also found in the hSOX10- mT dataset, despite the observed differences in peptide yields. However, the lower peptide count for mT-hSOX10 samples suggests that some interactors may have been below the level of detection threshold for this construct.

**Figure 2.**
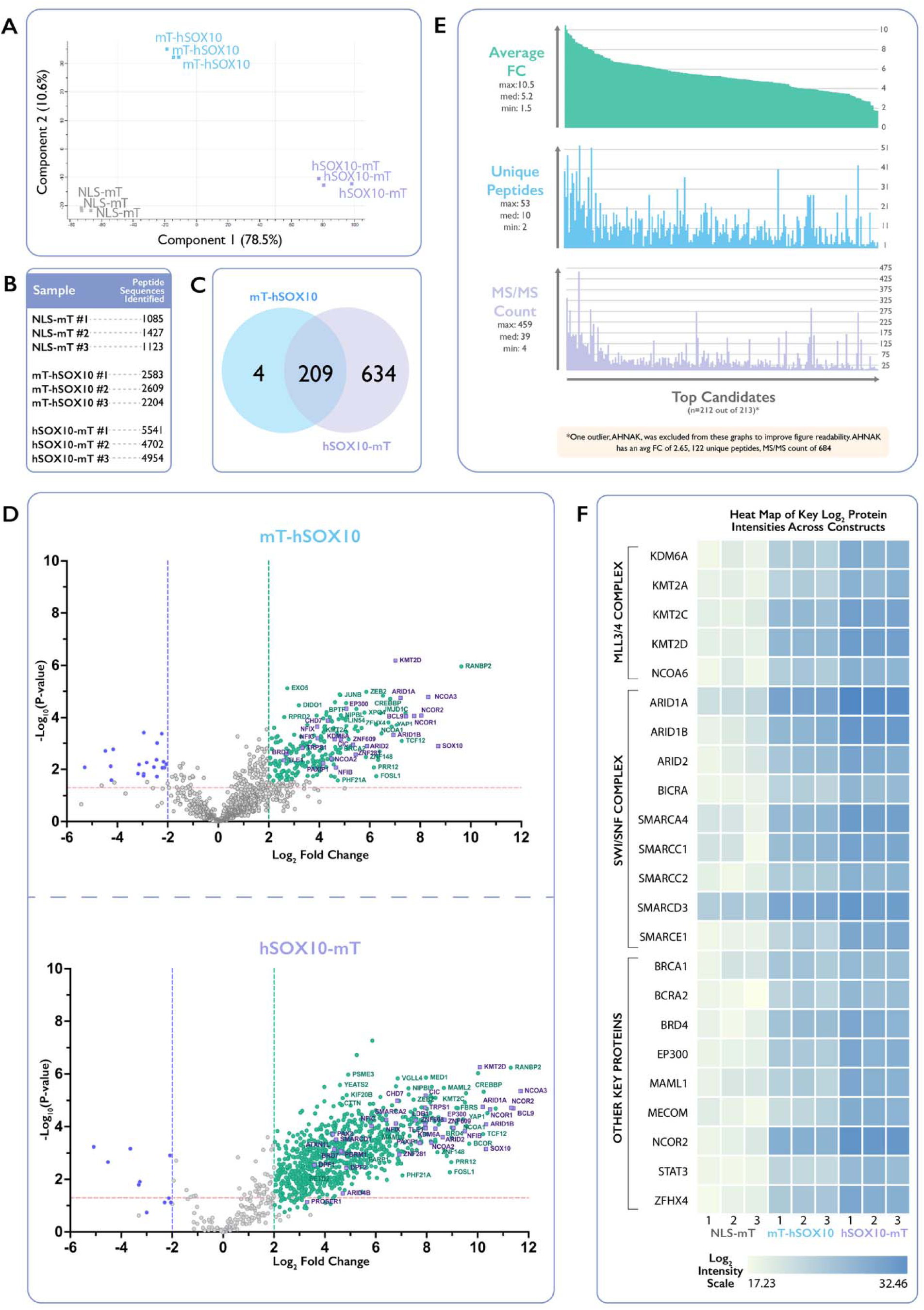
Identification of hSOX10 protein interactors using proximity biotinylation and mass spectrometry. (A) Principal Component Analysis (PCA) showing distinct clustering of mT-hSOX10, hSOX10-mT, and mT-NLS control samples based on MS data. (B) Quantification of peptides detected per sample, highlighting enrichment in mT-hSOX10 and hSOX10-mT constructs compared to controls. (C) Venn diagram of upregulated proteins in N-terminal and C-terminal hSOX10-mT interactomes versus mT-NLS control, revealing 209 shared candidates. (D) Volcano plots displaying fold change and statistical significance of enriched proteins, highlighting novel interactors (green) and known hSOX10 interactors (purple). (E) Summary metrics of top 213 interactors, excluding AHNAK as an outlier. (F) Heat map of log_2_ protein intensities for key candidate interactors, including members of the MLL3/4 and SWI/SNF complexes.

Overlap analysis of the 847 enriched proteins from N- and C-terminal hSOX10 interactomes (compared to mT-NLS controls) identified 209 shared interactors, with 4 proteins unique to mT-hSOX10 and 634 unique to hSOX10-mT (**Figure 2C**). These findings underscore both the strong similarity between the two interactomes and potential for orientation-specific differences in the proteins captured.

To refine the dataset and identify the highest-confidence interactors, additional stringent filtering criteria were applied to both the N- and C-terminal construct datasets (**Figure 2D-F**).

Protein changes were considered significant when the Log_2_ fold change (hSOX10 vs. NLS) was greater than 2.0 or less than -2.0 and the adjusted p-value (q-values) was less than 0.05. Volcano plots highlight proteins significantly enriched in mT-hSOX10 an hSOX10-mT interactomes relative to the NLS-mT controls (**Figure 2D**). Green circles denote novel hSOX10 interactors, while purple squares indicate previously known interactors. For figure clarity, labels are provided only for a selection of representative proteins, especially those that belong to known protein complexes (see below and **Figure 4**). However, a comprehensive list of interactors is available in **Supplementary Table 1**. For both fusion proteins, enriched proteins included previously identified interactors such as KMT2D, EP300, ARID1A, and NCOR2, as well as novel interactors like SMARCA4, KMT2C, MECOM and ZFHX4, to be discussed below (**Figure 2F**). From this stringent analysis, 213 candidates were identified, comprising 209 shared between mT-hSOX10 and hSOX10-mT datasets and 4 unique to mT-hSOX10. These proteins were ranked based on average Log_2_ fold-change (FC). A summary of key dataset metrics of this data is provided in **Figure 2E** with one outlier, AHNAK (average Log_2_ FC = 2.65, 122 unique peptides, MS/MS count = 684) excluded from visualization to improve readability. Across the 212 remaining proteins, the average Log_2_ FC values for proteins ranged from 1.5 to 10.5, with a median of 5.2. Unique peptide counts ranged from 2 to 53, with a median of 10. MS/MS counts ranged from 4 to 459, with a median of 39. Together, these findings demonstrate the robustness of proximity biotinylation and MS-based interactome mapping, enabling the reliable identification of putative hSOX10-associated proteins complexes and pathways.

### Validation of hSOX10 candidate interactors using western blotting

To validate candidate hSOX10 interactors identified through MS, western blot analysis was performed using biotinylated protein samples from A375 cells expressing mT-NLS, mT-hSOX10, or hSOX10-mT constructs (**Figure 3A**). Seven interactors with highly validated antibodies were selected for confirmation: hSOX10 itself, YAP1 (average Log_2_ FC = 8.25), MAML1 (average Log_2_ FC = 5.55), ZNF609 (average Log_2_ FC = 7.15), TRIM24 (average Log_2_ FC = 4.05), STAT3 (average Log_2_ FC = 5.85), and KMT2A (average Log_2_ FC = 5.95). As expected, hSOX10 blotting confirmed its strong presence in both mT-hSOX10 and hSOX10-mT samples, but not mT-NLS controls. Among the candidate interactors, YAP1, MAML1, ZNF609, TRIM24, STAT3, and KMT2A were all detected with varying degrees of signal enrichment in the hSOX10-tagged samples, compared to the mT-NLS control. Consistent with the differences in expression levels between the N-terminal and C-terminal mT fusions, western blot signals were generally weaker in mT-hSOX10 samples than in hSOX10-mT samples (e.g. YAP1, ZNF609 and MAML1). For STAT3, two distinct bands were detected in both hSOX10-mT and mT-hSOX10 samples, with stronger signals again observed in hSOX10-mT samples, whereas mT-NLS controls showed only a faint band, indicating specific enrichment in hSOX10 proximity interactions. KMT2A, the largest protein (∼180 kDa) among the validated interactors, exhibited faint enrichment in hSOX10-mT samples and was minimally detectable in mT-hSOX10 samples. Overall, these western blot results provide experimental validation and quality control for select candidate hSOX10 interactors identified in MS-based interactome mapping.

**Figure 3.**
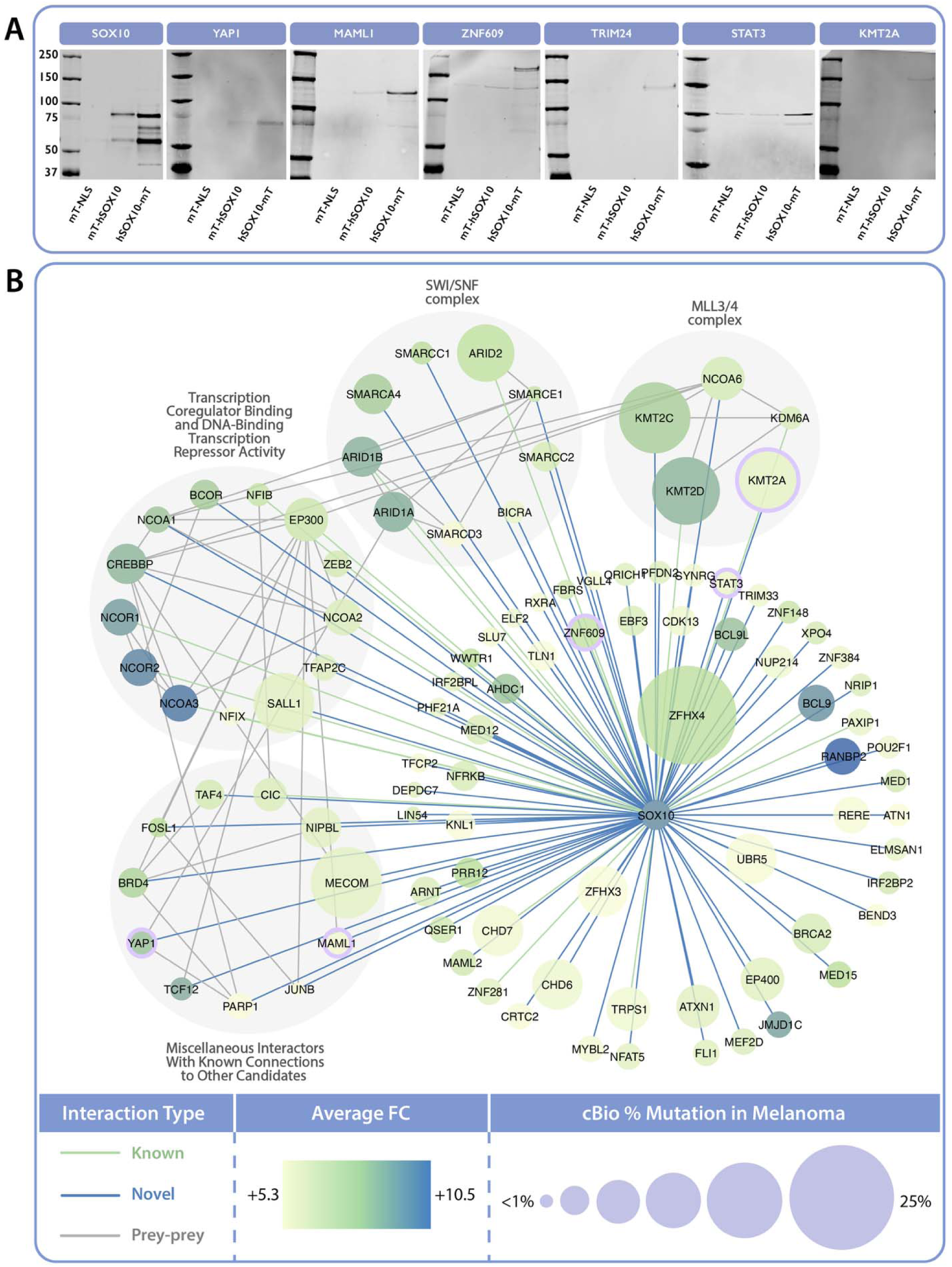
Western blot validation and network analysis of candidate hSOX10 protein interactors identified by proximity biotinylation and mass spectrometry. (A) Immunoprecipitation (IP)-Western blot validation of six candidate hSOX10 interactors spanning a range of enrichment levels (YAP1, MAML1, ZNF609, TRIM24, STAT3, and KMT2A). Biotinylated proteins purified with streptavidin beads from mT-NLS, mT-hSOX10, and hSOX10- mT expressing cells were probed with protein-specific antibodies. Blotting for hSOX10 is shown as a control to confirm the presence of the bait protein. (B) Network visualization of hSOX10 interactors identified by proximity biotinylation and mass spectrometry. Nodes represent individual proteins, and edges represent known interactions from BioGRID. Node size correlates with mutation frequency in melanoma from cBioPortal data (<1% to 25%), and node color represents the average fold change (FC) enrichment in hSOX10-proximity experiments, with a gradient from green (+5.3) to blue (+10.5). Known hSOX10 interactions are represented by green edges, novel interactions by blue edges, prey-prey interactions by grey edges, and nodes with purple outlines indicate proteins validated by IP-Western blot in panel A. Key complexes, including SWI/SNF and MLL3/4 complexes, as well as transcriptional regulators and additional grouped interactors, are highlighted.

**Figure 4.**
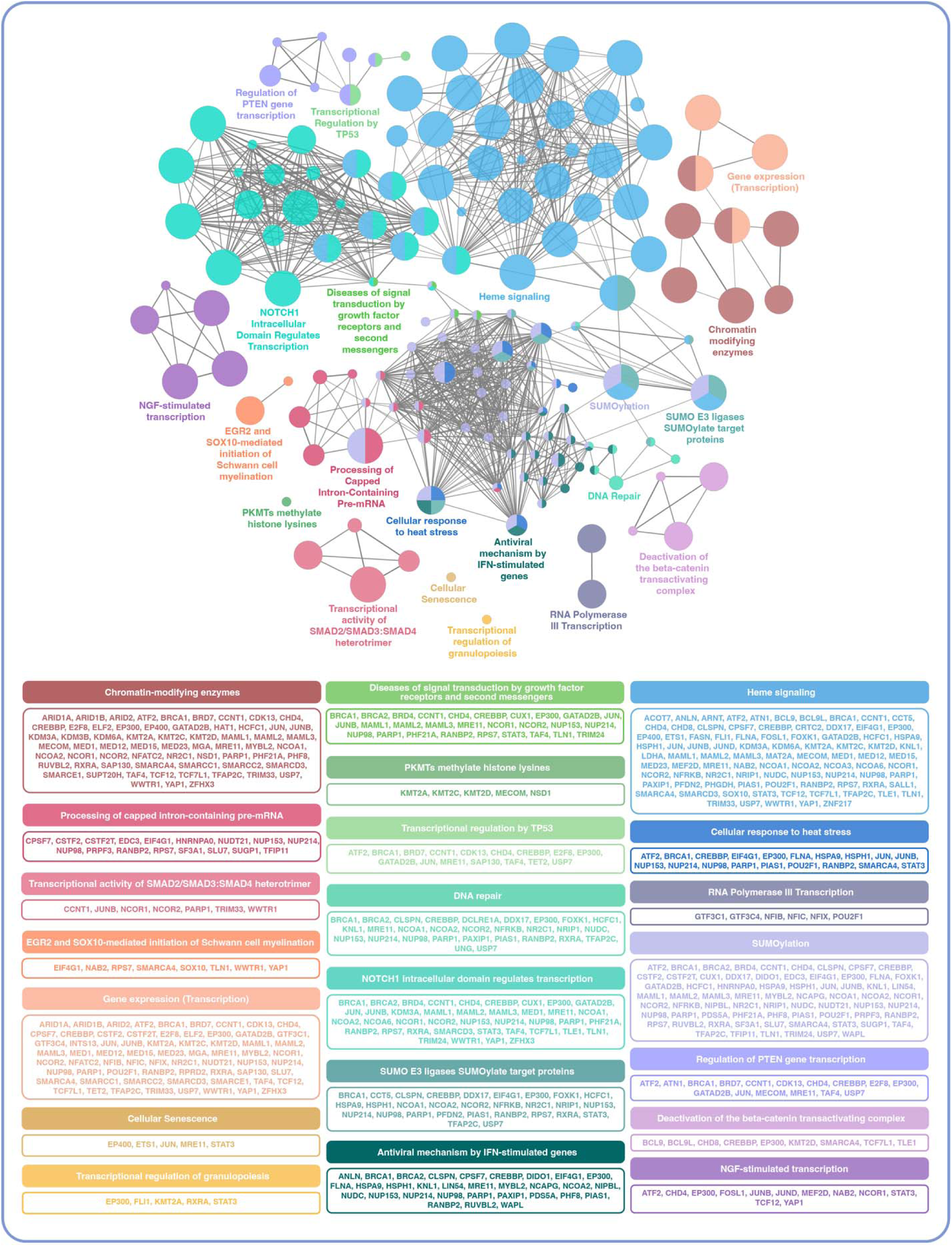
Functional enrichment network of hSOX10 protein interactors reveals key biological pathways. Functional enrichment analysis of hSOX10 protein interactors generated using ClueGO in Cytoscape with REACTOME pathways for data analysis. Nodes represent enriched pathways, with node size corresponding to the number of associated proteins, and edge thickness indicating the degree of shared proteins between pathways. Pathways are grouped by color to represent biological themes.

### Analysis of hSOX10 proximity network, potential key interactors and enriched complexes

The proximity interactome of hSOX10, identified through biotinylation and MS, was visualized as a network using Cytoscape to highlight functional associations among candidate interactors (**Figure 3B**). For figure readability, this network includes only the top 100 candidates ranked by highest Log_2_ FC. Both novel (circular nodes) and known (square nodes) hSOX10 interactors are represented, with node sizes corresponding to mutation frequency in melanoma according to cBioPortal. Gradient shading of the nodes indicates average Log_2_ FC (abundance over control), ranging from lighter green (+5.3) to darker green (+10.5).

Several functionally related groups and protein complexes were enriched in the hSOX10 interactome. The SWI/SNF complex, a chromatin-remodeling complex, was prominently enriched. Members of this complex, including SMARCC1, SMARCC2, SMARCA4, and ARID1A/ARID1B, displayed high FC values and/or high mutation rates in cBioPortal. Another enriched group was the MLL3/4 (or COMPASS) complex, involved in histone methylation and transcriptional activation. Proteins such as KMT2A, KMT2C (MLL3), and KMT2D (MLL4), key members of this complex, display high mutation frequencies in melanoma (13%, 17%, and 16%, respectively). The network also highlighted transcriptional co-regulators, such as NCOR2, SALL1, and EP300, and proteins with established connections to multiple other hSOX10 interactome components (e.g., BRD4, MECOM, and NIPBL) that are mutated in a notable percentage of melanomas (5%, 17%, and 8%, respectively). Interestingly, this network also highlights ZFHX4, a zinc finger transcription factors that is mutated in 25% of melanomas but remains largely unexplored. Together, these findings validate previously discovered hSOX10 interactors and identify novel candidates, providing a high confidence interactome that highlights hSOX10’s role in transcriptional and chromatin regulatory networks.

### Functional enrichment of hSOX10 interactome network highlights key biological pathways

To further explore the biological functions of the hSOX10 interactome, a functional enrichment analysis of the top 213 candidate interactors was performed using ClueGO in Cytoscape, with Reactome pathways serving as the reference database. The results include an interactome map that illustrates functional relationships among enriched pathways and a detailed breakdown of genes contributing to each pathway (**Figure 4**). In this network, nodes represent specific biological pathways, with node size reflecting the number of contributing genes, while connections between nodes indicate shared gene associations, revealing functionally linked processes within the hSOX10 interactome.

Several biologically significant pathways were enriched, linking hSOX10’s to chromatin-modifying enzymes (e.g., KMT2A, KMT2C, BRD7, and NCOR2), heme signaling (e.g., BRCA1, CHDs, ETS1, and MAML3), SUMOylation (e.g., BRCA1/2, BRD4, CHD4, and JUN),

DNA repair (e.g., BRCA1/2, EP300, NCOA1/2, and TFAP2C), NGF-stimulated transcription (e.g., ATF2, CHD4, JUNB, and STAT3), and cellular stress responses (e.g., ATF2, EP300, HSPs, and PARP1). Together, these findings establish a functional framework for the hSOX10 interactome, linking it to key regulatory pathways involved in melanoma progression.

## DISCUSSION

This study represents the first comprehensive characterization of the hSOX10 interactome in human melanoma cells, leveraging mT-based proximity biotinylation and MS. We identified 847 candidate interactors, of which 213 met stringent criteria for high-confidence inclusion based on fold change and statistical significance. These proteins were enriched in pathways related to chromatin remodeling, transcriptional regulation, SUMOylation, and stress response, all of which are implicated in melanoma biology. Notably, our dataset revealed proximity of hSOX10 to several proteins that are recurrently mutated in melanoma, including KMT2C, KMT2D, BRD4, ZFHX4, and STAT3, suggesting that genetic alterations within the hSOX10 interactome may rewire downstream transcriptional programs. While hSOX10 itself is not frequently mutated in melanoma, its dysregulation through altered expression, enhancer rewiring, or post-translational modification is well-documented and contributes to tumor growth, phenotype switching, and therapeutic resistance. This study serves as a proteomics-based resource to support hypothesis generation and guide future investigations into hSOX10’s mechanistic role in melanoma progression and plasticity.

### SWI/SNF complex: a pivotal role in melanoma biology

The SWI/SNF chromatin-remodeling complex emerged as a prominent component of the hSOX10 interactome, with nine members ranked among the top 100 interactors (**Figure 4**). These include known hSOX10 partners (e.g., ARID2, ARID1B, and ARID1A) and novel candidates (e.g., SMARCA4, SMARCC1, SMARCE1, SMARCC2, BICRA, and SMARCD3).

Given the essential role of SWI/SNF in chromatin accessibility and lineage specification, its proximity to hSOX10 suggests a potential convergence of transcriptional and epigenetic regulation. Mutations in SWI/SNF subunits are common in melanoma and are known to alter transcriptional control, promote progression, and influence therapeutic resistance, particularly in the context of immune checkpoint inhibitors and synthetic lethality.^32–42^ The findings presented here nominate SWI/SNF components as candidates for deeper investigation in the context of hSOX10-regulated transcriptional programs.

### MLL3/4 complex: linking enhancer dysregulation to melanoma

STRING network analysis also highlighted the MLL3/4 complex, a COMPASS-family enhancer regulator, as a central component of the hSOX10 interactome. Although the role of this complex in melanoma remains largely unexplored, multiple core members were identified in our dataset, including MLL3 (KMT2C) and MLL4 (KMT2D).^43,44^ Mutations in MLL3 and MLL4 are among the most frequent genetic alterations in bladder, colorectal, and breast cancers, as well as hematologic malignancies, where they typically lead to loss of function.^43–46^ These mutations disrupt enhancer activity by reducing H3K4 mono-methylation and H3K27 acetylation, histone marks essential for active enhancer states. Additionally, MLL3/4 loss-of-function mutations create metabolic dependencies, such as heightened reliance on purine synthesis, which may represent therapeutic vulnerabilities.

Although melanoma has not been extensively studied in the context of MLL3/4 function, our findings suggest its potential relevance. Two of the five MLL3/4 complex members identified in this study, KMT2D and KDM6A, have been previously reported to interact with hSOX10 (**Figure 4)**. Moreover, KMT2A, KMT2C, and KMT2D show high mutation frequencies in melanoma (13%, 17%, and 16%, respectively, per cBioPortal). Given the strong association between hSOX10 and multiple MLL3/4 complex components, further investigation is warranted to determine whether hSOX10-MLL3/4 interactions contribute to enhancer remodeling, cell state transitions, or response to epigenetic therapies in melanoma.

### Key known interactors: NCOR2 and EP300 in hSOX10 regulation

Among the known hSOX10 interactors identified in our dataset, NCOR2 and EP300 were particularly notable. NCOR2, a nuclear receptor corepressor, plays a key role in chromatin remodeling and has been implicated in melanoma prognosis.^47^ A genome-wide association study investigating functional single nucleotide polymorphisms (SNPs) in the Notch signaling pathway identified two SNPs in NCOR2 as predictors of disease-specific and overall survival in cutaneous melanoma, emphasizing its potential as a biomarker for personalized melanoma management. Given NCOR’s established role in transcriptional regulation and cancer,^47–49^ its proximity to hSOX10 in our dataset suggests a possible contribution to hSOX10-mediated transcriptional programs, consistent with broader enhancer dysregulation mechanisms in melanoma. Notably, MAML2, another gene identified in the same melanoma survival study, also appeared among our top 100 candidate interactors, further supporting its potential role in hSOX10 regulatory networks.

EP300, a lysine acetyltransferase, has been shown to interact with hSOX10, stabilizing it through acetylation and enhancing its function in melanoma proliferation and tumor growth.^50,51^ Co-amplification of EP300 and hSOX10 in melanoma tumors further underscores their synergistic role in disease progression. Inhibition of EP300 using a small-molecule inhibitor reduces hSOX10 protein levels and suppresses melanoma proliferation and invasion by downregulating metastasis-associated genes.^50^ These findings position EP300-hSOX10 as a central regulatory axis in maintaining the epigenetic landscape that drives melanoma growth and plasticity.

Together, these observations reinforce the functional relevance of these known hSOX10 interaction partners and support their continued exploration as co-regulators of hSOX10 activity.

### Novel candidate interactors of hSOX10

Beyond the SWI/SNF and MLL3/4 complexes, several novel hSOX10 candidate interactors identified in this study—including MECOM, ZFHX4, BRCA1/2, and BRD4— highlight potential new regulatory pathways in melanoma biology.

MECOM, a transcriptional regulator, plays key roles in enhancer activity, lineage identity, and tumor progression.^52–55^ Its functions in stemness and chromatin remodeling suggest overlap with hSOX10-regulated transcriptional networks in melanoma. In leukemia and solid tumors, MECOM promotes proliferation and inhibits apoptosis via AKT/mTOR signaling and polycomb interactions. In lung squamous cell carcinoma, it drives growth and stemness through SOX2 regulation, with CRISPR-mediated MECOM depletion significantly reducing tumor burden.^52^ MECOM also activates endothelial identity genes (e.g., VEGF pathway) through enhancer binding, establishing its broader role in lineage specification.^55^ In melanoma, it acts as a secondary driver, especially in BRAF-mutant tumors, modulating immune checkpoint inhibitor responses and the epigenetic tumor microenvironment.^56^ Given hSOX10’s role in melanoma identity and transcriptional plasticity, its interaction with MECOM suggests a potential mechanism for therapy resistance and tumor progression, warranting further investigation.

ZFHX4, a zinc-finger transcription factor, is mutated in 25% of melanomas (cBioPortal) yet remains poorly characterized in this context.^57–61^ It has been implicated in transcriptional regulation, immune modulation, and tumor progression, with a recent study linking ZFHX4 mutations to immune checkpoint inhibitor efficacy in non-small cell lung cancer and melanoma.^57^ ZFHX4-mutant tumors exhibit enrichment of DNA damage repair pathways, reinforcing its potential role in genomic instability and transcriptional adaptation. Given that hSOX10 regulates lineage identity and chromatin remodeling, its interaction with ZFHX4 may influence transcriptional plasticity and melanoma therapy resistance.

BRCA1 and BRCA2, known tumor suppressors and DNA repair regulators, were also identified in our dataset, suggesting a potential, albeit previously unestablished, link to hSOX10- mediated transcriptional control. While BRCA1 mutations have not been consistently associated with melanoma, BRCA2 mutations are linked to a 2.5- to 2.7-fold increased risk of familial melanoma, likely due to homologous recombination deficiencies driving genomic instability in UV-exposed melanocytes.^62–65^ BRCA mutations also alter p53-related genomic stability pathways, which may further promote melanoma progression.^64^ Our data suggest that BRCA1/2 may influence hSOX10-regulated cell identity programs, potentially bridging transcriptional plasticity and DNA repair deficiencies in melanoma.

BRD4, a Bromodomain and Extraterminal (BET) family chromatin regulator, emerged as a key hSOX10 interactor, consistent with its role in transcriptional activation, enhancer function, and melanoma progression.^66–70^ BRD4 regulates super-enhancer activity, driving c-MYC expression and melanoma proliferation. Inhibiting BRD4 with BET inhibitors reduces melanoma growth, even in BRAF- and NRAS-mutant tumors, highlighting its therapeutic relevance.^66,69^ BRD4 also facilitates non-canonical Hedgehog/GLI signaling activation via its interaction with SOX2, a pathway implicated in melanoma metastasis.^71^ Notably, BRD4 contributes to melanoma invasion and lung metastasis through the SPINK6/EGFR-EphA2 pathway, activating signaling cascades like the ERK1/2 and AKT pathways.^66^ Given that both BRD4 and hSOX10 regulate enhancer function and transcriptional plasticity, their interaction may represent a critical regulatory axis in melanoma progression and therapy resistance. Further investigation into the BRD4-hSOX10 relationship could uncover novel therapeutic targets in melanoma epigenetics.

Collectively, these findings expand our understanding of hSOX10’s functional network, reinforcing its central role in transcriptional regulation, enhancer function, and melanoma plasticity. The identification of these novel interactors highlights potential new therapeutic vulnerabilities and underscores the importance of hSOX10-targeted research in melanoma biology.

### Pathways enriched in the hSOX10 human melanoma interactome

Beyond individual interactors, our analysis identified several biologically enriched pathways enriched in the hSOX10 interactome, including heme signaling, NOTCH1 intracellular domain (NICD)-mediated transcription, and SUMOylation. These pathways are critical regulators of transcriptional plasticity, melanoma cell identity, and tumor adaptability, positioning them as key components of hSOX10-associated regulatory networks.

Heme signaling, particularly through heme oxygenase-1 (HO-1), plays a pro-tumorigenic role in melanoma progression and therapy resistance.^72–76^ HO-1 interacts with BRAF, activating the ERK1/2 pathway to drive cell cycle progression, while its overexpression enhances tumor growth and stem cell-like properties, including non-adherent growth and vasculogenic mimicry. HO-1 also contributes to immune evasion by suppressing CD8+ T-cell infiltration, and its inhibition sensitizes melanoma cells to BRAF inhibitors, such as vemurafenib. The enrichment of heme signaling in our top 213 proteins is notable, as both HO-1 and hSOX10 regulate transcriptional plasticity, cell identity, and tumor adaptability. Investigating this relationship may reveal novel pathways underlying melanoma heterogeneity and resistance, offering potential therapeutic targets at the intersection of HO-1-driven signaling and hSOX10-mediated transcriptional control.

NOTCH1 signaling, particularly through the NICD, also emerged as an enriched pathway in our dataset. Once cleaved from the Notch1 receptor, NICD translocates into the nucleus, where it forms a transcriptional complex with RBP-J and MAML (another of our top interactors, **Figure 4**) to activate genes involved in proliferation, survival, and adhesion.^77,78^ Aberrant NOTCH1 activation has been linked to enhanced melanoma growth and aggressiveness, particularly in vertical growth phase melanoma, where it drives MAPK and PI3K-Akt signaling.^79–81^ NOTCH1 also enhances metastatic capacity and supports melanoma survival in nutrient-poor environments. Interestingly, its interaction with other factors, such as NRN1, can modulate its activity and redirect downstream signaling pathways like STAT3 (another top interactor in our dataset, **Figure 4**), suggests a broader transcriptional adaptability network in melanoma.^82^ The role of NOTCH1 in neural crest-derived lineages, from which melanocytes and melanoma originate, suggests that melanoma cells co-opt developmental programs to maintain progenitor-like states and plasticity, a mechanism reminiscent of hSOX10’s function in lineage regulation. Investigating this interplay may provide critical insights into melanoma’s transcriptional dependencies and offer new therapeutic targets aimed at disrupting these oncogenic networks.

SUMOylation, a post-translational modification involving small ubiquitin-like modifier (SUMO) proteins, emerged as another key regulatory pathway in the hSOX10 interactome. This modification is crucial for fine-tuning hSOX10 transcriptional activity in melanoma.^83,84^ SUMOylation of hSOX10 at lysine residues K55, K246, and K357 alters its interactions with transcriptional coactivators and corepressors, shifting its regulatory role. This influences tumor proliferation, survival, and plasticity, with dysregulated SUMOylation driving a slow-cycling, invasive state that contributes to resistance to BRAF and MEK inhibitors. The enrichment of SUMOylation-related pathways in our dataset underscores its importance in hSOX10-mediated melanoma progression and suggests that targeting SUMOylation machinery or related regulatory pathways could disrupt key transcriptional nodes and enhance therapeutic efficacy in melanoma.

### Unveiling the hSOX10 interactome: implications for melanoma biology and therapy

This study presents the first high-resolution proteomic map of the hSOX10 interactome in melanoma, advancing our understanding of hSOX10’s role as a transcriptional regulator in this aggressive skin cancer. By identifying novel candidate interactors and enriched regulatory pathways (e.g., chromatin remodeling, SUMOylation, and heme signaling), our findings position hSOX10 as a central node in melanoma transcriptional networks. These results validate the robustness of the mT-based proximity labeling approach and reveal new layers of hSOX10- mediated regulation.

Importantly, this work was designed as a broad, proteomics-focused resource to prioritize reproducibility and coverage over mechanistic detail. As such, the candidate interactors and enriched pathways serve as a hypothesis-generating framework to guide future investigations into hSOX10’s roles in enhancer regulation, chromatin remodeling, and therapy resistance. This resource is intended to accelerate the discovery of context-specific co-regulators and support downstream studies in melanoma biology and beyond.

Future work should focus on functionally characterizing these newly identified interactors and pathways to elucidate their roles in melanoma progression, cellular plasticity, and therapeutic resistance. Expanding these analyses across diverse melanoma models and incorporating orthogonal validation approaches will provide deeper insight into the transcriptional dependencies and vulnerabilities associated with hSOX10. Ultimately, understanding how hSOX10 integrates with key regulatory complexes and signaling cascades may uncover novel therapeutic targets and help overcome the adaptability that underlies melanoma’s resistance to treatment. Together, these findings lay the groundwork for precision medicine strategies aimed at disrupting hSOX10-driven transcriptional programs in melanoma.

## CONCLUSIONS

This study provides the first comprehensive map of the SOX10 interactome in human melanoma, identifying 213 high-confidence candidate protein interactors using proximity biotinylation and mass spectrometry. These include both known and novel partners involved in chromatin remodeling, transcriptional regulation, SUMOylation, and DNA repair. Notably, we identified multiple members of the SWI/SNF and MLL3/4 complexes, as well as interactors like EP300, NCOR2, BRD4, MAML1, and STAT3, highlighting key pathways through which SOX10 may influence melanoma cell identity, plasticity, and resistance. This dataset offers a valuable resource for uncovering new mechanisms of SOX10 function and identifying potential therapeutic targets in melanoma.

## CONTRIBUTIONS

C.M.N.S. and C.K.K. developed the research question and overall research plan; D.P.B. and M.B.M. provided guidance and training, experimental protocols, proteomics expertise, and key reagents; C.M.N.S executed and troubleshooted most experimental protocols; D.P.B. conducted mass spectrometry sample preparation, data acquisition, and initial computational analyses; C.M.N.S and C.K.K. conducted additional analyses and performed literature reviews; C.M.N.S. and D.P.B. prepared manuscript figures; All authors contributed to manuscript preparation.

## ACKNOWLEDGEMENTS

The authors have no acknowledgements to declare.

## ABBREVIATIONS

ABC: Ammonium Bicarbonate
AGC: Automatic Gain Control
BSA: Bovine Serum Albumin
COMPASS: Complex Proteins Associated with Set1 (histone methyltransferase complex, includes MLL3/4)
COSMIC²: Collaborative Science via Cyberinfrastructure² (platform at San Diego Supercomputer Center)
CRISPR: Clustered Regularly Interspaced Short Palindromic Repeats
DMSO: Dimethyl Sulfoxide
DMEM: Dulbecco’s Modified Eagle Medium
DTT: Dithiothreitol
EDTA: Ethylenediaminetetraacetic Acid
FA: Formic Acid
FBS: Fetal Bovine Serum
FC: Fold Change
FDR: False Discovery Rate
GRN: Gene Regulatory Network
HBS: HEPES-Buffered Saline
HMG: High-Mobility Group (domain)
HO-1: Heme Oxygenase-1
HSPs: Heat Shock Proteins
IP: Immunoprecipitation
IRF1: Interferon Regulatory Factor 1
LFQ: Label-Free Quantification
MAPK: Mitogen-Activated Protein Kinase
mAU: Milli-Absorbance Unit
mT: miniTurbo (proximity-dependent biotinylation enzyme)
MS: Mass Spectrometry
NC: Neural Crest
NICD: Notch Intracellular Domain
NLS: Nuclear Localization Signal
NP-40 (LDS buffer): Nonidet P-40 detergent; Lithium Dodecyl Sulfate loading buffer
PBS: Phosphate-Buffered Saline
PCA: Principal Component Analysis
PDB: Protein Data Bank
PD-L1: Programmed Death-Ligand 1
PPI: Protein–Protein Interaction
PVDF: Polyvinylidene Difluoride
RIPA: Radioimmunoprecipitation Assay (buffer)
SDS: Sodium Dodecyl Sulfate
SDS-PAGE: Sodium Dodecyl Sulfate–Polyacrylamide Gel Electrophoresis
SNP: Single Nucleotide Polymorphism
SOX10 / hSOX10: SRY-Box Transcription Factor 10 / human SOX10
SUMOylation: Small Ubiquitin-like Modifier conjugation
TAM/TAC: Transactivation Motifs (domains of SOX10)
TBS: Tris-Buffered Saline
TBS-T: Tris-Buffered Saline with Tween-20
TF: Transcription Factor
UBC: Ubiquitin C (promoter)
UV: Ultraviolet

