## Supplementary Data for "Mapping of the hSOX10 protein interactome in human melanoma"

**Table S1. High-confidence candidate hSOX10 interactors identified by proximity labeling and mass spectrometry.**

|  | | **mT-hSOX10_NLS-mT** | | | | | **hSOX10-mT_NLS-mT** | | | |
| --- | --- | --- | --- | --- | --- | --- | --- | --- | --- | --- |
|  |  | | **Log2 FC** | **-Log10**  **(P-value)** | **Adjusted P-value** | **Log2 FC** | | **-Log10**  **(P-value)** | **Adjusted P-value** |  |
| Majority protein IDs | Gene names | | Student's T-test Difference mT-hSOX10-NLS-mT | -Log Student's T-test p-value mT-hSOX10-NLS-mT | Student's T-test q-value mT-hSOX10-NLS-mT | Student's T-test Difference hSOX10-mT-NLS-mT | | -Log Student's T-test p-value hSOX10-mT-NLS-mT | Student's T-test q-value hSOX10-mT-NLS-mT | **Unique peptides** |
| P49792 | RANBP2 | | 9.6 | 6.0 | 0.0000 | 11.3 | | 6.2 | 0.0000 | 40 |
| P56693 | SOX10 | | 8.7 | 2.9 | 0.0056 | 10.3 | | 3.2 | 0.0000 | 9 |
| Q9Y6Q9 | NCOA3 | | 8.3 | 4.8 | 0.0000 | 11.7 | | 5.4 | 0.0000 | 27 |
| Q9Y618 | NCOR2 | | 8.0 | 4.1 | 0.0000 | 11.3 | | 4.7 | 0.0000 | 48 |
| O75376 | NCOR1 | | 7.8 | 4.1 | 0.0000 | 10.5 | | 4.7 | 0.0000 | 39 |
| O00512 | BCL9 | | 7.4 | 4.1 | 0.0000 | 11.4 | | 4.7 | 0.0000 | 29 |
| Q15652 | JMJD1C | | 7.4 | 4.2 | 0.0000 | 10.7 | | 4.7 | 0.0000 | 34 |
| Q99081 | TCF12 | | 7.3 | 3.1 | 0.0056 | 10.2 | | 3.7 | 0.0000 | 15 |
| O14497 | ARID1A | | 7.2 | 4.8 | 0.0000 | 10.2 | | 4.8 | 0.0000 | 40 |
| P46937 | YAP1 | | 7.1 | 3.7 | 0.0024 | 9.4 | | 4.3 | 0.0000 | 11 |
| O14686 | KMT2D | | 7.0 | 6.2 | 0.0000 | 10.1 | | 6.3 | 0.0000 | 53 |
| Q8NFD5 | ARID1B | | 6.9 | 3.3 | 0.0054 | 10.3 | | 4.1 | 0.0000 | 25 |
| Q15788 | NCOA1 | | 6.9 | 3.6 | 0.0045 | 9.6 | | 4.0 | 0.0000 | 17 |
| Q92793 | CREBBP | | 6.8 | 4.7 | 0.0000 | 10.2 | | 5.3 | 0.0000 | 21 |
| O60885 | BRD4 | | 6.7 | 3.8 | 0.0026 | 9.0 | | 4.0 | 0.0000 | 10 |
| Q5TGY3 | AHDC1 | | 6.5 | 4.8 | 0.0000 | 10.0 | | 6.0 | 0.0000 | 25 |
| P51532 | SMARCA4 | | 6.5 | 3.4 | 0.0057 | 9.6 | | 4.0 | 0.0000 | 21 |
| Q8NEZ4 | KMT2C | | 6.4 | 4.2 | 0.0000 | 9.5 | | 4.9 | 0.0000 | 52 |
| Q6W2J9 | BCOR | | 6.4 | 2.7 | 0.0063 | 9.7 | | 3.4 | 0.0000 | 37 |
| Q92922 | SMARCC1 | | 6.3 | 2.5 | 0.0086 | 8.7 | | 3.1 | 0.0000 | 11 |
| Q96RN5 | MED15 | | 6.3 | 2.4 | 0.0140 | 8.8 | | 2.9 | 0.0000 | 12 |
| P15407 | FOSL1 | | 6.3 | 1.7 | 0.0373 | 8.9 | | 2.3 | 0.0005 | 4 |
| Q86UU0 | BCL9L | | 6.3 | 3.7 | 0.0024 | 10.4 | | 4.5 | 0.0000 | 23 |
| Q9ULL5 | PRR12 | | 6.2 | 2.1 | 0.0213 | 9.0 | | 2.7 | 0.0000 | 29 |
| Q68CP9 | ARID2 | | 5.9 | 2.9 | 0.0057 | 8.6 | | 3.6 | 0.0000 | 18 |
| O60315 | ZEB2 | | 5.9 | 5.0 | 0.0000 | 7.5 | | 5.0 | 0.0000 | 14 |
| Q9UQR1 | ZNF148 | | 5.8 | 2.5 | 0.0097 | 8.4 | | 3.0 | 0.0000 | 18 |
| Q9C0E2 | XPO4 | | 5.8 | 4.2 | 0.0000 | 7.5 | | 4.5 | 0.0000 | 10 |
| Q86UP3 | ZFHX4 | | 5.7 | 3.6 | 0.0043 | 9.2 | | 4.1 | 0.0000 | 24 |
| Q9GZV5 | WWTR1 | | 5.5 | 3.5 | 0.0062 | 9.2 | | 4.3 | 0.0000 | 6 |
| Q15648 | MED1 | | 5.5 | 4.3 | 0.0000 | 8.0 | | 5.9 | 0.0000 | 13 |
| Q6P4R8 | NFRKB | | 5.4 | 3.7 | 0.0048 | 7.6 | | 4.1 | 0.0000 | 12 |
| Q9Y2X9 | ZNF281 | | 5.4 | 2.6 | 0.0079 | 6.9 | | 3.0 | 0.0000 | 17 |
| Q7Z5L9 | IRF2BP2 | | 5.4 | 3.6 | 0.0041 | 8.0 | | 4.2 | 0.0000 | 7 |
| O15014 | ZNF609 | | 5.3 | 2.9 | 0.0061 | 9.0 | | 4.0 | 0.0000 | 12 |
| P51587 | BRCA2 | | 5.2 | 2.9 | 0.0056 | 7.9 | | 3.9 | 0.0000 | 22 |
| Q2KHR3 | QSER1 | | 5.2 | 3.2 | 0.0059 | 8.3 | | 5.1 | 0.0000 | 19 |
| Q14686 | NCOA6 | | 5.1 | 2.8 | 0.0060 | 8.0 | | 3.7 | 0.0000 | 12 |
| Q8TAQ2 | SMARCC2 | | 5.1 | 4.0 | 0.0000 | 7.9 | | 4.4 | 0.0000 | 10 |
| P35658 | NUP214 | | 5.1 | 3.7 | 0.0049 | 6.2 | | 3.7 | 0.0000 | 20 |
| Q09472 | EP300 | | 5.1 | 4.3 | 0.0000 | 8.7 | | 4.3 | 0.0000 | 12 |
| O94916 | NFAT5 | | 5.1 | 3.4 | 0.0061 | 7.1 | | 4.4 | 0.0000 | 6 |
| P09874 | PARP1 | | 5.1 | 4.6 | 0.0000 | 5.5 | | 2.7 | 0.0000 | 11 |
| Q8TF68 | ZNF384 | | 5.0 | 3.9 | 0.0028 | 6.4 | | 3.4 | 0.0000 | 3 |
| Q6FI13;P0C0S8;Q99878;Q9BTM1;Q16777;Q93077;Q7L7L0;P04908;Q96KK5;P20671;P16104;Q8IUE6;Q71UI9;P0C0S5;Q96QV6 | HIST2H2AA3;HIST1H2AG;HIST1H2AJ;H2AFJ;HIST2H2AC;HIST1H2AC;HIST3H2A;HIST1H2AB;HIST1H2AH;HIST1H2AD;H2AFX;HIST2H2AB;H2AFV;H2AFZ;HIST1H2AA | | 5.0 | 1.3 | 0.0838 | 0.6 | | 0.1 | 0.9067 | 2 |
| Q6MZP7 | LIN54 | | 4.9 | 4.0 | 0.0000 | 8.2 | | 5.1 | 0.0000 | 11 |
| Q6KC79 | NIPBL | | 4.9 | 4.1 | 0.0000 | 7.3 | | 5.5 | 0.0000 | 18 |
| Q96RK0 | CIC | | 4.9 | 3.2 | 0.0061 | 8.0 | | 5.2 | 0.0000 | 15 |
| Q15723 | ELF2 | | 4.8 | 2.8 | 0.0062 | 6.5 | | 2.7 | 0.0000 | 8 |
| P17275 | JUNB | | 4.8 | 4.9 | 0.0000 | 6.1 | | 4.5 | 0.0000 | 8 |
| P54253 | ATXN1 | | 4.8 | 2.6 | 0.0077 | 7.4 | | 3.1 | 0.0000 | 9 |
| Q6STE5 | SMARCD3 | | 4.8 | 4.9 | 0.0000 | 6.1 | | 3.9 | 0.0000 | 11 |
| Q8IZL2 | MAML2 | | 4.8 | 3.9 | 0.0030 | 8.7 | | 5.5 | 0.0000 | 18 |
| Q9H4W6 | EBF3 | | 4.8 | 2.7 | 0.0055 | 8.2 | | 3.5 | 0.0000 | 3 |
| Q96QD5 | DEPDC7 | | 4.7 | 3.3 | 0.0052 | 7.8 | | 3.5 | 0.0000 | 7 |
| Q2TAL8 | QRICH1 | | 4.7 | 2.6 | 0.0078 | 7.8 | | 3.4 | 0.0000 | 7 |
| Q96BD5 | PHF21A | | 4.7 | 1.6 | 0.0478 | 7.0 | | 2.1 | 0.0006 | 10 |
| Q14814 | MEF2D | | 4.7 | 3.4 | 0.0058 | 8.1 | | 4.5 | 0.0000 | 9 |
| O00712 | NFIB | | 4.7 | 2.1 | 0.0211 | 9.5 | | 3.8 | 0.0000 | 5 |
| Q9UPN9 | TRIM33 | | 4.7 | 2.8 | 0.0056 | 6.8 | | 3.3 | 0.0000 | 13 |
| Q969G3 | SMARCE1 | | 4.6 | 2.9 | 0.0065 | 9.5 | | 5.3 | 0.0000 | 9 |
| P48552 | NRIP1 | | 4.6 | 4.6 | 0.0000 | 8.2 | | 5.5 | 0.0000 | 16 |
| P62805 | HIST1H4A | | 4.6 | 1.7 | 0.0383 | 0.9 | | 0.6 | 0.0790 | 3 |
| O15550;O14607 | KDM6A | | 4.6 | 3.2 | 0.0060 | 8.3 | | 3.9 | 0.0000 | 16 |
| Q92754 | TFAP2C | | 4.6 | 3.6 | 0.0042 | 7.9 | | 4.2 | 0.0000 | 3 |
| O00268 | TAF4 | | 4.6 | 2.2 | 0.0156 | 8.7 | | 4.3 | 0.0000 | 12 |
| Q93074 | MED12 | | 4.5 | 2.4 | 0.0120 | 8.3 | | 3.5 | 0.0000 | 20 |
| Q6PJG2 | ELMSAN1 | | 4.5 | 2.6 | 0.0075 | 7.4 | | 3.3 | 0.0000 | 13 |
| Q9HAH7 | FBRS | | 4.5 | 3.9 | 0.0026 | 9.2 | | 4.7 | 0.0000 | 8 |
| P38398 | BRCA1 | | 4.5 | 2.3 | 0.0134 | 5.9 | | 2.9 | 0.0000 | 13 |
| Q15596 | NCOA2 | | 4.5 | 2.4 | 0.0124 | 8.2 | | 3.4 | 0.0000 | 16 |
| Q8WXI9 | GATAD2B | | 4.5 | 1.7 | 0.0372 | 6.0 | | 2.1 | 0.0008 | 9 |
| P27540 | ARNT | | 4.4 | 3.9 | 0.0029 | 8.5 | | 4.4 | 0.0000 | 8 |
| Q9H1B7 | IRF2BPL | | 4.4 | 2.5 | 0.0087 | 7.7 | | 3.9 | 0.0000 | 9 |
| Q96L91 | EP400 | | 4.4 | 4.0 | 0.0000 | 8.0 | | 4.4 | 0.0000 | 28 |
| Q9P2D1 | CHD7 | | 4.3 | 3.9 | 0.0027 | 6.8 | | 5.0 | 0.0000 | 37 |
| O94992 | HEXIM1 | | 4.3 | 2.4 | 0.0122 | 6.2 | | 3.7 | 0.0000 | 3 |
| Q12830 | BPTF | | 4.3 | 4.2 | 0.0000 | 6.3 | | 4.3 | 0.0000 | 15 |
| P10244 | MYBL2 | | 4.3 | 2.1 | 0.0211 | 6.9 | | 2.7 | 0.0000 | 10 |
| Q9NZM4 | GLTSCR1 | | 4.3 | 4.1 | 0.0000 | 7.4 | | 4.6 | 0.0000 | 14 |
| Q9ULK4 | MED23 | | 4.3 | 2.6 | 0.0081 | 5.6 | | 3.0 | 0.0000 | 21 |
| O95391 | SLU7 | | 4.2 | 3.7 | 0.0050 | 6.8 | | 4.6 | 0.0000 | 5 |
| P39880 | CUX1 | | 4.2 | 3.5 | 0.0039 | 6.1 | | 4.0 | 0.0000 | 16 |
| Q9NSC2 | SALL1 | | 4.2 | 3.1 | 0.0055 | 8.0 | | 4.7 | 0.0000 | 11 |
| O75362 | ZNF217 | | 4.2 | 3.6 | 0.0044 | 6.4 | | 4.3 | 0.0000 | 17 |
| Q5T5X7 | BEND3 | | 4.2 | 3.1 | 0.0067 | 6.5 | | 3.9 | 0.0000 | 19 |
| Q03164 | KMT2A | | 4.2 | 3.4 | 0.0060 | 7.7 | | 4.2 | 0.0000 | 20 |
| Q14135 | VGLL4 | | 4.2 | 4.0 | 0.0000 | 6.9 | | 5.8 | 0.0000 | 4 |
| Q12800 | TFCP2 | | 4.2 | 2.4 | 0.0138 | 6.8 | | 5.5 | 0.0000 | 4 |
| Q6ZW49 | PAXIP1 | | 4.2 | 2.2 | 0.0141 | 7.7 | | 3.4 | 0.0000 | 7 |
| Q9UHV9 | PFDN2 | | 4.1 | 2.8 | 0.0061 | 8.5 | | 4.2 | 0.0000 | 2 |
| Q9UMZ2 | SYNRG | | 4.1 | 2.8 | 0.0063 | 7.1 | | 4.0 | 0.0000 | 18 |
| Q96DT7 | ZBTB10 | | 4.1 | 2.5 | 0.0085 | 6.4 | | 4.2 | 0.0000 | 7 |
| Q12789 | GTF3C1 | | 4.1 | 3.2 | 0.0048 | 5.8 | | 3.8 | 0.0000 | 13 |
| Q8NG31 | CASC5 | | 4.1 | 2.4 | 0.0137 | 6.5 | | 3.7 | 0.0000 | 22 |
| P14859 | POU2F1 | | 4.1 | 3.3 | 0.0051 | 7.4 | | 4.2 | 0.0000 | 8 |
| Q9Y230 | RUVBL2 | | 4.1 | 3.1 | 0.0058 | 5.5 | | 3.6 | 0.0000 | 5 |
| P18583 | SON | | 4.0 | 2.2 | 0.0170 | 5.3 | | 2.7 | 0.0000 | 11 |
| P19793 | RXRA | | 4.0 | 2.7 | 0.0055 | 6.9 | | 5.0 | 0.0000 | 8 |
| P33778;Q99880;Q93079;Q99877;Q16778;P62807;P58876;O60814;Q5QNW6;P23527;Q99879;Q8N257;P57053;P06899;Q96A08;A0A2R8Y619 | HIST1H2BB;HIST1H2BL;HIST1H2BH;HIST1H2BN;HIST2H2BE;HIST1H2BC;HIST1H2BD;HIST1H2BK;HIST2H2BF;HIST1H2BO;HIST1H2BM;HIST3H2BB;H2BFS;HIST1H2BJ;HIST1H2BA | | 4.0 | 0.8 | 0.2192 | 3.4 | | 0.6 | 0.0577 | 2 |
| Q96F86 | EDC3 | | 4.0 | 2.1 | 0.0208 | 6.2 | | 2.6 | 0.0000 | 4 |
| Q9P2N5 | RBM27 | | 4.0 | 1.9 | 0.0267 | 6.3 | | 2.6 | 0.0000 | 9 |
| Q9HAW4 | CLSPN | | 4.0 | 2.3 | 0.0138 | 5.3 | | 6.7 | 0.0000 | 7 |
| Q14004 | CDK13 | | 4.0 | 2.1 | 0.0190 | 7.4 | | 3.0 | 0.0000 | 12 |
| Q96RE7 | NACC1 | | 4.0 | 1.2 | 0.0942 | 7.6 | | 3.1 | 0.0000 | 7 |
| P08651 | NFIC | | 3.9 | 3.2 | 0.0062 | 5.8 | | 4.0 | 0.0000 | 3 |
| Q14938 | NFIX | | 3.9 | 3.6 | 0.0047 | 6.8 | | 4.1 | 0.0000 | 9 |
| P33240 | CSTF2 | | 3.9 | 2.3 | 0.0129 | 5.0 | | 2.6 | 0.0000 | 2 |
| P21333 | FLNA | | 3.9 | 1.8 | 0.0333 | 5.3 | | 2.2 | 0.0005 | 41 |
| P40763 | STAT3 | | 3.9 | 4.1 | 0.0000 | 7.8 | | 4.7 | 0.0000 | 11 |
| Q29RF7 | PDS5A | | 3.8 | 2.1 | 0.0205 | 5.6 | | 2.7 | 0.0000 | 12 |
| O95071 | UBR5 | | 3.8 | 2.3 | 0.0128 | 7.0 | | 3.1 | 0.0000 | 18 |
| Q9UK61 | FAM208A | | 3.8 | 2.2 | 0.0141 | 5.7 | | 2.9 | 0.0000 | 12 |
| Q7Z3K3 | POGZ | | 3.8 | 1.3 | 0.0851 | 6.4 | | 2.0 | 0.0012 | 16 |
| Q92766 | RREB1 | | 3.8 | 2.9 | 0.0060 | 6.1 | | 3.7 | 0.0000 | 6 |
| Q9HCS4 | TCF7L1 | | 3.8 | 3.3 | 0.0053 | 5.5 | | 1.5 | 0.0038 | 3 |
| P54259 | ATN1 | | 3.8 | 2.1 | 0.0185 | 6.9 | | 3.1 | 0.0000 | 8 |
| Q92618 | ZNF516 | | 3.8 | 2.3 | 0.0142 | 6.1 | | 3.1 | 0.0000 | 8 |
| Q9HCK8 | CHD8 | | 3.8 | 2.5 | 0.0086 | 6.2 | | 3.8 | 0.0000 | 13 |
| Q9Y490 | TLN1 | | 3.8 | 2.3 | 0.0133 | 6.9 | | 3.6 | 0.0000 | 24 |
| P17535 | JUND | | 3.8 | 1.7 | 0.0374 | 6.4 | | 4.1 | 0.0000 | 8 |
| Q9P2R6 | RERE | | 3.7 | 2.1 | 0.0194 | 7.1 | | 3.0 | 0.0000 | 7 |
| Q15911 | ZFHX3 | | 3.7 | 2.4 | 0.0127 | 7.0 | | 3.6 | 0.0000 | 19 |
| Q96JK9 | MAML3 | | 3.6 | 3.1 | 0.0066 | 6.1 | | 3.6 | 0.0000 | 6 |
| Q8TD26 | CHD6 | | 3.6 | 2.6 | 0.0077 | 7.8 | | 4.3 | 0.0000 | 31 |
| P62316 | SNRPD2 | | 3.6 | 1.5 | 0.0576 | 6.3 | | 2.3 | 0.0003 | 3 |
| Q9NVM9 | ASUN | | 3.6 | 2.8 | 0.0057 | 5.9 | | 3.2 | 0.0000 | 5 |
| P52655 | GTF2A1 | | 3.6 | 1.0 | 0.1458 | 7.3 | | 3.6 | 0.0000 | 2 |
| P49750 | YLPM1 | | 3.6 | 2.6 | 0.0081 | 6.3 | | 3.2 | 0.0000 | 14 |
| Q03112 | MECOM | | 3.5 | 2.3 | 0.0144 | 8.5 | | 4.1 | 0.0000 | 16 |
| Q01543 | FLI1 | | 3.5 | 1.8 | 0.0370 | 9.0 | | 3.6 | 0.0000 | 10 |
| Q9Y263 | PLAA | | 3.5 | 3.7 | 0.0025 | 6.4 | | 4.4 | 0.0000 | 7 |
| Q5VUA4 | ZNF318 | | 3.5 | 2.9 | 0.0059 | 6.6 | | 3.8 | 0.0000 | 18 |
| Q7Z5K2 | WAPAL | | 3.5 | 3.0 | 0.0062 | 5.9 | | 3.5 | 0.0000 | 9 |
| Q8IZD2 | KMT2E | | 3.5 | 1.5 | 0.0524 | 7.0 | | 2.6 | 0.0000 | 9 |
| P13056 | NR2C1 | | 3.5 | 2.1 | 0.0188 | 6.0 | | 3.2 | 0.0000 | 5 |
| Q9H0E3 | SAP130 | | 3.4 | 2.1 | 0.0193 | 6.2 | | 3.3 | 0.0000 | 6 |
| P06576 | ATP5B | | 3.4 | 2.9 | 0.0058 | 0.1 | | 0.1 | 0.9322 | 4 |
| Q9H0C8 | ILKAP | | 3.4 | 1.3 | 0.0848 | 5.1 | | 2.1 | 0.0006 | 4 |
| P78318 | IGBP1 | | 3.4 | 1.4 | 0.0656 | 6.9 | | 2.6 | 0.0000 | 6 |
| Q8N684 | CPSF7 | | 3.4 | 2.9 | 0.0059 | 6.8 | | 3.8 | 0.0000 | 6 |
| Q96L73 | NSD1 | | 3.4 | 2.5 | 0.0087 | 6.7 | | 3.8 | 0.0000 | 12 |
| P49790 | NUP153 | | 3.4 | 2.8 | 0.0064 | 4.6 | | 3.4 | 0.0000 | 17 |
| Q92585 | MAML1 | | 3.4 | 2.3 | 0.0139 | 7.7 | | 3.6 | 0.0000 | 13 |
| Q9Y4C1 | KDM3A | | 3.3 | 3.0 | 0.0064 | 6.8 | | 3.9 | 0.0000 | 10 |
| Q9UHF7 | TRPS1 | | 3.3 | 2.8 | 0.0062 | 7.9 | | 4.7 | 0.0000 | 12 |
| P62081 | RPS7 | | 3.3 | 2.2 | 0.0169 | 2.6 | | 1.6 | 0.0026 | 2 |
| Q6Y7W6 | GIGYF2 | | 3.3 | 2.3 | 0.0133 | 6.8 | | 3.7 | 0.0000 | 11 |
| P16949;Q93045 | STMN1;STMN2 | | 3.3 | 3.3 | 0.0049 | 4.8 | | 3.5 | 0.0000 | 2 |
| Q14839 | CHD4 | | 3.3 | 2.9 | 0.0057 | 4.7 | | 3.2 | 0.0000 | 28 |
| Q6N021 | TET2 | | 3.3 | 2.2 | 0.0143 | 6.2 | | 3.2 | 0.0000 | 15 |
| P48643 | CCT5 | | 3.2 | 2.3 | 0.0143 | 5.2 | | 2.9 | 0.0000 | 6 |
| O75925 | PIAS1 | | 3.2 | 2.2 | 0.0142 | 6.0 | | 3.3 | 0.0000 | 5 |
| Q9BTC0 | DIDO1 | | 3.2 | 4.5 | 0.0000 | 4.8 | | 4.0 | 0.0000 | 28 |
| P85037 | FOXK1 | | 3.2 | 2.8 | 0.0058 | 5.9 | | 4.4 | 0.0000 | 7 |
| Q9UKD1 | GMEB2 | | 3.2 | 1.9 | 0.0277 | 6.0 | | 2.5 | 0.0000 | 5 |
| Q9H4I2 | ZHX3 | | 3.1 | 1.4 | 0.0646 | 6.2 | | 2.4 | 0.0000 | 10 |
| P52948 | NUP98 | | 3.1 | 1.9 | 0.0310 | 5.8 | | 4.2 | 0.0000 | 8 |
| Q6PJP8 | DCLRE1A | | 3.1 | 2.1 | 0.0187 | 4.4 | | 2.7 | 0.0000 | 11 |
| Q7LBC6 | KDM3B | | 3.1 | 2.7 | 0.0078 | 5.9 | | 3.5 | 0.0000 | 18 |
| Q13469 | NFATC2 | | 3.1 | 1.8 | 0.0329 | 6.0 | | 3.1 | 0.0000 | 9 |
| Q9NQW6 | ANLN | | 3.1 | 2.1 | 0.0210 | 6.4 | | 3.7 | 0.0000 | 10 |
| Q96Q89 | KIF20B | | 3.1 | 1.8 | 0.0358 | 4.9 | | 5.2 | 0.0000 | 11 |
| Q96PV6 | LENG8 | | 3.1 | 2.3 | 0.0138 | 6.3 | | 4.7 | 0.0000 | 5 |
| Q9Y520 | PRRC2C | | 3.0 | 2.0 | 0.0214 | 5.0 | | 3.0 | 0.0000 | 3 |
| P36578 | RPL4 | | 3.0 | 0.5 | 0.4079 | -1.1 | | 0.1 | 0.8823 | 3 |
| P35711 | SOX5 | | 3.0 | 3.3 | 0.0055 | 5.9 | | 4.2 | 0.0000 | 12 |
| P31153 | MAT2A | | 3.0 | 2.8 | 0.0056 | 4.1 | | 2.8 | 0.0000 | 3 |
| P49321 | NASP | | 3.0 | 1.5 | 0.0558 | 4.4 | | 2.0 | 0.0009 | 5 |
| P11021 | HSPA5 | | 3.0 | 1.1 | 0.1094 | 3.7 | | 1.6 | 0.0036 | 10 |
| P27816 | MAP4 | | 2.9 | 2.9 | 0.0055 | 5.2 | | 4.3 | 0.0000 | 17 |
| Q15459 | SF3A1 | | 2.9 | 1.9 | 0.0311 | 6.5 | | 3.0 | 0.0000 | 12 |
| Q96PN7 | TRERF1 | | 2.9 | 1.6 | 0.0431 | 6.7 | | 2.9 | 0.0000 | 7 |
| P52597 | HNRNPF | | 2.9 | 1.4 | 0.0645 | 4.6 | | 2.1 | 0.0006 | 4 |
| Q5VT06 | CEP350 | | 2.9 | 2.7 | 0.0062 | 5.2 | | 3.4 | 0.0000 | 11 |
| Q9Y266 | NUDC | | 2.9 | 2.3 | 0.0129 | 4.7 | | 3.4 | 0.0000 | 3 |
| Q96EA4 | SPDL1 | | 2.9 | 1.5 | 0.0557 | 6.8 | | 4.1 | 0.0000 | 11 |
| Q5SW79 | CEP170 | | 2.9 | 1.3 | 0.0733 | 4.3 | | 1.9 | 0.0013 | 9 |
| O95785 | WIZ | | 2.9 | 2.6 | 0.0076 | 4.9 | | 2.9 | 0.0000 | 11 |
| P54277 | PMS1 | | 2.8 | 3.0 | 0.0061 | 4.3 | | 3.4 | 0.0000 | 19 |
| P10809 | HSPD1 | | 2.8 | 1.0 | 0.1386 | 0.1 | | 0.0 | 0.9858 | 5 |
| Q9UEE9 | CFDP1 | | 2.8 | 1.1 | 0.1242 | 4.1 | | 1.4 | 0.0045 | 3 |
| Q9NW82 | WDR70 | | 2.8 | 1.4 | 0.0641 | 4.5 | | 2.1 | 0.0006 | 5 |
| Q92841 | DDX17 | | 2.8 | 2.7 | 0.0064 | 4.2 | | 3.4 | 0.0000 | 4 |
| P05412 | JUN | | 2.8 | 1.9 | 0.0256 | 7.2 | | 4.4 | 0.0000 | 5 |
| O60563 | CCNT1 | | 2.8 | 2.3 | 0.0134 | 4.4 | | 3.2 | 0.0000 | 4 |
| Q13506 | NAB1 | | 2.8 | 1.1 | 0.1127 | 4.3 | | 1.7 | 0.0026 | 4 |
| O94842 | TOX4 | | 2.8 | 1.7 | 0.0382 | 4.4 | | 2.5 | 0.0000 | 3 |
| P51610 | HCFC1 | | 2.8 | 2.4 | 0.0119 | 5.1 | | 3.2 | 0.0000 | 28 |
| Q96PK6 | RBM14 | | 2.8 | 1.1 | 0.1115 | 3.8 | | 1.5 | 0.0038 | 6 |
| P49411 | TUFM | | 2.8 | 0.6 | 0.3334 | 2.2 | | 0.4 | 0.1693 | 6 |
| Q16594 | TAF9 | | 2.7 | 0.9 | 0.1503 | 4.8 | | 1.6 | 0.0033 | 2 |
| Q9Y4B4 | RAD54L2 | | 2.7 | 1.6 | 0.0458 | 7.7 | | 3.9 | 0.0000 | 8 |
| Q9H790 | EXO5 | | 2.7 | 5.1 | 0.0000 | 6.6 | | 4.8 | 0.0000 | 7 |
| P02545 | LMNA | | 2.7 | 0.7 | 0.2425 | 4.0 | | 1.1 | 0.0084 | 11 |
| Q13495 | MAMLD1 | | 2.7 | 1.4 | 0.0668 | 6.3 | | 3.0 | 0.0000 | 6 |
| Q8NC51 | SERBP1 | | 2.7 | 1.0 | 0.1364 | 2.6 | | 0.6 | 0.0723 | 2 |
| P15408 | FOSL2 | | 2.7 | 1.9 | 0.0286 | 2.8 | | 1.6 | 0.0029 | 2 |
| Q12872 | SFSWAP | | 2.7 | 1.2 | 0.0890 | 5.7 | | 2.6 | 0.0000 | 6 |
| Q9BWU0 | SLC4A1AP | | 2.7 | 1.8 | 0.0369 | 4.6 | | 3.0 | 0.0000 | 8 |
| Q9NPI1 | BRD7 | | 2.7 | 2.6 | 0.0075 | 4.4 | | 3.1 | 0.0000 | 9 |
| Q9BPX3 | NCAPG | | 2.7 | 2.8 | 0.0059 | 4.9 | | 4.0 | 0.0000 | 4 |
| Q9H0L4 | CSTF2T | | 2.7 | 2.2 | 0.0138 | 5.1 | | 3.7 | 0.0000 | 3 |
| O14929 | HAT1 | | 2.7 | 2.4 | 0.0118 | 4.7 | | 3.3 | 0.0000 | 3 |
| Q13151 | HNRNPA0 | | 2.7 | 1.6 | 0.0468 | 2.9 | | 1.2 | 0.0071 | 3 |
| Q93009 | USP7 | | 2.7 | 2.5 | 0.0088 | 5.3 | | 3.3 | 0.0000 | 27 |
| Q6UX04 | CWC27 | | 2.6 | 0.8 | 0.1946 | 6.0 | | 3.9 | 0.0000 | 5 |
| Q5VT52 | RPRD2 | | 2.6 | 4.0 | 0.0000 | 4.4 | | 3.9 | 0.0000 | 28 |
| Q92973 | TNPO1 | | 2.6 | 0.6 | 0.3333 | 8.3 | | 3.4 | 0.0000 | 10 |
| Q04724;Q04725 | TLE1 | | 2.6 | 2.3 | 0.0135 | 8.0 | | 3.9 | 0.0000 | 4 |
| Q4LE39 | ARID4B | | 2.6 | 0.8 | 0.2157 | 4.7 | | 1.5 | 0.0042 | 8 |
| Q07021 | C1QBP | | 2.6 | 0.4 | 0.4863 | 4.4 | | 1.7 | 0.0024 | 5 |
| P04406 | GAPDH | | 2.6 | 1.9 | 0.0255 | 1.0 | | 1.2 | 0.0066 | 4 |
| O43175 | PHGDH | | 2.6 | 1.9 | 0.0260 | 2.8 | | 1.1 | 0.0082 | 3 |
| O43670 | ZNF207 | | 2.6 | 1.5 | 0.0613 | 4.5 | | 2.1 | 0.0006 | 2 |
| Q92769 | HDAC2 | | 2.6 | 1.4 | 0.0665 | 3.8 | | 1.9 | 0.0012 | 2 |
| Q53ET0 | CRTC2 | | 2.5 | 1.9 | 0.0278 | 8.3 | | 4.2 | 0.0000 | 10 |
| O60281 | ZNF292 | | 2.5 | 1.4 | 0.0681 | 5.8 | | 3.5 | 0.0000 | 16 |
| Q9UKJ3 | GPATCH8 | | 2.5 | 1.0 | 0.1363 | 8.0 | | 3.7 | 0.0000 | 5 |
| Q8IWZ8 | SUGP1 | | 2.5 | 1.9 | 0.0268 | 5.1 | | 2.8 | 0.0000 | 12 |
| O43809 | NUDT21 | | 2.5 | 1.7 | 0.0399 | 4.5 | | 2.7 | 0.0000 | 5 |
| P38646 | HSPA9 | | 2.5 | 1.8 | 0.0336 | 2.1 | | 1.8 | 0.0016 | 5 |
| Q14157 | UBAP2L | | 2.5 | 3.1 | 0.0057 | 5.8 | | 3.3 | 0.0000 | 10 |
| Q8N163 | CCAR2 | | 2.5 | 1.1 | 0.1196 | 4.7 | | 1.8 | 0.0016 | 15 |
| Q15699 | ALX1 | | 2.5 | 0.8 | 0.2093 | 5.5 | | 3.3 | 0.0000 | 4 |
| E9PAV3;Q13765 | NACA | | 2.4 | 2.3 | 0.0132 | 1.0 | | 0.3 | 0.3969 | 2 |
| Q5VWN6 | FAM208B | | 2.4 | 1.3 | 0.0719 | 4.5 | | 2.5 | 0.0000 | 10 |
| P15336 | ATF2 | | 2.4 | 1.6 | 0.0433 | 5.8 | | 3.6 | 0.0000 | 4 |
| Q9UPP1 | PHF8 | | 2.4 | 1.8 | 0.0333 | 4.1 | | 2.6 | 0.0000 | 5 |
| Q04637 | EIF4G1 | | 2.4 | 2.2 | 0.0147 | 5.8 | | 4.5 | 0.0000 | 3 |
| P78371 | CCT2 | | 2.4 | 1.2 | 0.0906 | 4.7 | | 3.0 | 0.0000 | 8 |
| Q92576 | PHF3 | | 2.4 | 1.5 | 0.0588 | 5.1 | | 2.5 | 0.0000 | 12 |
| Q2TBE0 | CWF19L2 | | 2.4 | 1.3 | 0.0842 | 6.1 | | 3.4 | 0.0000 | 7 |
| P14651;O43365;P31249 | HOXB3;HOXA3;HOXD3 | | 2.4 | 2.1 | 0.0192 | 6.2 | | 4.3 | 0.0000 | 2 |
| O95402 | MED26 | | 2.4 | 1.4 | 0.0685 | 5.4 | | 2.8 | 0.0000 | 6 |
| Q8IWI9 | MGA | | 2.4 | 3.0 | 0.0065 | 3.7 | | 3.9 | 0.0000 | 43 |
| Q8NEM7 | SUPT20H | | 2.4 | 1.7 | 0.0384 | 5.0 | | 3.3 | 0.0000 | 3 |
| Q8WVM7 | STAG1 | | 2.4 | 1.1 | 0.1186 | 7.6 | | 3.4 | 0.0000 | 9 |
| P49959 | MRE11A | | 2.4 | 3.3 | 0.0051 | 3.7 | | 3.4 | 0.0000 | 16 |
| P09211 | GSTP1 | | 2.3 | 1.4 | 0.0643 | 6.2 | | 2.9 | 0.0000 | 5 |
| Q9H6T3 | RPAP3 | | 2.3 | 1.8 | 0.0334 | 5.1 | | 2.9 | 0.0000 | 7 |
| P05387 | RPLP2 | | 2.3 | 0.9 | 0.1710 | 0.8 | | 0.4 | 0.1991 | 3 |
| P00338 | LDHA | | 2.3 | 1.8 | 0.0372 | 2.2 | | 1.4 | 0.0046 | 4 |
| P54132 | BLM | | 2.3 | 0.8 | 0.1899 | 5.3 | | 2.7 | 0.0000 | 5 |
| Q16543 | CDC37 | | 2.3 | 1.0 | 0.1431 | 3.4 | | 2.1 | 0.0006 | 2 |
| Q96AE4 | FUBP1 | | 2.3 | 1.1 | 0.1130 | 4.5 | | 1.9 | 0.0013 | 3 |
| Q6UN15 | FIP1L1 | | 2.3 | 0.8 | 0.1943 | 5.1 | | 1.8 | 0.0016 | 4 |
| Q92598 | HSPH1 | | 2.3 | 3.2 | 0.0048 | 4.7 | | 4.8 | 0.0000 | 11 |
| Q9UBB9 | TFIP11 | | 2.3 | 2.3 | 0.0141 | 5.6 | | 4.1 | 0.0000 | 7 |
| P23284 | PPIB | | 2.3 | 1.2 | 0.0864 | -1.2 | | 0.4 | 0.2877 | 2 |
| A0AVK6 | E2F8 | | 2.3 | 1.9 | 0.0254 | 4.7 | | 3.1 | 0.0000 | 7 |
| Q13118 | KLF10 | | 2.3 | 3.0 | 0.0064 | 4.9 | | 4.0 | 0.0000 | 3 |
| P14921 | ETS1 | | 2.3 | 2.2 | 0.0160 | 4.2 | | 3.2 | 0.0000 | 3 |
| O15164 | TRIM24 | | 2.3 | 2.3 | 0.0131 | 5.8 | | 3.5 | 0.0000 | 15 |
| P52756 | RBM5 | | 2.3 | 1.1 | 0.1086 | 4.6 | | 3.4 | 0.0000 | 4 |
| P49327 | FASN | | 2.2 | 2.0 | 0.0252 | 3.1 | | 2.8 | 0.0000 | 19 |
| P49848 | TAF6 | | 2.2 | 1.3 | 0.0807 | 7.3 | | 4.1 | 0.0000 | 10 |
| Q09666 | AHNAK | | 2.2 | 3.6 | 0.0043 | 3.1 | | 3.8 | 0.0000 | 122 |
| Q9NW64 | RBM22 | | 2.2 | 1.2 | 0.0925 | 6.3 | | 3.9 | 0.0000 | 3 |
| O14981 | BTAF1 | | 2.2 | 2.3 | 0.0137 | 5.8 | | 4.2 | 0.0000 | 12 |
| Q9UKN8 | GTF3C4 | | 2.2 | 1.6 | 0.0484 | 5.4 | | 2.6 | 0.0000 | 11 |
| Q9BX63 | BRIP1 | | 2.2 | 1.4 | 0.0656 | 5.5 | | 2.9 | 0.0000 | 10 |
| Q5VTR2 | RNF20 | | 2.2 | 0.8 | 0.1878 | 5.6 | | 3.1 | 0.0000 | 9 |
| P05166 | PCCB | | 2.2 | 1.1 | 0.1065 | 1.1 | | 0.5 | 0.0887 | 9 |
| Q7L2H7 | EIF3M | | 2.2 | 1.4 | 0.0644 | 3.7 | | 2.3 | 0.0005 | 2 |
| P61247 | RPS3A | | 2.1 | 1.1 | 0.1152 | 2.7 | | 1.4 | 0.0045 | 2 |
| Q8N1G2 | CMTR1 | | 2.1 | 1.3 | 0.0718 | 4.1 | | 2.2 | 0.0005 | 15 |
| Q15742 | NAB2 | | 2.1 | 2.8 | 0.0058 | 5.1 | | 4.2 | 0.0000 | 7 |
| Q6ZN30 | BNC2 | | 2.1 | 3.2 | 0.0063 | 4.2 | | 4.3 | 0.0000 | 3 |
| Q9BW85 | CCDC94 | | 2.1 | 1.5 | 0.0582 | 2.9 | | 2.3 | 0.0005 | 2 |
| P13051 | UNG | | 2.1 | 2.2 | 0.0143 | 4.6 | | 3.4 | 0.0000 | 4 |
| Q6P2C8 | MED27 | | 2.1 | 1.4 | 0.0643 | 5.9 | | 4.9 | 0.0000 | 4 |
| P17980 | PSMC3 | | 2.1 | 1.1 | 0.1083 | 4.4 | | 3.4 | 0.0000 | 7 |
| O00154 | ACOT7 | | 2.1 | 2.4 | 0.0110 | 3.4 | | 3.5 | 0.0000 | 3 |
| O14979 | HNRNPDL | | 2.0 | 1.0 | 0.1325 | 3.7 | | 3.1 | 0.0000 | 2 |
| O43395 | PRPF3 | | 2.0 | 1.7 | 0.0381 | 3.9 | | 2.6 | 0.0000 | 17 |
| Q9H2Z4 | NKX2-4 | | 2.0 | 1.1 | 0.1187 | 5.5 | | 2.5 | 0.0000 | 2 |
| O15056 | SYNJ2 | | 2.0 | 1.0 | 0.1471 | 6.8 | | 3.9 | 0.0000 | 12 |
| Q13573 | SNW1 | | 2.0 | 1.5 | 0.0605 | 4.3 | | 2.5 | 0.0000 | 8 |
| Q5H9F3 | BCORL1 | | 2.0 | 1.4 | 0.0647 | 7.3 | | 3.2 | 0.0000 | 10 |
| Q9H2P0 | ADNP | | 2.0 | 1.8 | 0.0373 | 3.5 | | 2.6 | 0.0000 | 16 |
| O95239 | KIF4A | | 2.0 | 2.4 | 0.0136 | 3.7 | | 2.9 | 0.0000 | 16 |
| P23528 | CFL1 | | 2.0 | 1.7 | 0.0384 | 2.2 | | 2.4 | 0.0000 | 3 |
| O75575 | CRCP | | 2.0 | 0.5 | 0.3535 | 4.4 | | 1.4 | 0.0051 | 3 |
| P50219 | MNX1 | | 2.0 | 0.7 | 0.2239 | 5.5 | | 3.4 | 0.0000 | 2 |
| Q8IXK0 | PHC2 | | 2.0 | 1.2 | 0.0959 | 6.0 | | 3.2 | 0.0000 | 6 |
| Q7L014 | DDX46 | | 1.9 | 2.4 | 0.0123 | 3.3 | | 3.0 | 0.0000 | 27 |
| O75582 | RPS6KA5 | | 1.9 | 0.8 | 0.1978 | 4.7 | | 3.0 | 0.0000 | 5 |
| Q14687 | GSE1 | | 1.9 | 0.6 | 0.3118 | 9.2 | | 4.7 | 0.0000 | 15 |
| A3KN83 | SBNO1 | | 1.9 | 2.0 | 0.0227 | 3.9 | | 3.1 | 0.0000 | 25 |
| O94913 | PCF11 | | 1.9 | 2.8 | 0.0061 | 5.6 | | 3.4 | 0.0000 | 10 |
| Q96JM3 | CHAMP1 | | 1.9 | 3.5 | 0.0040 | 4.1 | | 3.8 | 0.0000 | 11 |
| P50991 | CCT4 | | 1.9 | 1.5 | 0.0570 | 1.8 | | 1.3 | 0.0064 | 7 |
| Q13263 | TRIM28 | | 1.9 | 2.3 | 0.0134 | 3.3 | | 3.1 | 0.0000 | 17 |
| Q14320 | FAM50A | | 1.9 | 0.4 | 0.4495 | 4.2 | | 1.1 | 0.0097 | 3 |
| P25685 | DNAJB1 | | 1.9 | 1.0 | 0.1445 | 4.0 | | 4.0 | 0.0000 | 2 |
| Q9BTA9 | WAC | | 1.9 | 2.3 | 0.0130 | 2.9 | | 2.1 | 0.0006 | 2 |
| Q8WXF1 | PSPC1 | | 1.9 | 0.6 | 0.2920 | 4.6 | | 1.6 | 0.0026 | 3 |
| Q69YH5 | CDCA2 | | 1.9 | 1.3 | 0.0744 | 4.7 | | 3.0 | 0.0000 | 11 |
| O00213 | APBB1 | | 1.8 | 1.4 | 0.0654 | 4.8 | | 2.6 | 0.0000 | 5 |
| Q9NSE4 | IARS2 | | 1.8 | 1.2 | 0.0945 | 1.6 | | 0.7 | 0.0440 | 4 |
| P0DMV8;P0DMV9 | HSPA1A;HSPA1B | | 1.8 | 1.9 | 0.0276 | 4.0 | | 3.4 | 0.0000 | 6 |
| O00303 | EIF3F | | 1.8 | 2.7 | 0.0063 | 4.0 | | 3.0 | 0.0000 | 3 |
| P27694 | RPA1 | | 1.8 | 2.2 | 0.0138 | 3.6 | | 3.5 | 0.0000 | 13 |
| O43151 | TET3 | | 1.8 | 1.8 | 0.0334 | 7.5 | | 3.8 | 0.0000 | 11 |
| Q6P1J9 | CDC73 | | 1.8 | 1.3 | 0.0831 | 3.7 | | 2.5 | 0.0000 | 10 |
| Q13887 | KLF5 | | 1.8 | 0.8 | 0.1926 | 7.1 | | 3.4 | 0.0000 | 5 |
| Q6VMQ6 | ATF7IP | | 1.8 | 2.3 | 0.0141 | 4.9 | | 4.1 | 0.0000 | 7 |
| Q6PJT7 | ZC3H14 | | 1.8 | 2.0 | 0.0222 | 4.4 | | 3.3 | 0.0000 | 9 |
| P18615 | NELFE | | 1.8 | 1.3 | 0.0831 | 4.3 | | 2.7 | 0.0000 | 7 |
| Q5T1R4 | HIVEP3 | | 1.8 | 1.1 | 0.1232 | 7.0 | | 4.1 | 0.0000 | 9 |
| Q96S55 | WRNIP1 | | 1.8 | 2.5 | 0.0098 | 4.8 | | 4.4 | 0.0000 | 4 |
| P12956 | XRCC6 | | 1.8 | 1.4 | 0.0670 | 3.4 | | 2.4 | 0.0000 | 13 |
| P46100 | ATRX | | 1.7 | 1.4 | 0.0643 | 3.9 | | 2.3 | 0.0003 | 15 |
| Q9H1D9 | POLR3F | | 1.7 | 0.9 | 0.1528 | 3.2 | | 1.5 | 0.0038 | 5 |
| Q9UJA5 | TRMT6 | | 1.7 | 2.1 | 0.0206 | 4.9 | | 3.5 | 0.0000 | 3 |
| P78347 | GTF2I | | 1.7 | 2.8 | 0.0059 | 3.4 | | 4.1 | 0.0000 | 19 |
| Q76L83 | ASXL2 | | 1.7 | 1.6 | 0.0473 | 4.8 | | 3.6 | 0.0000 | 3 |
| Q02241 | KIF23 | | 1.7 | 1.5 | 0.0560 | 4.1 | | 2.9 | 0.0000 | 8 |
| O43474 | KLF4 | | 1.7 | 2.1 | 0.0174 | 5.2 | | 2.8 | 0.0000 | 3 |
| Q8N954 | GPATCH11 | | 1.7 | 1.1 | 0.1063 | 2.8 | | 2.2 | 0.0007 | 2 |
| P51531 | SMARCA2 | | 1.7 | 0.9 | 0.1555 | 6.4 | | 4.3 | 0.0000 | 9 |
| Q9ULJ6 | ZMIZ1 | | 1.7 | 1.8 | 0.0332 | 8.0 | | 4.2 | 0.0000 | 4 |
| Q14980 | NUMA1 | | 1.7 | 0.9 | 0.1834 | 4.9 | | 2.9 | 0.0000 | 7 |
| Q15293 | RCN1 | | 1.7 | 1.3 | 0.0752 | -1.1 | | 0.7 | 0.0412 | 2 |
| P20585 | MSH3 | | 1.7 | 1.7 | 0.0385 | 4.0 | | 3.1 | 0.0000 | 8 |
| P98175 | RBM10 | | 1.7 | 2.2 | 0.0137 | 3.7 | | 3.0 | 0.0000 | 10 |
| Q04760 | GLO1 | | 1.7 | 1.2 | 0.0898 | 3.5 | | 2.4 | 0.0000 | 2 |
| Q13435 | SF3B2 | | 1.7 | 2.4 | 0.0121 | 4.3 | | 4.0 | 0.0000 | 18 |
| Q96ST3 | SIN3A | | 1.7 | 0.8 | 0.2148 | 4.5 | | 2.7 | 0.0000 | 5 |
| O75179 | ANKRD17 | | 1.6 | 1.3 | 0.0835 | 3.0 | | 1.3 | 0.0059 | 4 |
| Q9ULR0 | ISY1 | | 1.6 | 1.2 | 0.1041 | 3.8 | | 2.2 | 0.0006 | 3 |
| O94906 | PRPF6 | | 1.6 | 3.1 | 0.0058 | 4.0 | | 5.0 | 0.0000 | 5 |
| O94776 | MTA2 | | 1.6 | 0.5 | 0.4370 | 4.8 | | 2.6 | 0.0000 | 3 |
| Q9NVI1 | FANCI | | 1.6 | 0.7 | 0.2283 | 3.2 | | 1.5 | 0.0039 | 8 |
| Q02413 | DSG1 | | 1.6 | 0.5 | 0.3522 | -1.0 | | 0.3 | 0.4327 | 3 |
| P15822 | HIVEP1 | | 1.6 | 0.4 | 0.4759 | 5.7 | | 3.4 | 0.0000 | 10 |
| Q92620 | DHX38 | | 1.6 | 2.9 | 0.0064 | 4.4 | | 4.1 | 0.0000 | 19 |
| O15119 | TBX3 | | 1.6 | 1.0 | 0.1475 | 8.3 | | 4.2 | 0.0000 | 3 |
| Q16576;Q09028 | RBBP7;RBBP4 | | 1.6 | 1.3 | 0.0830 | 3.7 | | 4.8 | 0.0000 | 4 |
| Q00613 | HSF1 | | 1.6 | 1.9 | 0.0269 | 4.7 | | 4.8 | 0.0000 | 2 |
| Q3T8J9 | GON4L | | 1.6 | 1.0 | 0.1436 | 4.4 | | 2.8 | 0.0000 | 5 |
| P08243 | ASNS | | 1.6 | 1.5 | 0.0611 | 2.2 | | 2.0 | 0.0009 | 2 |
| Q06830 | PRDX1 | | 1.5 | 1.2 | 0.0965 | -2.0 | | 1.1 | 0.0095 | 3 |
| Q92945 | KHSRP | | 1.5 | 2.0 | 0.0242 | 4.4 | | 3.0 | 0.0000 | 16 |
| P11142 | HSPA8 | | 1.5 | 2.3 | 0.0138 | 2.8 | | 3.1 | 0.0000 | 8 |
| Q92797 | SYMPK | | 1.5 | 2.7 | 0.0065 | 3.0 | | 3.6 | 0.0000 | 23 |
| Q99832 | CCT7 | | 1.5 | 1.5 | 0.0608 | 4.6 | | 3.1 | 0.0000 | 5 |
| Q8NFC6 | BOD1L1 | | 1.5 | 1.2 | 0.1032 | 3.3 | | 2.1 | 0.0006 | 22 |
| P61204;P84077;P84085 | ARF3;ARF1;ARF5 | | 1.5 | 1.0 | 0.1347 | 2.4 | | 1.9 | 0.0012 | 3 |
| P61758 | VBP1 | | 1.5 | 0.9 | 0.1587 | 9.0 | | 3.9 | 0.0000 | 4 |
| O43502 | RAD51C | | 1.5 | 1.7 | 0.0404 | 3.0 | | 1.7 | 0.0026 | 2 |
| P26599 | PTBP1 | | 1.5 | 0.6 | 0.3106 | 3.4 | | 2.1 | 0.0006 | 3 |
| P22234 | PAICS | | 1.5 | 1.0 | 0.1293 | 1.9 | | 0.9 | 0.0164 | 3 |
| Q8NHM5 | KDM2B | | 1.5 | 0.9 | 0.1796 | 3.7 | | 2.2 | 0.0005 | 2 |
| Q9BWT3 | PAPOLG | | 1.5 | 0.7 | 0.2287 | 3.8 | | 2.2 | 0.0006 | 6 |
| Q9UPN6 | SCAF8 | | 1.5 | 0.9 | 0.1599 | 5.3 | | 3.2 | 0.0000 | 3 |
| Q6NWY9 | PRPF40B | | 1.5 | 1.2 | 0.0885 | 7.1 | | 3.7 | 0.0000 | 6 |
| Q96GX5 | MASTL | | 1.5 | 1.8 | 0.0331 | 5.0 | | 4.2 | 0.0000 | 6 |
| Q9UBD5 | ORC3 | | 1.5 | 0.7 | 0.2627 | 4.5 | | 1.9 | 0.0014 | 6 |
| Q8IX12 | CCAR1 | | 1.5 | 1.0 | 0.1414 | 6.1 | | 3.5 | 0.0000 | 6 |
| Q9UKX7 | NUP50 | | 1.5 | 0.6 | 0.3070 | 3.8 | | 1.9 | 0.0013 | 5 |
| P09884 | POLA1 | | 1.5 | 2.1 | 0.0191 | 3.9 | | 3.9 | 0.0000 | 2 |
| Q9NTZ6 | RBM12 | | 1.5 | 2.2 | 0.0155 | 4.3 | | 3.7 | 0.0000 | 8 |
| Q13356 | PPIL2 | | 1.5 | 0.7 | 0.2688 | 4.8 | | 3.9 | 0.0000 | 2 |
| P57740 | NUP107 | | 1.5 | 2.0 | 0.0238 | 5.9 | | 7.3 | 0.0000 | 8 |
| P11940;Q13310 | PABPC1;PABPC4 | | 1.5 | 0.7 | 0.2794 | 4.3 | | 3.7 | 0.0000 | 4 |
| Q9C0J8 | WDR33 | | 1.4 | 1.0 | 0.1366 | 3.4 | | 1.9 | 0.0012 | 3 |
| Q13177 | PAK2 | | 1.4 | 0.9 | 0.1797 | 3.9 | | 3.3 | 0.0000 | 11 |
| Q8N3X6 | LCORL | | 1.4 | 1.0 | 0.1424 | 5.6 | | 3.3 | 0.0000 | 6 |
| Q00341 | HDLBP | | 1.4 | 1.0 | 0.1412 | 3.6 | | 2.1 | 0.0008 | 5 |
| Q9UPT8 | ZC3H4 | | 1.4 | 1.1 | 0.1248 | 5.5 | | 2.9 | 0.0000 | 6 |
| Q9Y265 | RUVBL1 | | 1.4 | 0.8 | 0.1947 | 3.2 | | 2.1 | 0.0008 | 4 |
| P49368 | CCT3 | | 1.4 | 0.5 | 0.3553 | 3.1 | | 1.2 | 0.0070 | 7 |
| Q17R98 | ZNF827 | | 1.4 | 0.7 | 0.2681 | 6.0 | | 3.5 | 0.0000 | 10 |
| Q92547 | TOPBP1 | | 1.4 | 1.3 | 0.0788 | 3.6 | | 2.9 | 0.0000 | 3 |
| P17544 | ATF7 | | 1.4 | 0.4 | 0.4980 | 4.8 | | 2.3 | 0.0005 | 5 |
| O15417 | TNRC18 | | 1.4 | 2.5 | 0.0099 | 7.6 | | 4.3 | 0.0000 | 20 |
| Q92733 | PRCC | | 1.4 | 1.1 | 0.1245 | 6.4 | | 3.7 | 0.0000 | 6 |
| Q01664 | TFAP4 | | 1.4 | 1.0 | 0.1365 | 7.5 | | 4.4 | 0.0000 | 5 |
| P37275 | ZEB1 | | 1.4 | 0.6 | 0.2920 | 6.2 | | 4.5 | 0.0000 | 5 |
| P62826 | RAN | | 1.4 | 0.6 | 0.2933 | 0.8 | | 0.3 | 0.4430 | 3 |
| Q12824 | SMARCB1 | | 1.4 | 0.9 | 0.1603 | 3.7 | | 2.2 | 0.0006 | 6 |
| Q92750 | TAF4B | | 1.4 | 1.0 | 0.1411 | 3.1 | | 1.7 | 0.0026 | 2 |
| P62195 | PSMC5 | | 1.4 | 1.0 | 0.1358 | 4.4 | | 3.1 | 0.0000 | 9 |
| Q9UPW0 | FOXJ3 | | 1.4 | 0.4 | 0.5317 | 4.8 | | 2.7 | 0.0000 | 2 |
| Q9BZJ0 | CRNKL1 | | 1.4 | 1.0 | 0.1383 | 2.9 | | 2.3 | 0.0002 | 3 |
| Q9NPG3 | UBN1 | | 1.4 | 1.3 | 0.0757 | 4.4 | | 4.0 | 0.0000 | 4 |
| Q86YA3 | ZGRF1 | | 1.4 | 0.4 | 0.5200 | 6.3 | | 3.1 | 0.0000 | 10 |
| Q6ZRI6 | C15orf39 | | 1.4 | 0.4 | 0.4898 | 7.3 | | 5.2 | 0.0000 | 12 |
| Q9UPQ9 | TNRC6B | | 1.4 | 1.1 | 0.1124 | 4.4 | | 2.9 | 0.0000 | 6 |
| Q8IYH5 | ZZZ3 | | 1.3 | 1.1 | 0.1082 | 2.9 | | 2.0 | 0.0009 | 4 |
| Q96C24 | SYTL4 | | 1.3 | 1.7 | 0.0401 | 3.4 | | 3.2 | 0.0000 | 8 |
| Q99666 | RGPD5 | | 1.3 | 0.4 | 0.4834 | 3.4 | | 2.1 | 0.0006 | 2 |
| Q13442 | PDAP1 | | 1.3 | 1.9 | 0.0259 | 5.5 | | 4.2 | 0.0000 | 2 |
| P25705 | ATP5A1 | | 1.3 | 1.3 | 0.0739 | 2.8 | | 3.2 | 0.0000 | 9 |
| Q07157 | TJP1 | | 1.3 | 0.9 | 0.1569 | 5.8 | | 2.7 | 0.0000 | 9 |
| Q9Y5Q8 | GTF3C5 | | 1.3 | 0.5 | 0.3777 | 3.3 | | 2.0 | 0.0012 | 3 |
| Q86XP3 | DDX42 | | 1.3 | 1.4 | 0.0667 | 3.4 | | 2.5 | 0.0000 | 12 |
| Q9UHL9 | GTF2IRD1 | | 1.3 | 1.2 | 0.0935 | 4.9 | | 2.9 | 0.0000 | 6 |
| P43694 | GATA4 | | 1.3 | 2.0 | 0.0223 | 5.2 | | 4.1 | 0.0000 | 3 |
| O75940 | SMNDC1 | | 1.3 | 0.6 | 0.2921 | 3.2 | | 2.4 | 0.0000 | 2 |
| Q9NX08 | COMMD8 | | 1.3 | 0.7 | 0.2592 | 6.1 | | 3.4 | 0.0000 | 5 |
| Q14966 | ZNF638 | | 1.3 | 0.8 | 0.2149 | 2.6 | | 2.1 | 0.0008 | 21 |
| P78527 | PRKDC | | 1.3 | 2.4 | 0.0139 | 2.6 | | 2.9 | 0.0000 | 29 |
| P21675;Q8IZX4 | TAF1;TAF1L | | 1.3 | 0.6 | 0.2925 | 5.4 | | 4.3 | 0.0000 | 9 |
| Q15020 | SART3 | | 1.3 | 2.0 | 0.0230 | 2.5 | | 3.4 | 0.0000 | 10 |
| O15266;O60902 | SHOX;SHOX2 | | 1.3 | 0.6 | 0.3442 | 5.8 | | 3.9 | 0.0000 | 3 |
| P29401 | TKT | | 1.3 | 1.4 | 0.0654 | 2.6 | | 2.5 | 0.0000 | 6 |
| P42166 | TMPO | | 1.3 | 1.9 | 0.0270 | 3.2 | | 2.7 | 0.0000 | 9 |
| Q96QC0 | PPP1R10 | | 1.3 | 1.2 | 0.0887 | 3.6 | | 3.0 | 0.0000 | 6 |
| P34932 | HSPA4 | | 1.3 | 1.0 | 0.1322 | 3.9 | | 3.0 | 0.0000 | 8 |
| P61289 | PSME3 | | 1.2 | 2.2 | 0.0139 | 4.9 | | 6.0 | 0.0000 | 5 |
| P04083 | ANXA1 | | 1.2 | 1.8 | 0.0328 | 2.5 | | 3.2 | 0.0000 | 12 |
| O95359;O75410 | TACC2;TACC1 | | 1.2 | 0.4 | 0.4840 | 3.8 | | 2.6 | 0.0000 | 2 |
| P29590 | PML | | 1.2 | 1.3 | 0.0714 | 2.0 | | 3.4 | 0.0000 | 2 |
| O14950;P19105 | MYL12B;MYL12A | | 1.2 | 0.5 | 0.3939 | -0.2 | | 0.2 | 0.8346 | 2 |
| P46821 | MAP1B | | 1.2 | 0.8 | 0.1902 | 5.7 | | 4.2 | 0.0000 | 12 |
| O00571;O15523 | DDX3X;DDX3Y | | 1.2 | 0.5 | 0.4080 | 3.2 | | 1.4 | 0.0046 | 8 |
| Q96PZ0 | PUS7 | | 1.2 | 0.6 | 0.3231 | 4.0 | | 3.1 | 0.0000 | 6 |
| Q15382 | RHEB | | 1.2 | 2.1 | 0.0214 | 4.1 | | 3.7 | 0.0000 | 2 |
| Q99633 | PRPF18 | | 1.2 | 2.6 | 0.0080 | 3.3 | | 2.5 | 0.0000 | 5 |
| Q96A49 | SYAP1 | | 1.2 | 0.6 | 0.3385 | 4.4 | | 2.1 | 0.0008 | 3 |
| P14866 | HNRNPL | | 1.2 | 0.3 | 0.5622 | 2.9 | | 0.9 | 0.0164 | 4 |
| Q92917 | GPKOW | | 1.2 | 0.6 | 0.3370 | 4.7 | | 2.7 | 0.0000 | 7 |
| Q13330 | MTA1 | | 1.2 | 0.5 | 0.3700 | 5.2 | | 2.4 | 0.0000 | 8 |
| Q9H3P2 | NELFA | | 1.2 | 0.5 | 0.3532 | 3.8 | | 1.9 | 0.0012 | 5 |
| Q9NVP2 | ASF1B | | 1.2 | 1.1 | 0.1112 | 3.7 | | 2.4 | 0.0000 | 5 |
| P15924 | DSP | | 1.2 | 0.3 | 0.6954 | -3.0 | | 0.7 | 0.0361 | 10 |
| Q9Y2W1 | THRAP3 | | 1.2 | 0.2 | 0.7193 | 1.7 | | 0.4 | 0.3390 | 6 |
| Q9P270 | SLAIN2 | | 1.2 | 0.8 | 0.1948 | 3.0 | | 1.7 | 0.0025 | 2 |
| P08238 | HSP90AB1 | | 1.2 | 1.0 | 0.1379 | 1.5 | | 1.2 | 0.0066 | 7 |
| Q86X53 | ERICH1 | | 1.2 | 0.7 | 0.2660 | 4.3 | | 2.8 | 0.0000 | 3 |
| O95983 | MBD3 | | 1.2 | 0.9 | 0.1573 | 4.3 | | 2.8 | 0.0000 | 2 |
| O15294 | OGT | | 1.2 | 0.9 | 0.1566 | 4.7 | | 3.4 | 0.0000 | 5 |
| Q7Z4H7 | HAUS6 | | 1.2 | 1.4 | 0.0641 | 5.0 | | 2.7 | 0.0000 | 6 |
| Q96ST2 | IWS1 | | 1.2 | 1.2 | 0.0898 | 2.9 | | 1.3 | 0.0064 | 2 |
| P52594 | AGFG1 | | 1.2 | 0.7 | 0.2486 | 8.1 | | 4.9 | 0.0000 | 6 |
| P38606 | ATP6V1A | | 1.1 | 0.5 | 0.3775 | 6.0 | | 4.6 | 0.0000 | 7 |
| Q86U70 | LDB1 | | 1.1 | 0.9 | 0.1642 | 7.6 | | 4.2 | 0.0000 | 4 |
| Q16630 | CPSF6 | | 1.1 | 0.7 | 0.2692 | 4.4 | | 2.2 | 0.0005 | 6 |
| Q96T58 | SPEN | | 1.1 | 1.1 | 0.1248 | 4.8 | | 3.0 | 0.0000 | 7 |
| Q86YP4 | GATAD2A | | 1.1 | 0.9 | 0.1567 | 4.9 | | 2.9 | 0.0000 | 5 |
| P23246 | SFPQ | | 1.1 | 0.9 | 0.1750 | 3.2 | | 1.6 | 0.0033 | 6 |
| Q9H910 | HN1L | | 1.1 | 2.9 | 0.0054 | 2.4 | | 2.8 | 0.0000 | 5 |
| Q6UB98 | ANKRD12 | | 1.1 | 0.7 | 0.2485 | 2.8 | | 2.3 | 0.0003 | 2 |
| P32519 | ELF1 | | 1.1 | 0.9 | 0.1691 | 6.9 | | 2.8 | 0.0000 | 5 |
| Q6PKG0 | LARP1 | | 1.1 | 0.9 | 0.1500 | 1.8 | | 1.3 | 0.0054 | 2 |
| Q8WWM7 | ATXN2L | | 1.1 | 0.8 | 0.2169 | 5.0 | | 3.5 | 0.0000 | 6 |
| P61011 | SRP54 | | 1.1 | 0.6 | 0.3433 | 2.9 | | 2.0 | 0.0012 | 4 |
| Q8IWX8 | CHERP | | 1.1 | 1.6 | 0.0500 | 3.6 | | 3.2 | 0.0000 | 8 |
| Q86T24 | ZBTB33 | | 1.1 | 0.5 | 0.3523 | 4.4 | | 3.4 | 0.0000 | 3 |
| Q9GZU8 | FAM192A | | 1.1 | 1.3 | 0.0837 | 2.9 | | 2.9 | 0.0000 | 2 |
| Q00403 | GTF2B | | 1.1 | 0.8 | 0.2153 | 3.3 | | 2.2 | 0.0007 | 4 |
| Q9HB71 | CACYBP | | 1.1 | 1.1 | 0.1086 | 3.3 | | 2.8 | 0.0000 | 3 |
| P46060 | RANGAP1 | | 1.1 | 0.9 | 0.1569 | 3.0 | | 2.1 | 0.0006 | 4 |
| Q86U06 | RBM23 | | 1.1 | 0.4 | 0.4500 | 3.1 | | 2.5 | 0.0000 | 3 |
| Q96RL1 | UIMC1 | | 1.0 | 0.2 | 0.7219 | 4.9 | | 2.3 | 0.0005 | 6 |
| Q92900 | UPF1 | | 1.0 | 1.2 | 0.0916 | 3.6 | | 2.8 | 0.0000 | 16 |
| Q9NXV6 | CDKN2AIP | | 1.0 | 0.8 | 0.1887 | 3.4 | | 2.1 | 0.0006 | 3 |
| P52701 | MSH6 | | 1.0 | 1.9 | 0.0271 | 3.7 | | 3.9 | 0.0000 | 13 |
| Q9H5V9 | CXorf56 | | 1.0 | 1.0 | 0.1321 | 1.8 | | 1.9 | 0.0014 | 2 |
| O43399 | TPD52L2 | | 1.0 | 0.7 | 0.2692 | 3.6 | | 3.2 | 0.0000 | 3 |
| P41162 | ETV3 | | 1.0 | 0.7 | 0.2694 | 4.5 | | 2.1 | 0.0008 | 2 |
| Q5SSJ5 | HP1BP3 | | 1.0 | 0.4 | 0.5138 | 4.7 | | 3.2 | 0.0000 | 2 |
| O95551 | TDP2 | | 1.0 | 1.6 | 0.0460 | 2.1 | | 3.8 | 0.0000 | 5 |
| P31689 | DNAJA1 | | 1.0 | 0.9 | 0.1710 | 3.0 | | 1.8 | 0.0021 | 2 |
| P51812 | RPS6KA3 | | 1.0 | 0.6 | 0.3259 | 5.5 | | 3.5 | 0.0000 | 5 |
| P39023 | RPL3 | | 1.0 | 0.6 | 0.3437 | -0.8 | | 0.2 | 0.8272 | 2 |
| P37802 | TAGLN2 | | 1.0 | 1.4 | 0.0642 | 3.5 | | 3.5 | 0.0000 | 5 |
| Q15545 | TAF7 | | 1.0 | 0.5 | 0.4111 | 4.2 | | 2.6 | 0.0000 | 2 |
| P12270 | TPR | | 1.0 | 0.5 | 0.4192 | 3.4 | | 1.9 | 0.0014 | 10 |
| O60231 | DHX16 | | 1.0 | 0.4 | 0.5303 | 4.4 | | 1.8 | 0.0016 | 8 |
| O15397 | IPO8 | | 1.0 | 0.5 | 0.4151 | 4.2 | | 1.8 | 0.0016 | 4 |
| Q9UL03 | INTS6 | | 1.0 | 0.9 | 0.1504 | 4.4 | | 3.4 | 0.0000 | 4 |
| Q9NRZ9 | HELLS | | 1.0 | 1.5 | 0.0552 | 2.2 | | 2.4 | 0.0000 | 23 |
| Q96M27 | PRRC1 | | 1.0 | 1.1 | 0.1182 | 4.4 | | 3.9 | 0.0000 | 2 |
| Q5T4S7 | UBR4 | | 1.0 | 0.5 | 0.4150 | 3.7 | | 1.8 | 0.0016 | 7 |
| Q9NPD8 | UBE2T | | 1.0 | 0.5 | 0.3441 | 5.8 | | 4.6 | 0.0000 | 3 |
| Q9HCS7 | XAB2 | | 1.0 | 0.9 | 0.1599 | 2.7 | | 2.2 | 0.0007 | 26 |
| Q9UHR5 | SAP30BP | | 1.0 | 0.4 | 0.4897 | 5.3 | | 2.9 | 0.0000 | 4 |
| P08758 | ANXA5 | | 1.0 | 0.6 | 0.2977 | 0.6 | | 0.3 | 0.3517 | 2 |
| Q14202 | ZMYM3 | | 1.0 | 0.9 | 0.1548 | 3.5 | | 2.5 | 0.0000 | 3 |
| Q8ND24 | RNF214 | | 1.0 | 1.6 | 0.0434 | 2.4 | | 2.4 | 0.0000 | 3 |
| Q96EV2 | RBM33 | | 1.0 | 0.7 | 0.2793 | 6.3 | | 3.2 | 0.0000 | 6 |
| Q9NXZ1 | SAGE1 | | 1.0 | 0.5 | 0.4293 | 4.7 | | 2.2 | 0.0007 | 6 |
| P63092;Q5JWF2 | GNAS | | 1.0 | 0.9 | 0.1753 | 0.0 | | 0.0 | 0.9998 | 2 |
| O00763 | ACACB | | 1.0 | 1.9 | 0.0257 | 0.1 | | 0.1 | 0.9212 | 43 |
| Q6UUV7 | CRTC3 | | 1.0 | 0.4 | 0.5208 | 8.2 | | 4.3 | 0.0000 | 9 |
| Q15365 | PCBP1 | | 1.0 | 1.2 | 0.0962 | 2.9 | | 2.9 | 0.0000 | 2 |
| Q9BY77 | POLDIP3 | | 0.9 | 1.0 | 0.1324 | 3.6 | | 3.0 | 0.0000 | 4 |
| Q14247 | CTTN | | 0.9 | 0.9 | 0.1711 | 4.6 | | 4.9 | 0.0000 | 4 |
| Q96RU2 | USP28 | | 0.9 | 0.6 | 0.3395 | 3.7 | | 1.9 | 0.0013 | 5 |
| P17987 | TCP1 | | 0.9 | 0.6 | 0.3439 | 3.5 | | 2.6 | 0.0000 | 7 |
| Q13085 | ACACA | | 0.9 | 1.4 | 0.0643 | 0.2 | | 0.2 | 0.7674 | 66 |
| Q6ZSZ6 | TSHZ1 | | 0.9 | 0.5 | 0.3511 | 4.9 | | 2.6 | 0.0000 | 9 |
| Q9UQ35 | SRRM2 | | 0.9 | 0.8 | 0.1961 | 3.6 | | 2.6 | 0.0000 | 8 |
| P06733 | ENO1 | | 0.9 | 0.7 | 0.2422 | 1.1 | | 1.2 | 0.0075 | 7 |
| Q9BYW2 | SETD2 | | 0.9 | 0.5 | 0.3783 | 3.2 | | 2.0 | 0.0012 | 5 |
| Q9P281 | BAHCC1 | | 0.9 | 1.0 | 0.1319 | 5.8 | | 4.9 | 0.0000 | 7 |
| Q9H8M2 | BRD9 | | 0.9 | 0.8 | 0.2167 | 3.2 | | 1.6 | 0.0037 | 2 |
| P46776 | RPL27A | | 0.9 | 0.2 | 0.7436 | 3.2 | | 0.9 | 0.0178 | 2 |
| Q96C19;Q9BUP0 | EFHD2;EFHD1 | | 0.9 | 0.4 | 0.4875 | 5.0 | | 2.4 | 0.0000 | 2 |
| P63244 | GNB2L1 | | 0.9 | 0.9 | 0.1536 | 1.1 | | 0.9 | 0.0202 | 2 |
| Q9H9J4 | USP42 | | 0.9 | 0.9 | 0.1639 | 3.0 | | 3.3 | 0.0000 | 8 |
| Q92826 | HOXB13 | | 0.9 | 1.2 | 0.0866 | 1.6 | | 0.6 | 0.0605 | 2 |
| Q15233 | NONO | | 0.9 | 1.1 | 0.1187 | 3.3 | | 2.6 | 0.0000 | 8 |
| Q9UHX1 | PUF60 | | 0.9 | 1.0 | 0.1323 | 3.4 | | 3.1 | 0.0000 | 12 |
| P20042 | EIF2S2 | | 0.9 | 0.3 | 0.7010 | 2.2 | | 0.9 | 0.0158 | 2 |
| Q96TA1 | FAM129B | | 0.9 | 1.1 | 0.1118 | 3.9 | | 2.8 | 0.0000 | 4 |
| P53618 | COPB1 | | 0.9 | 0.4 | 0.4700 | 1.4 | | 0.5 | 0.1079 | 4 |
| P23381 | WARS | | 0.9 | 0.4 | 0.4935 | 2.3 | | 1.5 | 0.0040 | 3 |
| Q15637 | SF1 | | 0.9 | 1.0 | 0.1283 | 4.7 | | 3.9 | 0.0000 | 9 |
| P32969 | RPL9 | | 0.9 | 1.8 | 0.0351 | 1.7 | | 3.6 | 0.0000 | 2 |
| O75132 | ZBED4 | | 0.9 | 0.4 | 0.4419 | 4.0 | | 2.7 | 0.0000 | 3 |
| O94762 | RECQL5 | | 0.9 | 0.4 | 0.4472 | 3.5 | | 1.8 | 0.0016 | 2 |
| Q14103 | HNRNPD | | 0.9 | 1.6 | 0.0465 | -0.3 | | 0.1 | 0.9311 | 3 |
| P37840 | SNCA | | 0.9 | 0.8 | 0.1898 | 2.2 | | 3.3 | 0.0000 | 2 |
| Q14676 | MDC1 | | 0.9 | 0.5 | 0.4162 | 3.1 | | 2.0 | 0.0012 | 4 |
| P41229 | KDM5C | | 0.9 | 0.8 | 0.2000 | 5.8 | | 4.2 | 0.0000 | 9 |
| O95433 | AHSA1 | | 0.9 | 0.5 | 0.4362 | 4.2 | | 1.9 | 0.0012 | 2 |
| Q969V6 | MKL1 | | 0.9 | 0.5 | 0.3807 | 3.4 | | 2.7 | 0.0000 | 2 |
| Q53EL6 | PDCD4 | | 0.9 | 0.6 | 0.3120 | 3.5 | | 2.9 | 0.0000 | 2 |
| Q3L8U1 | CHD9 | | 0.8 | 0.4 | 0.4396 | 5.6 | | 3.0 | 0.0000 | 13 |
| Q69YN2 | CWF19L1 | | 0.8 | 1.8 | 0.0332 | 4.8 | | 4.5 | 0.0000 | 3 |
| Q6IQ49 | SDE2 | | 0.8 | 0.7 | 0.2401 | 3.3 | | 2.5 | 0.0000 | 3 |
| P49023 | PXN | | 0.8 | 0.7 | 0.2310 | 4.0 | | 3.0 | 0.0000 | 3 |
| P17098 | ZNF8 | | 0.8 | 0.6 | 0.3048 | 3.0 | | 2.1 | 0.0006 | 2 |
| Q8N3C0 | ASCC3 | | 0.8 | 0.4 | 0.4874 | 6.0 | | 3.3 | 0.0000 | 13 |
| Q969R5 | L3MBTL2 | | 0.8 | 0.6 | 0.2831 | 4.3 | | 2.6 | 0.0000 | 4 |
| P28347 | TEAD1 | | 0.8 | 0.6 | 0.3410 | 7.9 | | 4.4 | 0.0000 | 5 |
| Q9UKK9 | NUDT5 | | 0.8 | 0.3 | 0.5882 | 4.9 | | 2.1 | 0.0006 | 5 |
| Q01105;P0DME0 | SET;SETSIP | | 0.8 | 0.3 | 0.6578 | 0.3 | | 0.2 | 0.7547 | 3 |
| P28715 | ERCC5 | | 0.8 | 1.5 | 0.0606 | 2.7 | | 3.1 | 0.0000 | 13 |
| Q9UM54 | MYO6 | | 0.8 | 0.3 | 0.6770 | 4.7 | | 2.0 | 0.0012 | 4 |
| Q6IBS0 | TWF2 | | 0.8 | 1.5 | 0.0606 | 2.4 | | 2.6 | 0.0000 | 6 |
| Q6NSZ9 | ZSCAN25 | | 0.8 | 0.5 | 0.4111 | 3.9 | | 2.1 | 0.0006 | 2 |
| P31948 | STIP1 | | 0.8 | 0.6 | 0.3069 | 3.4 | | 2.1 | 0.0006 | 3 |
| P50613 | CDK7 | | 0.8 | 0.4 | 0.4714 | 2.9 | | 1.5 | 0.0040 | 2 |
| O75152 | ZC3H11A | | 0.8 | 0.9 | 0.1765 | 3.1 | | 3.0 | 0.0000 | 8 |
| O60664 | PLIN3 | | 0.8 | 0.5 | 0.4088 | 3.4 | | 1.7 | 0.0025 | 5 |
| Q7KZ85 | SUPT6H | | 0.8 | 0.4 | 0.4713 | 2.3 | | 1.3 | 0.0056 | 3 |
| Q07020 | RPL18 | | 0.8 | 3.3 | 0.0050 | 0.2 | | 0.2 | 0.7617 | 2 |
| Q86Y91 | KIF18B | | 0.8 | 0.4 | 0.5237 | 4.7 | | 2.1 | 0.0006 | 3 |
| P24928 | POLR2A | | 0.8 | 1.2 | 0.0886 | 4.7 | | 4.3 | 0.0000 | 6 |
| Q14677 | CLINT1 | | 0.8 | 0.5 | 0.3793 | 4.0 | | 3.4 | 0.0000 | 2 |
| Q8N201 | INTS1 | | 0.8 | 0.6 | 0.3330 | 3.1 | | 1.8 | 0.0019 | 3 |
| P62937 | PPIA | | 0.8 | 1.7 | 0.0416 | 0.2 | | 0.4 | 0.3290 | 4 |
| Q6YHU6 | THADA | | 0.8 | 0.3 | 0.6605 | 6.6 | | 3.4 | 0.0000 | 17 |
| Q9P0W2 | HMG20B | | 0.8 | 0.8 | 0.2206 | 4.2 | | 2.4 | 0.0000 | 2 |
| O75533 | SF3B1 | | 0.7 | 1.3 | 0.0833 | 2.6 | | 2.8 | 0.0000 | 28 |
| Q15084 | PDIA6 | | 0.7 | 0.3 | 0.6858 | 0.9 | | 0.3 | 0.4710 | 6 |
| Q96FJ0 | STAMBPL1 | | 0.7 | 0.6 | 0.3110 | 3.8 | | 2.5 | 0.0000 | 6 |
| O95400 | CD2BP2 | | 0.7 | 0.6 | 0.3014 | 5.0 | | 4.2 | 0.0000 | 4 |
| Q99590 | SCAF11 | | 0.7 | 0.7 | 0.2412 | 5.0 | | 4.8 | 0.0000 | 6 |
| Q6IQ32 | ADNP2 | | 0.7 | 0.5 | 0.3526 | 3.4 | | 2.4 | 0.0000 | 3 |
| Q96I25 | RBM17 | | 0.7 | 0.5 | 0.4148 | 5.3 | | 2.9 | 0.0000 | 3 |
| Q9Y570 | PPME1 | | 0.7 | 0.2 | 0.7086 | 4.9 | | 2.5 | 0.0000 | 4 |
| Q9UIU6 | SIX4 | | 0.7 | 0.6 | 0.3259 | 7.3 | | 4.3 | 0.0000 | 6 |
| P49116 | NR2C2 | | 0.7 | 0.3 | 0.6212 | 5.1 | | 4.4 | 0.0000 | 5 |
| Q08945 | SSRP1 | | 0.7 | 1.0 | 0.1323 | 2.6 | | 3.8 | 0.0000 | 10 |
| P60842 | EIF4A1 | | 0.7 | 1.4 | 0.0658 | -0.1 | | 0.1 | 0.9494 | 6 |
| O43719 | HTATSF1 | | 0.7 | 1.1 | 0.1242 | 4.7 | | 3.2 | 0.0000 | 4 |
| Q7Z417 | NUFIP2 | | 0.7 | 0.6 | 0.3392 | 3.2 | | 2.1 | 0.0006 | 2 |
| Q14119 | VEZF1 | | 0.7 | 0.8 | 0.1979 | 3.8 | | 2.9 | 0.0000 | 3 |
| Q9BTE3 | MCMBP | | 0.7 | 0.9 | 0.1569 | 2.7 | | 2.9 | 0.0000 | 2 |
| Q14847 | LASP1 | | 0.7 | 0.3 | 0.6306 | 2.5 | | 1.7 | 0.0024 | 2 |
| Q9NYV4 | CDK12 | | 0.7 | 0.4 | 0.4508 | 5.8 | | 2.9 | 0.0000 | 5 |
| P14618 | PKM | | 0.7 | 0.9 | 0.1611 | 1.5 | | 1.7 | 0.0024 | 16 |
| Q9NVZ3 | NECAP2 | | 0.7 | 0.2 | 0.7064 | 5.7 | | 4.3 | 0.0000 | 4 |
| Q16666 | IFI16 | | 0.7 | 1.9 | 0.0312 | 0.6 | | 1.7 | 0.0026 | 29 |
| Q70CQ2 | USP34 | | 0.7 | 0.4 | 0.4902 | 4.6 | | 2.5 | 0.0000 | 27 |
| Q8IWZ3 | ANKHD1 | | 0.7 | 0.6 | 0.2981 | 3.2 | | 3.4 | 0.0000 | 3 |
| P10599 | TXN | | 0.7 | 1.7 | 0.0389 | 2.9 | | 3.4 | 0.0000 | 2 |
| Q6N043 | ZNF280D | | 0.7 | 0.3 | 0.6361 | 4.4 | | 2.0 | 0.0008 | 3 |
| P05023 | ATP1A1 | | 0.7 | 0.9 | 0.1690 | 2.6 | | 2.8 | 0.0000 | 3 |
| Q9UBC2 | EPS15L1 | | 0.7 | 0.3 | 0.5905 | 5.3 | | 4.6 | 0.0000 | 7 |
| P83731 | RPL24 | | 0.7 | 1.3 | 0.0741 | 0.9 | | 0.9 | 0.0161 | 2 |
| Q7Z3T8 | ZFYVE16 | | 0.7 | 0.4 | 0.4647 | 2.7 | | 1.2 | 0.0076 | 3 |
| Q9Y314 | NOSIP | | 0.6 | 0.6 | 0.3386 | 2.8 | | 2.0 | 0.0009 | 2 |
| Q01780 | EXOSC10 | | 0.6 | 1.0 | 0.1359 | 2.6 | | 2.5 | 0.0000 | 17 |
| Q9H410 | DSN1 | | 0.6 | 0.3 | 0.6796 | 4.5 | | 2.0 | 0.0009 | 3 |
| Q5T7W0 | ZNF618 | | 0.6 | 0.5 | 0.4137 | 2.9 | | 3.0 | 0.0000 | 3 |
| Q99496 | RNF2 | | 0.6 | 0.4 | 0.4484 | 4.4 | | 3.9 | 0.0000 | 3 |
| O75663 | TIPRL | | 0.6 | 0.4 | 0.4673 | 3.4 | | 2.6 | 0.0000 | 2 |
| Q5TKA1 | LIN9 | | 0.6 | 0.4 | 0.4869 | 7.4 | | 4.3 | 0.0000 | 8 |
| Q16649 | NFIL3 | | 0.6 | 0.8 | 0.1947 | 5.4 | | 3.9 | 0.0000 | 5 |
| P46063 | RECQL | | 0.6 | 0.2 | 0.7600 | 5.4 | | 3.8 | 0.0000 | 4 |
| Q9UK76 | HN1 | | 0.6 | 0.2 | 0.8215 | 4.5 | | 2.0 | 0.0012 | 4 |
| Q14151 | SAFB2 | | 0.6 | 0.3 | 0.6200 | 3.2 | | 1.9 | 0.0013 | 4 |
| P54821 | PRRX1 | | 0.6 | 0.8 | 0.2158 | 4.7 | | 5.1 | 0.0000 | 4 |
| P86791;P86790 | CCZ1;CCZ1B | | 0.6 | 0.2 | 0.7317 | 4.9 | | 2.9 | 0.0000 | 6 |
| P78344 | EIF4G2 | | 0.6 | 0.3 | 0.6689 | 2.9 | | 1.4 | 0.0043 | 7 |
| P05388;Q8NHW5 | RPLP0;RPLP0P6 | | 0.6 | 0.4 | 0.5549 | -1.4 | | 1.2 | 0.0066 | 2 |
| P62241 | RPS8 | | 0.6 | 0.2 | 0.7593 | 1.0 | | 0.5 | 0.1193 | 2 |
| Q9Y285 | FARSA | | 0.6 | 0.2 | 0.7009 | 4.9 | | 2.2 | 0.0005 | 4 |
| Q9ULM3 | YEATS2 | | 0.6 | 1.0 | 0.1325 | 4.6 | | 5.6 | 0.0000 | 4 |
| Q99504 | EYA3 | | 0.6 | 0.4 | 0.4387 | 6.3 | | 4.2 | 0.0000 | 5 |
| O15042 | U2SURP | | 0.6 | 0.6 | 0.3339 | 2.7 | | 2.3 | 0.0005 | 12 |
| Q00610 | CLTC | | 0.6 | 0.8 | 0.2068 | 1.3 | | 1.7 | 0.0024 | 13 |
| P53999 | SUB1 | | 0.6 | 0.1 | 0.8539 | 2.0 | | 1.0 | 0.0118 | 3 |
| P0DP25;P0DP24;P0DP23;P27482 | CALML3 | | 0.6 | 0.3 | 0.6651 | 2.0 | | 1.4 | 0.0051 | 2 |
| Q8NF64 | ZMIZ2 | | 0.6 | 0.4 | 0.5102 | 3.0 | | 1.9 | 0.0013 | 2 |
| P35268 | RPL22 | | 0.6 | 0.7 | 0.2411 | 0.3 | | 0.2 | 0.8237 | 2 |
| P15170 | GSPT1 | | 0.6 | 0.5 | 0.4005 | 4.0 | | 2.0 | 0.0012 | 3 |
| Q04726 | TLE3 | | 0.5 | 0.3 | 0.6608 | 3.8 | | 1.8 | 0.0016 | 2 |
| Q8WY36 | BBX | | 0.5 | 0.6 | 0.3116 | 3.7 | | 2.4 | 0.0000 | 5 |
| Q9Y244 | POMP | | 0.5 | 0.3 | 0.6785 | 5.3 | | 4.5 | 0.0000 | 3 |
| P35251 | RFC1 | | 0.5 | 0.2 | 0.7434 | 3.3 | | 3.0 | 0.0000 | 3 |
| Q9BWF3 | RBM4 | | 0.5 | 0.3 | 0.6562 | 3.5 | | 2.1 | 0.0006 | 2 |
| Q9BRR8 | GPATCH1 | | 0.5 | 0.5 | 0.4053 | 2.9 | | 2.3 | 0.0005 | 17 |
| O75391 | SPAG7 | | 0.5 | 0.7 | 0.2648 | 3.1 | | 3.3 | 0.0000 | 4 |
| P13010 | XRCC5 | | 0.5 | 0.3 | 0.6792 | 3.1 | | 1.8 | 0.0016 | 7 |
| P07737 | PFN1 | | 0.5 | 0.6 | 0.3361 | 1.3 | | 1.5 | 0.0038 | 4 |
| Q96RQ3 | MCCC1 | | 0.5 | 1.3 | 0.0706 | 0.1 | | 0.2 | 0.8344 | 26 |
| Q99460 | PSMD1 | | 0.5 | 0.4 | 0.5125 | 3.7 | | 3.2 | 0.0000 | 3 |
| Q14865 | ARID5B | | 0.5 | 0.1 | 0.8529 | 7.5 | | 4.7 | 0.0000 | 13 |
| Q15942 | ZYX | | 0.5 | 0.2 | 0.7200 | 2.5 | | 2.0 | 0.0009 | 2 |
| Q08211 | DHX9 | | 0.5 | 0.9 | 0.1577 | 2.2 | | 2.3 | 0.0005 | 11 |
| Q9NXU5 | ARL15 | | 0.5 | 0.6 | 0.2853 | 2.8 | | 2.8 | 0.0000 | 3 |
| P53396 | ACLY | | 0.5 | 0.2 | 0.7204 | 3.4 | | 3.3 | 0.0000 | 2 |
| P26358 | DNMT1 | | 0.5 | 0.5 | 0.4141 | 3.2 | | 2.2 | 0.0006 | 2 |
| O75150 | RNF40 | | 0.5 | 0.2 | 0.7587 | 2.3 | | 1.2 | 0.0066 | 2 |
| Q86U86 | PBRM1 | | 0.5 | 0.4 | 0.5207 | 4.7 | | 3.0 | 0.0000 | 6 |
| Q86U42 | PABPN1 | | 0.5 | 0.4 | 0.4413 | 2.9 | | 2.5 | 0.0000 | 2 |
| Q7Z589 | EMSY | | 0.5 | 0.3 | 0.7002 | 4.9 | | 2.9 | 0.0000 | 8 |
| Q9Y2G9 | SBNO2 | | 0.5 | 0.3 | 0.6517 | 4.6 | | 2.7 | 0.0000 | 5 |
| O75528 | TADA3 | | 0.5 | 0.2 | 0.8309 | 3.2 | | 2.1 | 0.0008 | 2 |
| P11216 | PYGB | | 0.5 | 0.5 | 0.3498 | 1.1 | | 1.3 | 0.0055 | 7 |
| O43290 | SART1 | | 0.5 | 0.8 | 0.1953 | 2.2 | | 2.6 | 0.0000 | 22 |
| O43823 | AKAP8 | | 0.5 | 0.3 | 0.6516 | 3.5 | | 2.8 | 0.0000 | 7 |
| Q14807 | KIF22 | | 0.5 | 0.7 | 0.2285 | 3.3 | | 3.1 | 0.0000 | 17 |
| P08195 | SLC3A2 | | 0.5 | 0.3 | 0.5995 | 1.7 | | 1.1 | 0.0087 | 2 |
| Q96I24 | FUBP3 | | 0.4 | 0.2 | 0.7125 | 3.8 | | 2.0 | 0.0012 | 6 |
| Q9BXF3 | CECR2 | | 0.4 | 0.2 | 0.7111 | 5.8 | | 3.0 | 0.0000 | 5 |
| O75534 | CSDE1 | | 0.4 | 0.2 | 0.7428 | 2.4 | | 2.3 | 0.0005 | 2 |
| P07355;A6NMY6 | ANXA2;ANXA2P2 | | 0.4 | 0.5 | 0.3792 | -0.1 | | 0.1 | 0.9158 | 9 |
| Q9BZ95 | WHSC1L1 | | 0.4 | 0.4 | 0.5269 | 3.1 | | 2.8 | 0.0000 | 3 |
| Q13416 | ORC2 | | 0.4 | 0.4 | 0.5031 | 5.0 | | 3.6 | 0.0000 | 5 |
| Q9NZI7 | UBP1 | | 0.4 | 0.5 | 0.3680 | 4.7 | | 3.8 | 0.0000 | 2 |
| O75928 | PIAS2 | | 0.4 | 0.3 | 0.6289 | 3.3 | | 2.4 | 0.0000 | 2 |
| P48382 | RFX5 | | 0.4 | 0.3 | 0.6787 | 4.9 | | 3.0 | 0.0000 | 6 |
| Q9NRY4 | ARHGAP35 | | 0.4 | 0.2 | 0.7451 | 3.1 | | 1.7 | 0.0026 | 3 |
| Q86UE8 | TLK2 | | 0.4 | 0.2 | 0.7883 | 3.5 | | 2.5 | 0.0000 | 2 |
| Q7Z6K3 | PTAR1 | | 0.4 | 0.8 | 0.2153 | 4.8 | | 3.7 | 0.0000 | 5 |
| O00267 | SUPT5H | | 0.4 | 0.5 | 0.4243 | 4.3 | | 4.8 | 0.0000 | 11 |
| O15054 | KDM6B | | 0.4 | 0.3 | 0.6767 | 6.5 | | 3.3 | 0.0000 | 14 |
| O75643 | SNRNP200 | | 0.4 | 0.4 | 0.5503 | 2.6 | | 2.2 | 0.0006 | 33 |
| P40261 | NNMT | | 0.4 | 0.1 | 0.8773 | 3.3 | | 3.0 | 0.0000 | 2 |
| Q5SXM2 | SNAPC4 | | 0.4 | 0.3 | 0.5688 | 4.9 | | 3.9 | 0.0000 | 4 |
| O94782 | USP1 | | 0.4 | 0.2 | 0.7970 | 2.3 | | 1.5 | 0.0042 | 2 |
| Q5VTE6 | ANGEL2 | | 0.4 | 0.2 | 0.7982 | 3.2 | | 1.5 | 0.0038 | 2 |
| O60502 | MGEA5 | | 0.4 | 0.1 | 0.8652 | 3.2 | | 1.8 | 0.0016 | 4 |
| Q6NS38 | ALKBH2 | | 0.4 | 1.1 | 0.1083 | 1.9 | | 1.3 | 0.0065 | 2 |
| Q75N03 | CBLL1 | | 0.4 | 0.2 | 0.7405 | 3.7 | | 2.9 | 0.0000 | 2 |
| Q9Y2L1 | DIS3 | | 0.4 | 0.5 | 0.3676 | 1.3 | | 1.5 | 0.0038 | 3 |
| Q9ULH7 | MKL2 | | 0.4 | 0.2 | 0.7306 | 6.0 | | 4.0 | 0.0000 | 8 |
| Q9Y2W2 | WBP11 | | 0.4 | 0.1 | 0.8433 | 5.3 | | 4.0 | 0.0000 | 3 |
| Q99471 | PFDN5 | | 0.4 | 0.1 | 0.8643 | 5.7 | | 2.2 | 0.0005 | 2 |
| O43172 | PRPF4 | | 0.4 | 0.2 | 0.7671 | 3.4 | | 3.3 | 0.0000 | 5 |
| Q6P2H3 | CEP85 | | 0.4 | 0.1 | 0.8647 | 4.3 | | 2.3 | 0.0005 | 5 |
| P31943 | HNRNPH1 | | 0.4 | 0.1 | 0.9114 | 3.4 | | 1.4 | 0.0043 | 3 |
| Q9UNF1 | MAGED2 | | 0.3 | 0.3 | 0.6212 | 2.5 | | 2.1 | 0.0006 | 3 |
| P37231 | PPARG | | 0.3 | 0.2 | 0.7785 | 5.3 | | 3.4 | 0.0000 | 4 |
| P07900 | HSP90AA1 | | 0.3 | 0.3 | 0.5668 | 1.3 | | 2.7 | 0.0000 | 4 |
| O43148 | RNMT | | 0.3 | 0.1 | 0.9378 | 4.2 | | 1.8 | 0.0016 | 3 |
| Q13148 | TARDBP | | 0.3 | 0.1 | 0.9423 | 4.3 | | 1.3 | 0.0060 | 3 |
| P49915 | GMPS | | 0.3 | 0.2 | 0.8030 | 3.6 | | 2.5 | 0.0000 | 2 |
| P31273;P17481;P13378 | HOXC8;HOXB8;HOXD8 | | 0.3 | 0.3 | 0.6774 | 4.6 | | 2.3 | 0.0005 | 2 |
| O14776 | TCERG1 | | 0.3 | 0.1 | 0.9120 | 4.4 | | 1.9 | 0.0013 | 7 |
| P50747 | HLCS | | 0.3 | 0.6 | 0.3415 | -0.6 | | 0.5 | 0.1542 | 6 |
| P53365 | ARFIP2 | | 0.3 | 0.3 | 0.6212 | 3.7 | | 2.9 | 0.0000 | 2 |
| Q5T3J3 | LRIF1 | | 0.3 | 0.2 | 0.7029 | 5.4 | | 3.1 | 0.0000 | 10 |
| P52272 | HNRNPM | | 0.3 | 0.3 | 0.5685 | 2.9 | | 3.1 | 0.0000 | 15 |
| Q9UM82 | SPATA2 | | 0.3 | 0.3 | 0.6204 | 3.4 | | 3.7 | 0.0000 | 3 |
| Q8IX01 | SUGP2 | | 0.3 | 0.5 | 0.3792 | 1.6 | | 3.1 | 0.0000 | 15 |
| Q8TCN5 | ZNF507 | | 0.3 | 0.1 | 0.8534 | 3.2 | | 3.7 | 0.0000 | 2 |
| Q15393 | SF3B3 | | 0.3 | 0.3 | 0.6524 | 2.5 | | 2.4 | 0.0000 | 18 |
| Q96H20 | SNF8 | | 0.3 | 0.1 | 0.8773 | 4.0 | | 1.8 | 0.0021 | 2 |
| Q14C86 | GAPVD1 | | 0.3 | 0.1 | 0.9128 | 2.9 | | 2.1 | 0.0006 | 2 |
| Q9H3S7 | PTPN23 | | 0.3 | 0.2 | 0.7895 | 2.6 | | 2.5 | 0.0000 | 2 |
| P05165 | PCCA | | 0.3 | 0.5 | 0.4241 | -0.4 | | 0.5 | 0.0935 | 25 |
| P23396 | RPS3 | | 0.3 | 0.4 | 0.5102 | 0.6 | | 0.5 | 0.1615 | 6 |
| Q13207 | TBX2 | | 0.3 | 0.3 | 0.6300 | 3.9 | | 3.6 | 0.0000 | 2 |
| Q9H869 | YY1AP1 | | 0.3 | 0.1 | 0.8868 | 4.0 | | 1.9 | 0.0013 | 2 |
| Q5JSZ5 | PRRC2B | | 0.3 | 0.1 | 0.8632 | 6.2 | | 5.1 | 0.0000 | 9 |
| O15212 | PFDN6 | | 0.3 | 0.3 | 0.6425 | 5.6 | | 3.1 | 0.0000 | 4 |
| Q14562 | DHX8 | | 0.3 | 0.5 | 0.4145 | 3.1 | | 4.6 | 0.0000 | 3 |
| P62424 | RPL7A | | 0.3 | 0.2 | 0.8172 | 3.0 | | 2.3 | 0.0005 | 3 |
| Q6PIW4 | FIGNL1 | | 0.3 | 0.2 | 0.7224 | 3.1 | | 3.0 | 0.0000 | 3 |
| Q14498 | RBM39 | | 0.3 | 0.2 | 0.7134 | 1.7 | | 1.6 | 0.0027 | 5 |
| O00429 | DNM1L | | 0.3 | 0.1 | 0.9118 | 3.1 | | 1.4 | 0.0046 | 3 |
| O00273 | DFFA | | 0.2 | 0.2 | 0.7482 | 4.1 | | 2.9 | 0.0000 | 3 |
| Q14974 | KPNB1 | | 0.2 | 0.2 | 0.7980 | 1.6 | | 1.8 | 0.0021 | 3 |
| P31629 | HIVEP2 | | 0.2 | 0.2 | 0.7813 | 3.9 | | 3.1 | 0.0000 | 4 |
| Q8NDV7 | TNRC6A | | 0.2 | 0.3 | 0.6807 | 2.2 | | 2.2 | 0.0006 | 3 |
| Q96JN0 | LCOR | | 0.2 | 0.2 | 0.7950 | 2.8 | | 1.5 | 0.0038 | 2 |
| P18124 | RPL7 | | 0.2 | 0.6 | 0.2888 | -1.0 | | 0.9 | 0.0152 | 2 |
| Q9UBF1 | MAGEC2 | | 0.2 | 0.1 | 0.9380 | 3.9 | | 1.4 | 0.0043 | 4 |
| Q96A19 | CCDC102A | | 0.2 | 0.1 | 0.8688 | 3.4 | | 2.0 | 0.0012 | 3 |
| Q96GN5 | CDCA7L | | 0.2 | 0.2 | 0.8013 | 1.8 | | 1.1 | 0.0086 | 2 |
| P13639 | EEF2 | | 0.2 | 0.2 | 0.8287 | 0.8 | | 0.5 | 0.0898 | 6 |
| Q12906 | ILF3 | | 0.2 | 0.1 | 0.8317 | 1.8 | | 1.2 | 0.0066 | 5 |
| Q13123 | IK | | 0.2 | 0.4 | 0.4948 | 2.2 | | 2.3 | 0.0005 | 8 |
| Q9ULU4 | ZMYND8 | | 0.2 | 0.1 | 0.8593 | 4.4 | | 3.8 | 0.0000 | 6 |
| Q92786 | PROX1 | | 0.2 | 0.1 | 0.9278 | 4.0 | | 2.1 | 0.0006 | 3 |
| Q96G25 | MED8 | | 0.2 | 0.1 | 0.9272 | 5.8 | | 2.5 | 0.0000 | 3 |
| Q9ULW0 | TPX2 | | 0.2 | 0.2 | 0.7942 | 2.1 | | 2.1 | 0.0006 | 8 |
| P29083 | GTF2E1 | | 0.2 | 0.3 | 0.6548 | 4.5 | | 4.2 | 0.0000 | 4 |
| Q02880 | TOP2B | | 0.2 | 0.1 | 0.9388 | 2.6 | | 1.4 | 0.0046 | 2 |
| Q9C0D4 | ZNF518B | | 0.2 | 0.1 | 0.9274 | 5.4 | | 2.6 | 0.0000 | 5 |
| Q9H9A7 | RMI1 | | 0.2 | 0.2 | 0.7539 | 3.2 | | 2.3 | 0.0005 | 4 |
| P43246 | MSH2 | | 0.2 | 0.1 | 0.9266 | 2.5 | | 1.9 | 0.0013 | 3 |
| Q92782 | DPF1 | | 0.2 | 0.1 | 0.8437 | 3.6 | | 2.5 | 0.0000 | 2 |
| P21127;Q9UQ88 | CDK11B;CDK11A | | 0.2 | 0.1 | 0.8538 | -0.5 | | 0.4 | 0.2385 | 5 |
| Q9UNS1 | TIMELESS | | 0.2 | 0.1 | 0.9413 | 4.4 | | 3.1 | 0.0000 | 6 |
| Q8IVD9 | NUDCD3 | | 0.2 | 0.1 | 0.9225 | 3.2 | | 2.0 | 0.0012 | 2 |
| O43683 | BUB1 | | 0.2 | 0.1 | 0.8596 | 4.4 | | 2.7 | 0.0000 | 4 |
| P62701 | RPS4X | | 0.2 | 0.0 | 0.9497 | 1.4 | | 0.6 | 0.0772 | 2 |
| Q6ZU65 | UBN2 | | 0.1 | 0.1 | 0.9230 | 4.4 | | 4.2 | 0.0000 | 2 |
| P14678;P63162 | SNRPB;SNRPN | | 0.1 | 0.1 | 0.9422 | 4.0 | | 3.5 | 0.0000 | 2 |
| Q9Y330 | ZBTB12 | | 0.1 | 0.1 | 0.9271 | 5.8 | | 3.2 | 0.0000 | 4 |
| Q92481;P05549 | TFAP2B;TFAP2A | | 0.1 | 0.0 | 0.9501 | 6.5 | | 2.9 | 0.0000 | 3 |
| Q8NEA6 | GLIS3 | | 0.1 | 0.1 | 0.9291 | 4.2 | | 2.1 | 0.0006 | 3 |
| P15880 | RPS2 | | 0.1 | 0.0 | 0.9676 | 2.1 | | 0.6 | 0.0824 | 3 |
| Q14671 | PUM1 | | 0.1 | 0.1 | 0.8964 | 3.2 | | 2.5 | 0.0000 | 2 |
| P11498 | PC | | 0.1 | 0.1 | 0.8534 | 0.1 | | 0.1 | 0.8486 | 42 |
| Q2NL82 | TSR1 | | 0.1 | 0.1 | 0.9405 | 2.8 | | 1.5 | 0.0038 | 4 |
| P18858 | LIG1 | | 0.1 | 0.1 | 0.9391 | 2.9 | | 1.9 | 0.0013 | 3 |
| Q9UFF9 | CNOT8 | | 0.1 | 0.1 | 0.9426 | 2.3 | | 2.3 | 0.0003 | 2 |
| O75420 | GIGYF1 | | 0.1 | 0.1 | 0.9287 | 3.5 | | 1.7 | 0.0024 | 3 |
| Q01196 | RUNX1 | | 0.1 | 0.1 | 0.9392 | 5.5 | | 2.9 | 0.0000 | 4 |
| Q8NDI1 | EHBP1 | | 0.1 | 0.1 | 0.9279 | 2.7 | | 1.7 | 0.0026 | 3 |
| Q8NEM2 | SHCBP1 | | 0.1 | 0.1 | 0.9405 | 3.0 | | 1.8 | 0.0018 | 4 |
| Q13283 | G3BP1 | | 0.1 | 0.1 | 0.9278 | 4.0 | | 3.1 | 0.0000 | 2 |
| O95235 | KIF20A | | 0.1 | 0.1 | 0.9418 | 2.5 | | 2.3 | 0.0005 | 2 |
| P31942 | HNRNPH3 | | 0.1 | 0.0 | 0.9504 | 3.1 | | 2.2 | 0.0006 | 2 |
| Q9ULD9 | ZNF608 | | 0.1 | 0.1 | 0.8824 | 7.9 | | 4.1 | 0.0000 | 8 |
| Q9UPW6 | SATB2 | | 0.1 | 0.1 | 0.9398 | 5.4 | | 4.5 | 0.0000 | 5 |
| Q96P70 | IPO9 | | 0.1 | 0.1 | 0.9333 | 4.4 | | 3.4 | 0.0000 | 4 |
| Q9Y4E8 | USP15 | | 0.1 | 0.0 | 0.9658 | 3.1 | | 1.6 | 0.0033 | 4 |
| Q6IN85 | SMEK1 | | 0.1 | 0.1 | 0.8868 | 4.9 | | 4.5 | 0.0000 | 4 |
| Q96MY1 | NOL4L | | 0.1 | 0.0 | 0.9648 | 3.0 | | 1.9 | 0.0012 | 2 |
| O94986 | CEP152 | | 0.1 | 0.0 | 0.9690 | 2.9 | | 1.9 | 0.0014 | 3 |
| Q15843 | NEDD8 | | 0.1 | 0.0 | 0.9655 | 1.6 | | 1.5 | 0.0038 | 2 |
| Q6IEG0 | SNRNP48 | | 0.1 | 0.0 | 0.9455 | 5.5 | | 3.5 | 0.0000 | 3 |
| P53621 | COPA | | 0.1 | 0.0 | 0.9595 | 1.0 | | 0.9 | 0.0185 | 4 |
| P13984 | GTF2F2 | | 0.1 | 0.1 | 0.9382 | 0.6 | | 0.7 | 0.0364 | 3 |
| P31274 | HOXC9 | | 0.1 | 0.0 | 0.9655 | 6.0 | | 3.1 | 0.0000 | 4 |
| Q13470 | TNK1 | | 0.0 | 0.0 | 0.9640 | 4.3 | | 3.0 | 0.0000 | 4 |
| Q9UKL0 | RCOR1 | | 0.0 | 0.0 | 0.9760 | 3.8 | | 1.7 | 0.0026 | 2 |
| O43143 | DHX15 | | 0.0 | 0.0 | 0.9503 | 2.3 | | 2.5 | 0.0000 | 11 |
| Q9Y5R5 | DMRT2 | | 0.0 | 0.0 | 0.9514 | 4.1 | | 2.4 | 0.0000 | 4 |
| P28749 | RBL1 | | 0.0 | 0.0 | 0.9699 | 2.3 | | 1.2 | 0.0075 | 2 |
| Q5T6F2 | UBAP2 | | 0.0 | 0.0 | 0.9580 | 3.8 | | 3.1 | 0.0000 | 3 |
| P0C7T5 | ATXN1L | | 0.0 | 0.0 | 0.9745 | 4.6 | | 3.1 | 0.0000 | 2 |
| Q9P215 | POGK | | 0.0 | 0.0 | 0.9738 | 4.9 | | 3.1 | 0.0000 | 4 |
| Q8N2W9 | PIAS4 | | 0.0 | 0.0 | 0.9798 | 3.7 | | 1.5 | 0.0038 | 3 |
| P26641 | EEF1G | | 0.0 | 0.0 | 0.9686 | 1.0 | | 0.9 | 0.0195 | 4 |
| P49916 | LIG3 | | 0.0 | 0.0 | 0.9847 | 2.0 | | 1.5 | 0.0038 | 4 |
| Q6UB99 | ANKRD11 | | 0.0 | 0.0 | 0.9768 | 4.0 | | 3.0 | 0.0000 | 5 |
| Q9Y5X3 | SNX5 | | 0.0 | 0.0 | 0.9851 | 3.6 | | 2.5 | 0.0000 | 4 |
| Q8WXX5 | DNAJC9 | | 0.0 | 0.0 | 0.9904 | 1.9 | | 1.5 | 0.0042 | 2 |
| O60264 | SMARCA5 | | 0.0 | 0.0 | 0.9968 | 3.6 | | 1.6 | 0.0031 | 4 |
| O75934 | BCAS2 | | 0.0 | 0.0 | 0.9979 | 4.5 | | 3.2 | 0.0000 | 3 |
| O00257 | CBX4 | | 0.0 | 0.0 | 0.9972 | 4.1 | | 3.8 | 0.0000 | 2 |
| Q8WWY3 | PRPF31 | | 0.0 | 0.0 | 0.9969 | 4.0 | | 1.9 | 0.0014 | 5 |
| Q86XN7 | PROSER1 | | 0.0 | 0.0 | 0.9963 | 3.3 | | 1.1 | 0.0077 | 3 |
| Q8WZA0 | LZIC | | 0.0 | 0.0 | 0.9849 | 7.0 | | 3.1 | 0.0000 | 2 |
| Q7Z3B3 | KANSL1 | | 0.0 | 0.0 | 0.9738 | 4.4 | | 3.8 | 0.0000 | 8 |
| Q9H814 | PHAX | | 0.0 | 0.0 | 0.9707 | -0.4 | | 0.2 | 0.8039 | 3 |
| Q07864 | POLE | | 0.0 | 0.0 | 0.9736 | 2.4 | | 3.3 | 0.0000 | 3 |
| Q9Y5B9 | SUPT16H | | 0.0 | 0.1 | 0.9124 | 1.3 | | 2.0 | 0.0012 | 8 |
| Q9NWK9 | ZNHIT6 | | 0.0 | 0.0 | 0.9611 | 4.2 | | 3.3 | 0.0000 | 4 |
| Q9NVC6 | MED17 | | -0.1 | 0.0 | 0.9713 | 1.9 | | 1.1 | 0.0082 | 2 |
| Q9UEY8 | ADD3 | | -0.1 | 0.0 | 0.9736 | 2.9 | | 1.9 | 0.0014 | 2 |
| Q96CW5 | TUBGCP3 | | -0.1 | 0.0 | 0.9558 | 8.1 | | 5.2 | 0.0000 | 14 |
| Q6P5Z2 | PKN3 | | -0.1 | 0.0 | 0.9632 | 2.2 | | 2.1 | 0.0006 | 2 |
| Q14204 | DYNC1H1 | | -0.1 | 0.1 | 0.9189 | 4.0 | | 5.5 | 0.0000 | 6 |
| P07195 | LDHB | | -0.1 | 0.0 | 0.9736 | 1.7 | | 2.2 | 0.0006 | 3 |
| Q8N4C8 | MINK1 | | -0.1 | 0.0 | 0.9441 | 2.3 | | 1.6 | 0.0026 | 2 |
| Q13363 | CTBP1 | | -0.1 | 0.1 | 0.9260 | 3.9 | | 3.7 | 0.0000 | 3 |
| Q8NI27 | THOC2 | | -0.1 | 0.0 | 0.9731 | 1.4 | | 0.4 | 0.2405 | 2 |
| Q9NQS7 | INCENP | | -0.1 | 0.1 | 0.9377 | 2.5 | | 1.8 | 0.0016 | 9 |
| O75821 | EIF3G | | -0.1 | 0.1 | 0.9080 | 2.9 | | 1.8 | 0.0016 | 2 |
| Q96HW7 | INTS4 | | -0.1 | 0.0 | 0.9450 | 6.4 | | 3.1 | 0.0000 | 10 |
| Q9C0C2 | TNKS1BP1 | | -0.1 | 0.1 | 0.9413 | 4.1 | | 2.3 | 0.0003 | 6 |
| P17844 | DDX5 | | -0.1 | 0.0 | 0.9501 | 1.9 | | 1.1 | 0.0088 | 9 |
| O75486 | SUPT3H | | -0.1 | 0.0 | 0.9497 | 3.0 | | 2.0 | 0.0009 | 2 |
| Q5MIZ7 | SMEK2 | | -0.1 | 0.1 | 0.9211 | 3.6 | | 4.5 | 0.0000 | 6 |
| Q96KR1 | ZFR | | -0.1 | 0.1 | 0.9367 | 3.2 | | 1.0 | 0.0134 | 5 |
| Q9Y5B6 | PAXBP1 | | -0.1 | 0.1 | 0.9215 | 1.3 | | 0.8 | 0.0247 | 2 |
| Q92560 | BAP1 | | -0.1 | 0.1 | 0.9405 | 3.0 | | 2.5 | 0.0000 | 4 |
| Q07666 | KHDRBS1 | | -0.1 | 0.1 | 0.9270 | 1.3 | | 0.7 | 0.0452 | 2 |
| Q96IZ5 | RBM41 | | -0.1 | 0.1 | 0.9285 | 4.4 | | 2.8 | 0.0000 | 5 |
| Q9BUJ2 | HNRNPUL1 | | -0.1 | 0.0 | 0.9685 | 5.7 | | 2.1 | 0.0008 | 5 |
| Q9HCJ3 | RAVER2 | | -0.1 | 0.1 | 0.8824 | 4.3 | | 3.4 | 0.0000 | 4 |
| A6NHR9 | SMCHD1 | | -0.1 | 0.4 | 0.4404 | 2.5 | | 2.8 | 0.0000 | 2 |
| P00558 | PGK1 | | -0.1 | 0.1 | 0.8468 | 0.5 | | 0.5 | 0.1382 | 9 |
| Q92540 | SMG7 | | -0.1 | 0.1 | 0.9300 | 5.2 | | 2.7 | 0.0000 | 8 |
| O75113 | N4BP1 | | -0.2 | 0.1 | 0.9419 | 3.3 | | 1.4 | 0.0046 | 3 |
| Q7Z5J4 | RAI1 | | -0.2 | 0.3 | 0.6947 | 3.9 | | 3.4 | 0.0000 | 4 |
| O14503 | BHLHE40 | | -0.2 | 0.2 | 0.7633 | 3.6 | | 2.0 | 0.0012 | 2 |
| Q96AY2 | EME1 | | -0.2 | 0.1 | 0.9431 | 3.8 | | 1.3 | 0.0056 | 2 |
| P33981 | TTK | | -0.2 | 0.1 | 0.9172 | 3.7 | | 2.9 | 0.0000 | 2 |
| P19447 | ERCC3 | | -0.2 | 0.1 | 0.8709 | 2.3 | | 1.9 | 0.0014 | 2 |
| Q96BP3 | PPWD1 | | -0.2 | 0.1 | 0.9095 | 2.3 | | 1.4 | 0.0045 | 2 |
| Q92879 | CELF1 | | -0.2 | 0.1 | 0.9237 | 6.2 | | 3.5 | 0.0000 | 2 |
| Q08050 | FOXM1 | | -0.2 | 0.1 | 0.9280 | 3.5 | | 2.5 | 0.0000 | 3 |
| P35269 | GTF2F1 | | -0.2 | 0.1 | 0.8444 | 1.7 | | 2.2 | 0.0007 | 7 |
| Q8N0X7 | SPG20 | | -0.2 | 0.1 | 0.9255 | 2.6 | | 1.3 | 0.0057 | 2 |
| P04792 | HSPB1 | | -0.2 | 0.1 | 0.9267 | 1.4 | | 1.0 | 0.0146 | 3 |
| Q9BUL5 | PHF23 | | -0.2 | 0.1 | 0.8826 | 4.0 | | 3.0 | 0.0000 | 3 |
| Q06546 | GABPA | | -0.2 | 0.2 | 0.7819 | 4.9 | | 3.5 | 0.0000 | 6 |
| Q00839 | HNRNPU | | -0.2 | 0.5 | 0.4154 | 0.2 | | 0.4 | 0.2263 | 5 |
| P55771;P15863 | PAX9;PAX1 | | -0.2 | 0.1 | 0.9265 | 5.6 | | 2.5 | 0.0000 | 5 |
| P43358;P43363;P43362 | MAGEA4;MAGEA10;MAGEA9 | | -0.2 | 0.0 | 0.9650 | 2.6 | | 0.8 | 0.0238 | 2 |
| Q9UQL6 | HDAC5 | | -0.2 | 0.2 | 0.8314 | 1.5 | | 0.6 | 0.0598 | 3 |
| Q99541 | PLIN2 | | -0.2 | 0.2 | 0.7184 | 3.8 | | 4.0 | 0.0000 | 4 |
| Q8WYP5 | AHCTF1 | | -0.2 | 0.2 | 0.7778 | 2.3 | | 1.2 | 0.0068 | 9 |
| P55081 | MFAP1 | | -0.2 | 0.2 | 0.7767 | 1.9 | | 2.1 | 0.0006 | 5 |
| P10243 | MYBL1 | | -0.2 | 0.3 | 0.6592 | 4.4 | | 2.7 | 0.0000 | 2 |
| Q92785 | DPF2 | | -0.2 | 0.1 | 0.8665 | 4.8 | | 2.4 | 0.0000 | 6 |
| O15090 | ZNF536 | | -0.3 | 0.2 | 0.8119 | 4.2 | | 2.8 | 0.0000 | 6 |
| A1X283 | SH3PXD2B | | -0.3 | 0.2 | 0.7821 | 1.4 | | 1.8 | 0.0016 | 2 |
| Q9Y3F4 | STRAP | | -0.3 | 0.2 | 0.7230 | 0.1 | | 0.1 | 0.9256 | 7 |
| Q16512 | PKN1 | | -0.3 | 0.2 | 0.7051 | 2.2 | | 3.2 | 0.0000 | 3 |
| P50990 | CCT8 | | -0.3 | 0.4 | 0.5034 | 3.8 | | 3.6 | 0.0000 | 16 |
| P42167 | TMPO | | -0.3 | 0.4 | 0.5451 | 1.4 | | 1.7 | 0.0024 | 4 |
| Q9Y5Z7 | HCFC2 | | -0.3 | 0.2 | 0.7843 | 1.8 | | 1.2 | 0.0070 | 2 |
| Q9HCM7 | FBRSL1 | | -0.3 | 0.1 | 0.9014 | 5.7 | | 2.2 | 0.0007 | 4 |
| P53539 | FOSB | | -0.3 | 0.3 | 0.6766 | 4.2 | | 2.8 | 0.0000 | 3 |
| Q8IZH2 | XRN1 | | -0.3 | 0.2 | 0.7977 | 3.7 | | 2.2 | 0.0006 | 8 |
| O14980 | XPO1 | | -0.3 | 0.2 | 0.7306 | 2.2 | | 2.7 | 0.0000 | 2 |
| Q9UN79 | SOX13 | | -0.3 | 0.3 | 0.6581 | 5.7 | | 3.2 | 0.0000 | 6 |
| P39748 | FEN1 | | -0.3 | 0.2 | 0.7748 | 2.8 | | 2.6 | 0.0000 | 2 |
| Q99613;B5ME19 | EIF3C;EIF3CL | | -0.3 | 0.1 | 0.9266 | 2.4 | | 0.8 | 0.0335 | 5 |
| Q9Y6X9 | MORC2 | | -0.3 | 0.9 | 0.1662 | 2.0 | | 2.4 | 0.0000 | 3 |
| P46781 | RPS9 | | -0.3 | 0.3 | 0.6253 | -1.1 | | 1.4 | 0.0049 | 2 |
| Q9HCM1 | KIAA1551 | | -0.4 | 0.2 | 0.8031 | 4.5 | | 2.2 | 0.0006 | 5 |
| Q9UIF8 | BAZ2B | | -0.4 | 0.2 | 0.7428 | 4.4 | | 3.0 | 0.0000 | 5 |
| O95677 | EYA4 | | -0.4 | 0.2 | 0.7533 | 4.0 | | 3.1 | 0.0000 | 2 |
| Q9BT92 | TCHP | | -0.4 | 0.3 | 0.6045 | 3.0 | | 2.5 | 0.0000 | 3 |
| P23588 | EIF4B | | -0.4 | 0.1 | 0.8330 | 2.3 | | 1.5 | 0.0042 | 2 |
| P08865;A0A8I5KQE6 | RPSA | | -0.4 | 0.1 | 0.8470 | 0.4 | | 0.2 | 0.7259 | 2 |
| Q9P2K3 | RCOR3 | | -0.4 | 0.3 | 0.6863 | 1.6 | | 1.8 | 0.0023 | 2 |
| Q8WUA2 | PPIL4 | | -0.4 | 0.4 | 0.5460 | 3.3 | | 3.9 | 0.0000 | 4 |
| Q8NCD3 | HJURP | | -0.4 | 0.3 | 0.6293 | 3.9 | | 3.2 | 0.0000 | 4 |
| P40227 | CCT6A | | -0.4 | 0.2 | 0.8281 | 2.1 | | 2.0 | 0.0012 | 5 |
| Q9H1A4 | ANAPC1 | | -0.4 | 0.3 | 0.6443 | 2.1 | | 2.0 | 0.0009 | 2 |
| Q9P2N6 | KANSL3 | | -0.4 | 0.2 | 0.8128 | 5.1 | | 2.5 | 0.0000 | 7 |
| Q9NXR8 | ING3 | | -0.4 | 0.2 | 0.8191 | 3.2 | | 1.5 | 0.0038 | 2 |
| O14737 | PDCD5 | | -0.4 | 0.2 | 0.7951 | 2.6 | | 1.8 | 0.0021 | 3 |
| P35611 | ADD1 | | -0.4 | 0.3 | 0.6800 | 2.3 | | 1.8 | 0.0016 | 2 |
| Q6PCB5 | RSBN1L | | -0.4 | 0.2 | 0.8011 | 0.8 | | 0.3 | 0.3679 | 3 |
| O00148;Q13838 | DDX39A;DDX39B | | -0.4 | 0.2 | 0.8124 | 3.0 | | 1.7 | 0.0025 | 2 |
| Q14669 | TRIP12 | | -0.4 | 0.4 | 0.4945 | 1.4 | | 1.4 | 0.0049 | 18 |
| Q02086 | SP2 | | -0.4 | 0.3 | 0.6782 | 4.4 | | 2.9 | 0.0000 | 2 |
| Q9NR30 | DDX21 | | -0.4 | 0.2 | 0.8184 | -0.6 | | 0.2 | 0.7923 | 2 |
| Q8TEK3 | DOT1L | | -0.4 | 0.4 | 0.4528 | 2.5 | | 2.5 | 0.0000 | 2 |
| Q8WXE1 | ATRIP | | -0.4 | 0.2 | 0.7095 | 3.4 | | 2.1 | 0.0008 | 3 |
| Q8NCF5 | NFATC2IP | | -0.4 | 0.2 | 0.7308 | 4.4 | | 3.4 | 0.0000 | 5 |
| Q14207 | NPAT | | -0.4 | 0.1 | 0.8533 | 3.5 | | 1.5 | 0.0038 | 4 |
| Q96SI9 | STRBP | | -0.4 | 0.4 | 0.5534 | 1.5 | | 1.1 | 0.0087 | 2 |
| O95793 | STAU1 | | -0.4 | 0.2 | 0.8302 | 3.1 | | 1.3 | 0.0057 | 6 |
| Q9NVN8 | GNL3L | | -0.4 | 0.3 | 0.6774 | 2.7 | | 2.1 | 0.0006 | 2 |
| P56545 | CTBP2 | | -0.4 | 0.2 | 0.7530 | 3.2 | | 1.8 | 0.0016 | 2 |
| P52565 | ARHGDIA | | -0.5 | 0.1 | 0.8465 | -0.2 | | 0.2 | 0.8235 | 2 |
| Q92997 | DVL3 | | -0.5 | 0.5 | 0.3772 | 4.8 | | 3.4 | 0.0000 | 2 |
| Q9Y2U8 | LEMD3 | | -0.5 | 0.8 | 0.2171 | 2.7 | | 2.3 | 0.0003 | 2 |
| P26373 | RPL13 | | -0.5 | 0.1 | 0.8819 | -3.3 | | 0.6 | 0.0609 | 3 |
| Q9H2U1 | DHX36 | | -0.5 | 0.2 | 0.8196 | 2.4 | | 1.3 | 0.0057 | 2 |
| Q8N196 | SIX5 | | -0.5 | 0.5 | 0.3815 | 5.3 | | 3.8 | 0.0000 | 3 |
| P29084 | GTF2E2 | | -0.5 | 0.3 | 0.6470 | 3.8 | | 2.0 | 0.0012 | 3 |
| P50402 | EMD | | -0.5 | 0.3 | 0.6487 | 1.0 | | 0.5 | 0.1318 | 2 |
| P43268 | ETV4 | | -0.5 | 0.2 | 0.8176 | 6.8 | | 3.3 | 0.0000 | 2 |
| O96028 | WHSC1 | | -0.5 | 0.1 | 0.8378 | 3.0 | | 2.9 | 0.0000 | 4 |
| O00170 | AIP | | -0.5 | 0.7 | 0.2806 | 3.9 | | 4.1 | 0.0000 | 2 |
| P08670 | VIM | | -0.5 | 0.1 | 0.8532 | 0.3 | | 0.1 | 0.9385 | 5 |
| Q03933 | HSF2 | | -0.5 | 0.3 | 0.6547 | 3.3 | | 1.6 | 0.0027 | 2 |
| Q8N3X1 | FNBP4 | | -0.5 | 0.3 | 0.6983 | 3.1 | | 1.5 | 0.0040 | 2 |
| Q86V81 | ALYREF | | -0.5 | 0.6 | 0.3113 | 0.7 | | 1.0 | 0.0121 | 3 |
| P45974 | USP5 | | -0.5 | 0.3 | 0.5987 | 3.1 | | 2.3 | 0.0002 | 3 |
| P41182 | BCL6 | | -0.5 | 0.7 | 0.2628 | 2.2 | | 2.0 | 0.0012 | 2 |
| P23760 | PAX3 | | -0.5 | 0.7 | 0.2488 | 4.3 | | 3.7 | 0.0000 | 2 |
| P78332 | RBM6 | | -0.6 | 0.6 | 0.3417 | 3.2 | | 2.9 | 0.0000 | 3 |
| Q9HBU1 | BARX1 | | -0.6 | 0.3 | 0.6600 | 5.2 | | 3.8 | 0.0000 | 3 |
| P35579 | MYH9 | | -0.6 | 0.2 | 0.7457 | 2.8 | | 1.2 | 0.0070 | 3 |
| Q14587 | ZNF268 | | -0.6 | 0.1 | 0.8794 | 3.3 | | 0.9 | 0.0185 | 5 |
| Q9NS91 | RAD18 | | -0.6 | 1.7 | 0.0410 | 2.9 | | 3.7 | 0.0000 | 4 |
| P62318 | SNRPD3 | | -0.6 | 0.5 | 0.4239 | 1.1 | | 1.0 | 0.0124 | 2 |
| O95817 | BAG3 | | -0.6 | 0.3 | 0.6303 | 3.5 | | 2.3 | 0.0003 | 5 |
| P48634 | PRRC2A | | -0.6 | 0.2 | 0.8243 | 5.4 | | 3.9 | 0.0000 | 13 |
| Q14186 | TFDP1 | | -0.6 | 0.3 | 0.6427 | 3.9 | | 2.2 | 0.0006 | 4 |
| Q12873 | CHD3 | | -0.6 | 0.5 | 0.3818 | 3.7 | | 3.0 | 0.0000 | 5 |
| Q9UGU0 | TCF20 | | -0.6 | 0.4 | 0.4467 | 4.3 | | 3.0 | 0.0000 | 6 |
| P35869 | AHR | | -0.6 | 0.4 | 0.5196 | 6.4 | | 3.1 | 0.0000 | 8 |
| Q13185 | CBX3 | | -0.7 | 0.4 | 0.4666 | 3.7 | | 3.0 | 0.0000 | 4 |
| O43237 | DYNC1LI2 | | -0.7 | 0.6 | 0.3362 | 2.7 | | 2.0 | 0.0009 | 4 |
| Q9NRN7 | AASDHPPT | | -0.7 | 0.8 | 0.2202 | 3.7 | | 2.8 | 0.0000 | 3 |
| P56177;P56179 | DLX1;DLX6 | | -0.7 | 0.5 | 0.3824 | 5.6 | | 2.9 | 0.0000 | 3 |
| P22670 | RFX1 | | -0.7 | 0.4 | 0.5087 | 4.3 | | 2.7 | 0.0000 | 6 |
| Q7Z6Z7 | HUWE1 | | -0.7 | 0.7 | 0.2800 | 2.3 | | 2.2 | 0.0006 | 4 |
| O75400 | PRPF40A | | -0.7 | 0.2 | 0.7895 | 5.1 | | 3.3 | 0.0000 | 6 |
| Q96DF8 | DGCR14 | | -0.7 | 0.4 | 0.5495 | 5.2 | | 3.0 | 0.0000 | 7 |
| P51858 | HDGF | | -0.7 | 0.7 | 0.2379 | 1.6 | | 1.4 | 0.0043 | 4 |
| Q5HY92 | FIGN | | -0.7 | 0.4 | 0.5530 | 1.4 | | 1.6 | 0.0027 | 2 |
| Q9H0D6 | XRN2 | | -0.7 | 0.4 | 0.5202 | 4.1 | | 2.3 | 0.0005 | 4 |
| P61978 | HNRNPK | | -0.7 | 0.5 | 0.3736 | 2.6 | | 1.9 | 0.0013 | 8 |
| Q9P2J5 | LARS | | -0.7 | 0.2 | 0.7199 | 0.0 | | 0.0 | 0.9923 | 4 |
| Q8IU81 | IRF2BP1 | | -0.7 | 1.1 | 0.1113 | 2.3 | | 2.3 | 0.0005 | 6 |
| Q9H0E9 | BRD8 | | -0.7 | 0.4 | 0.4667 | 3.3 | | 1.9 | 0.0012 | 3 |
| Q14241 | TCEB3 | | -0.7 | 0.9 | 0.1569 | 0.8 | | 0.7 | 0.0468 | 8 |
| Q6P2Q9 | PRPF8 | | -0.8 | 0.6 | 0.2963 | 3.4 | | 2.1 | 0.0006 | 7 |
| Q8IX90 | SKA3 | | -0.8 | 0.5 | 0.3529 | 3.0 | | 4.5 | 0.0000 | 4 |
| Q14527 | HLTF | | -0.8 | 0.7 | 0.2768 | 2.3 | | 2.6 | 0.0000 | 3 |
| Q9UKY1 | ZHX1 | | -0.8 | 0.6 | 0.3335 | 4.1 | | 4.1 | 0.0000 | 3 |
| Q9UMN6 | KMT2B | | -0.8 | 0.4 | 0.5557 | 1.8 | | 0.7 | 0.0442 | 4 |
| P07814 | EPRS | | -0.8 | 0.4 | 0.5041 | 2.1 | | 1.6 | 0.0037 | 4 |
| Q9UNH6 | SNX7 | | -0.8 | 2.1 | 0.0207 | 1.9 | | 3.1 | 0.0000 | 2 |
| Q6ZRS2 | SRCAP | | -0.8 | 0.6 | 0.3097 | 3.9 | | 2.2 | 0.0007 | 5 |
| Q96CB8 | INTS12 | | -0.8 | 0.6 | 0.3110 | 2.3 | | 1.3 | 0.0054 | 2 |
| Q96G74 | OTUD5 | | -0.8 | 0.8 | 0.1864 | 5.1 | | 2.7 | 0.0000 | 4 |
| Q9P1Y6 | PHRF1 | | -0.8 | 0.6 | 0.3117 | 3.7 | | 3.0 | 0.0000 | 2 |
| O75461 | E2F6 | | -0.9 | 0.5 | 0.3540 | 3.9 | | 3.6 | 0.0000 | 4 |
| Q9NVM6 | DNAJC17 | | -0.9 | 1.1 | 0.1251 | 3.2 | | 3.1 | 0.0000 | 3 |
| P32780 | GTF2H1 | | -0.9 | 0.5 | 0.4279 | 1.1 | | 0.4 | 0.1769 | 3 |
| O43390 | HNRNPR | | -0.9 | 0.2 | 0.7630 | 2.2 | | 0.6 | 0.0577 | 6 |
| P62851 | RPS25 | | -0.9 | 0.4 | 0.5400 | 0.1 | | 0.1 | 0.9155 | 2 |
| Q15036 | SNX17 | | -0.9 | 0.7 | 0.2681 | 6.5 | | 3.6 | 0.0000 | 8 |
| Q13619 | CUL4A | | -0.9 | 0.9 | 0.1799 | 3.7 | | 3.3 | 0.0000 | 4 |
| A5YKK6 | CNOT1 | | -0.9 | 0.6 | 0.2926 | 1.9 | | 1.3 | 0.0064 | 2 |
| Q9UNZ2 | NSFL1C | | -0.9 | 0.6 | 0.3389 | 0.6 | | 0.6 | 0.0566 | 2 |
| Q9H0A0 | NAT10 | | -0.9 | 0.4 | 0.5244 | 0.0 | | 0.0 | 0.9980 | 3 |
| P28070 | PSMB4 | | -0.9 | 0.7 | 0.2691 | 4.1 | | 2.1 | 0.0006 | 3 |
| Q9UI26 | IPO11 | | -0.9 | 0.6 | 0.2829 | 3.1 | | 3.3 | 0.0000 | 3 |
| Q9BRD0 | BUD13 | | -0.9 | 1.1 | 0.1165 | 1.4 | | 1.6 | 0.0026 | 8 |
| Q7L590 | MCM10 | | -1.0 | 0.7 | 0.2710 | 1.1 | | 1.0 | 0.0140 | 2 |
| O60244 | MED14 | | -1.0 | 0.8 | 0.1996 | 4.5 | | 3.4 | 0.0000 | 8 |
| P26368 | U2AF2 | | -1.0 | 0.3 | 0.5800 | 4.5 | | 1.8 | 0.0016 | 4 |
| O43294 | TGFB1I1 | | -1.0 | 1.7 | 0.0391 | 2.5 | | 3.0 | 0.0000 | 3 |
| Q8IY67 | RAVER1 | | -1.0 | 0.3 | 0.6193 | 5.1 | | 2.3 | 0.0005 | 9 |
| Q15054 | POLD3 | | -1.0 | 0.6 | 0.3420 | 3.2 | | 1.9 | 0.0013 | 2 |
| Q5T3I0 | GPATCH4 | | -1.0 | 1.7 | 0.0372 | 0.3 | | 0.4 | 0.2554 | 5 |
| P51948 | MNAT1 | | -1.0 | 0.7 | 0.2266 | 5.6 | | 2.9 | 0.0000 | 4 |
| O15516 | CLOCK | | -1.0 | 1.2 | 0.1026 | 3.3 | | 1.8 | 0.0016 | 2 |
| Q86VM9 | ZC3H18 | | -1.1 | 0.7 | 0.2485 | 1.8 | | 1.8 | 0.0016 | 4 |
| Q9UKV3 | ACIN1 | | -1.1 | 0.5 | 0.3653 | 2.5 | | 2.5 | 0.0000 | 5 |
| P43243 | MATR3 | | -1.1 | 0.4 | 0.4632 | 3.5 | | 2.1 | 0.0006 | 4 |
| Q13547 | HDAC1 | | -1.1 | 0.6 | 0.2971 | 3.5 | | 1.6 | 0.0030 | 2 |
| Q9Y4W2 | LAS1L | | -1.1 | 0.8 | 0.2075 | 0.5 | | 0.2 | 0.6398 | 4 |
| O14646 | CHD1 | | -1.1 | 1.4 | 0.0645 | 0.7 | | 1.0 | 0.0145 | 8 |
| Q9BZZ5 | API5 | | -1.1 | 0.3 | 0.6248 | 1.4 | | 0.6 | 0.0680 | 3 |
| P55060 | CSE1L | | -1.1 | 1.0 | 0.1321 | 3.3 | | 2.7 | 0.0000 | 5 |
| O00716;Q14209 | E2F3;E2F2 | | -1.1 | 1.3 | 0.0847 | 3.1 | | 2.5 | 0.0000 | 2 |
| O75717 | WDHD1 | | -1.1 | 0.6 | 0.3115 | 2.2 | | 1.4 | 0.0045 | 5 |
| P55265 | ADAR | | -1.1 | 1.1 | 0.1115 | 0.8 | | 1.0 | 0.0138 | 9 |
| P08240 | SRPR | | -1.1 | 1.7 | 0.0422 | 1.6 | | 1.5 | 0.0038 | 4 |
| Q96GM5 | SMARCD1 | | -1.1 | 0.6 | 0.2976 | 4.4 | | 3.5 | 0.0000 | 4 |
| Q7Z6E9 | RBBP6 | | -1.2 | 1.3 | 0.0721 | 1.2 | | 0.9 | 0.0174 | 6 |
| O75691 | UTP20 | | -1.2 | 0.5 | 0.4355 | 0.3 | | 0.1 | 0.9157 | 3 |
| Q9ULJ7 | ANKRD50 | | -1.3 | 0.8 | 0.1939 | 3.5 | | 2.4 | 0.0000 | 5 |
| Q8NEZ2 | VPS37A | | -1.3 | 1.3 | 0.0831 | 5.4 | | 4.0 | 0.0000 | 4 |
| Q5UIP0 | RIF1 | | -1.3 | 1.0 | 0.1477 | 1.7 | | 1.1 | 0.0089 | 2 |
| Q8IVH2 | FOXP4 | | -1.4 | 1.7 | 0.0378 | 2.8 | | 3.6 | 0.0000 | 4 |
| Q1ED39 | KNOP1 | | -1.4 | 1.1 | 0.1118 | -2.3 | | 1.1 | 0.0082 | 2 |
| Q9BVJ6 | UTP14A | | -1.4 | 1.7 | 0.0421 | 0.6 | | 2.8 | 0.0000 | 14 |
| Q9Y6G5 | COMMD10 | | -1.4 | 1.0 | 0.1459 | 5.8 | | 3.8 | 0.0000 | 6 |
| Q96JP5 | ZFP91 | | -1.4 | 2.2 | 0.0140 | 2.0 | | 3.3 | 0.0000 | 6 |
| Q92541 | RTF1 | | -1.4 | 1.6 | 0.0486 | 1.1 | | 1.5 | 0.0040 | 7 |
| Q03701 | CEBPZ | | -1.4 | 1.3 | 0.0762 | -2.1 | | 2.9 | 0.0000 | 3 |
| P98170 | XIAP | | -1.5 | 1.8 | 0.0331 | 1.2 | | 2.0 | 0.0008 | 2 |
| P40425 | PBX2 | | -1.5 | 1.1 | 0.1120 | 4.3 | | 3.0 | 0.0000 | 5 |
| Q96N11 | C7orf26 | | -1.5 | 2.1 | 0.0212 | 3.2 | | 2.0 | 0.0012 | 3 |
| Q96PU8 | QKI | | -1.5 | 0.7 | 0.2653 | 2.4 | | 1.2 | 0.0066 | 3 |
| P09429 | HMGB1 | | -1.5 | 0.7 | 0.2769 | 0.0 | | 0.0 | 0.9898 | 2 |
| Q06265 | EXOSC9 | | -1.5 | 2.1 | 0.0186 | 0.8 | | 1.4 | 0.0045 | 3 |
| Q9ULJ3 | ZBTB21 | | -1.6 | 1.2 | 0.0906 | 3.5 | | 2.8 | 0.0000 | 4 |
| Q8N7H5 | PAF1 | | -1.6 | 0.8 | 0.1944 | 1.8 | | 1.2 | 0.0072 | 10 |
| P78316 | NOP14 | | -1.6 | 0.9 | 0.1624 | 1.4 | | 1.4 | 0.0043 | 3 |
| O15355 | PPM1G | | -1.6 | 1.1 | 0.1175 | 1.3 | | 1.8 | 0.0021 | 4 |
| Q8N2M8 | CLASRP | | -1.6 | 1.3 | 0.0751 | 1.2 | | 0.9 | 0.0187 | 3 |
| P09651;A0A2R8Y4L2;Q32P51 | HNRNPA1;HNRNPA1L2 | | -1.6 | 1.6 | 0.0431 | 0.9 | | 0.9 | 0.0216 | 4 |
| Q8IY81 | FTSJ3 | | -1.7 | 1.8 | 0.0352 | -0.4 | | 0.4 | 0.2904 | 10 |
| Q8IW35 | CEP97 | | -1.7 | 0.8 | 0.2162 | 3.0 | | 2.6 | 0.0000 | 5 |
| O00193 | SMAP | | -1.8 | 0.4 | 0.4989 | 2.3 | | 1.5 | 0.0038 | 3 |
| Q9BU76 | MMTAG2 | | -1.8 | 1.7 | 0.0385 | 1.9 | | 1.9 | 0.0013 | 4 |
| Q9NQZ2 | UTP3 | | -1.9 | 1.0 | 0.1383 | 0.5 | | 0.2 | 0.7352 | 2 |
| Q9UHB7 | AFF4 | | -2.0 | 1.3 | 0.0780 | 0.4 | | 0.2 | 0.8151 | 3 |
| O00505 | KPNA3 | | -2.0 | 1.5 | 0.0607 | 0.3 | | 0.4 | 0.3081 | 3 |
| P22626 | HNRNPA2B1 | | -2.1 | 2.2 | 0.0146 | 0.7 | | 0.7 | 0.0522 | 4 |
| Q13428 | TCOF1 | | -2.1 | 2.1 | 0.0209 | -1.3 | | 1.3 | 0.0055 | 4 |
| Q9UIG0 | BAZ1B | | -2.2 | 2.3 | 0.0138 | 0.7 | | 1.2 | 0.0066 | 6 |
| Q53F19 | C17orf85 | | -2.2 | 1.3 | 0.0847 | -1.9 | | 2.0 | 0.0012 | 4 |
| Q99459 | CDC5L | | -2.2 | 2.1 | 0.0206 | 3.9 | | 2.8 | 0.0000 | 10 |
| P06748 | NPM1 | | -2.2 | 0.7 | 0.2425 | -0.7 | | 0.4 | 0.2467 | 4 |
| P52943 | CRIP2 | | -2.2 | 3.4 | 0.0056 | -0.6 | | 0.1 | 0.9296 | 2 |
| Q8NI36 | WDR36 | | -2.4 | 0.6 | 0.3146 | 2.5 | | 1.8 | 0.0021 | 6 |
| Q9Y2H2 | INPP5F | | -2.4 | 3.0 | 0.0063 | 1.4 | | 0.7 | 0.0465 | 4 |
| P36873;P62136;P62140 | PPP1CC;PPP1CA;PPP1CB | | -2.4 | 1.7 | 0.0380 | 0.7 | | 0.5 | 0.1343 | 3 |
| P46013 | MKI67 | | -2.4 | 2.4 | 0.0141 | 0.1 | | 0.1 | 0.9502 | 17 |
| P14868 | DARS | | -2.5 | 2.1 | 0.0210 | -1.4 | | 1.6 | 0.0033 | 2 |
| Q8NEF9 | SRFBP1 | | -2.6 | 1.3 | 0.0836 | 0.3 | | 0.5 | 0.1257 | 2 |
| O60506 | SYNCRIP | | -2.8 | 1.5 | 0.0558 | 2.6 | | 1.8 | 0.0016 | 5 |
| Q9H6R4 | NOL6 | | -2.8 | 2.3 | 0.0140 | 0.9 | | 1.2 | 0.0066 | 10 |
| P46087 | NOP2 | | -2.9 | 3.4 | 0.0059 | -3.6 | | 3.2 | 0.0000 | 3 |
| Q6PD62 | CTR9 | | -3.0 | 1.8 | 0.0330 | 2.5 | | 1.8 | 0.0016 | 7 |
| Q9NYF8 | BCLAF1 | | -3.0 | 1.8 | 0.0370 | 0.0 | | 0.0 | 0.9977 | 7 |
| O95373 | IPO7 | | -3.1 | 2.2 | 0.0156 | -0.9 | | 0.5 | 0.0924 | 12 |
| Q9BQG0 | MYBBP1A | | -3.2 | 1.8 | 0.0318 | -2.0 | | 1.1 | 0.0082 | 7 |
| P55209 | NAP1L1 | | -3.7 | 1.1 | 0.1189 | -3.3 | | 1.8 | 0.0016 | 4 |
| Q14692 | BMS1 | | -4.1 | 1.5 | 0.0574 | -2.1 | | 1.3 | 0.0061 | 10 |
| Q9NVP1 | DDX18 | | -4.2 | 2.8 | 0.0057 | -3.3 | | 1.9 | 0.0013 | 5 |
| P51991 | HNRNPA3 | | -4.2 | 1.6 | 0.0474 | 1.2 | | 0.5 | 0.1029 | 3 |
| Q96EB6 | SIRT1 | | -4.3 | 2.1 | 0.0185 | -4.5 | | 2.7 | 0.0000 | 5 |
| Q9NXF1 | TEX10 | | -4.5 | 2.7 | 0.0064 | -1.2 | | 1.4 | 0.0052 | 6 |
| P19338 | NCL | | -4.6 | 1.4 | 0.0669 | -1.8 | | 2.7 | 0.0000 | 7 |
| Q8WTT2 | NOC3L | | -5.3 | 2.1 | 0.0215 | -5.1 | | 3.2 | 0.0000 | 6 |
| Q5T8P6 | RBM26 | | -5.4 | 0.7 | 0.2723 | -0.7 | | 0.1 | 0.9390 | 3 |

**Table Legend S1.** This table presents a ranked list of high-confidence candidate hSOX10-interacting proteins identified using mT proximity-dependent biotinylation followed by MS in A375 human melanoma cells. Two distinct hSOX10 fusion constructs were employed: mT-hSOX10 and hSOX10-mT, each compared to NLS-mT controls. For each construct, log_2_ fold change (Log_2_ FC), -log_10_(p-value), and adjusted p-value (q-value) were calculated relative to the NLS-mT control. In Log_2_ FC columns, light red indicates significant enrichment compared to control, while light blue indicates significant underrepresentation. In adjusted p-value columns, yellow marks statistical significance, while unshaded cells are not significant. In the gene names column, green highlights indicate previously reported hSOX10 interactors, while orange highlights the hSOX10 bait protein
